# Take it or leaf it: bees learn leaf shape as a cue when flower color is easily learned

**DOI:** 10.1101/2025.06.12.659384

**Authors:** Anthony Moth T. Castagna, Jenny K. Burrow, Ciara G. Stewart, Avery L. Russell

## Abstract

Over a century of research has demonstrated that pollinators, such as bees, associatively learn diverse flower cues, including tactile, visual, and olfactory cues, to find food rewards. However, floral cues are not always reliable, as flowers of different plant species often differ in terms of the qualities of their food rewards, even when flower types resemble each other. At the same time, some non-floral traits, such as leaf shape, differ reliably among plant species and might be associatively learned by bees to improve foraging success. In this laboratory study, we tested whether generalist bees (*Bombus impatiens*) can (1) associatively learn differences in leaf shape to discriminate rewarding from unrewarding flowers and (2) rely more on differences in leaf shape when a flower color cue is harder to discriminate. We therefore assigned bees to either of two treatments: in one treatment, rewarding and unrewarding artificial targets (‘flowers’) differed greatly in petal color, and in the other treatment, they differed little; each treatment’s targets differed in leaf shape in the same way. As expected, bees learned significantly faster when flower petal colors were more dissimilar and thus relatively easier to discriminate. These bees also learned and recalled the correct combination of petal color and leaf shape. Yet when petal colors differed relatively little, bees had a much harder time learning petal color and did not show evidence of having remembered leaf shape. Our results demonstrate that leaf shape is a cue that foraging bees can learn to associate with a pollen food reward. However, leaf shape may be learned secondarily to, or only in combination with floral cues (such as petal color). We discuss evidence of compound learning and overshadowing and implications of our results for pollinator-mediated selection on non-floral plant traits.

## INTRODUCTION

A central question in animal behavior is how animals decide which cues to use to find food (Kamil & Roitblat 1985; Stephens 2008). Generalist animals frequently encounter situations of uncertainty while foraging, such as when a cue cannot reliably be associated with food (McLinn & Stephens 2006; Page & Jones 2016). Generalist pollinators frequently learn to associate tactile, visual, olfactory, and other cues produced by flowers with floral food rewards (e.g., nectar and pollen) (Chittka & Raine 2006; Whitney et al. 2009; Riffell 2011; Clarke et al. 2013; Clarke et al. 2017; Harrap et al. 2017). However, pollinators also often encounter uncertainty while foraging on flowers. For instance, many plant species deceive their pollinators by offering floral cues that do not honestly indicate reward presence, quality, or amount (Ashman 2009; Lichtenberg et al. 2020; van der Kooi et al. 2023). Furthermore, co-flowering plant species may share similar floral cues but offer food rewards that differ in quality (e.g., Internicola et al. 2007), making those shared cues unreliable. For instance, up to 5% of plant species offer no rewards at all and instead attract pollinators by mimicking the floral displays of rewarding co-flowering plant species (Gigord et al. 2002; Schiestl 2005; Lichtenberg et al. 2020, Figure 1). Consequently, particularly when floral cues are unreliable, we expect pollinators should learn cues from other parts of the plant that can be reliably associated with floral rewards. However, whether and when non-floral plant cues alone or in combination with flower cues are used to discriminate among flower types by foraging pollinators has scarcely been explored (see Cepero et al. 2015).

**Figure 1.**
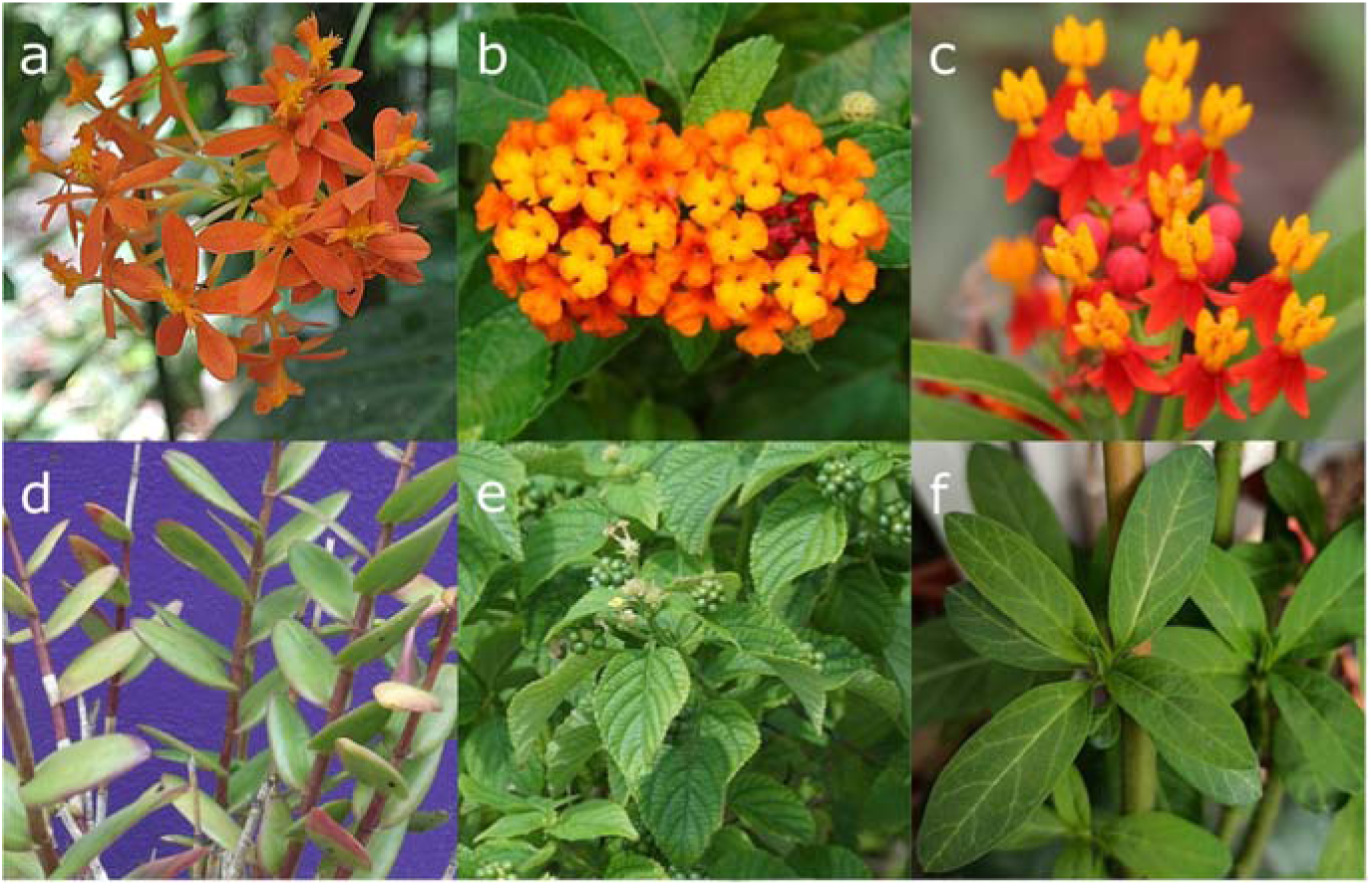
Three plant species whose flowers resemble each other and are thought to be involved in **(a, b)** Batesian and **(c)** Müllerian mimicry, but whose leaves vary in shape. Flower and leaves of (**a, d)** an unrewarding mimic, *Epidendrum ibaguense*, **(b, e)** the rewarding model, *Lantana camara*, and the **(c, f)** co-occurring rewarding Müllerian mimic, *Asclepias curassavica*. Photographs: **(a)** Dick Culbert, **(b)** Vineeth Vengolis, **(c)** Fan Wen, **(d)** Yercaud Elango, **(e)** Vengolis, **(f)** Juan Carlos Fonseca. Licensing: **(a)** CC BY 2.0, **(b)** CC BY-SA 4.0, **(c)** CC BY-SA 4.0, **(d)** CC BY-SA 4.0, **(e)** CC BY-SA 4.0, **(f)** CC BY-SA 4.0.

Over the past century, the vast majority of research conducted on how pollinators locate floral food rewards has focused solely on the role of floral cues (see Chittka & Thomson 2005; Giurfa 2007). Flower color, pattern, shape, and scents can differ significantly between species of flowering plants, enabling pollinators to learn and develop strong, stable preferences associated with these floral cues (Essenberg et al. 2015; Hempel de Ibarra & Somanathan 2019; Nicholls et al. 2022). However, non-floral traits such as leaf shape also often differ between animal-pollinated flowering plant species. For instance, leaves of a given plant species may be strap-like or obovate, pinnately or palmately compound (Kidner & Umbreen 2010; Fowler 2016). Notably, leaf shape can differ between plant species engaged in floral Batesian mimicry – when rewarding flowers of one plant species are mimicked by the unrewarding flowers of another plant species – and floral Müllerian mimicry – when flowers of multiple co-flowering plant species are rewarding and resemble one another (Figure 1) (Bierzychudek 1981). That leaf shape differs between plant species, especially when floral cues may be unreliable, raises the possibility that pollinators can learn to associate differences in leaf shape with floral rewards.

Leaf shape is often learned by animals in other contexts. For instance, various butterfly and fly taxa learn to use leaf shape to locate host plants for oviposition (Wiklund 1984; Allard & Papaj 1996; Degen & Städler 1996; Dell’Aglio et al. 2016). Similarly, braconid wasp parasitoids readily learn leaf shape to locate their caterpillar hosts (Wäckers & Lewis 1999). In addition, honey bees, bumble bees, and monarch butterflies can learn to associate flower shape with a nectar reward (Gould 1985; Zhang et al. 1995; Cepero et al. 2015), suggesting that pollinators might also be able to learn the shape of leaves while foraging. Even still, other floral cues are learned preferentially over flower shape (Rusch et al. 2017). Additionally, relative to other floral cues, flower shape is more difficult to learn for even generalist bees, a model system for the study of pollinator learning and memory (Menzel 1967; Lehrer et al. 1985; Lehrer 1990; Lehrer 1993). Assuming similar challenges when learning leaf shape, we expect that generalist bees may attend little to leaf shape when floral cues are easy to learn and instead rely primarily on floral cues. At the same time, because compound cues have enhanced saliency and facilitate associative learning (Telles et al. 2017), when floral cues are hard to learn, we expect that bees should attend to leaf shape and that leaf shape should be learned in compound with floral cues.

In this laboratory study, we assessed whether and within what context a generalist bee can learn to associate a floral food reward (pollen) with leaf shape. Given that shape is considered a difficult cue to learn, we hypothesized that bumble bees (*Bombus impatiens*) would learn leaf shape, but only when flower color was relatively difficult to learn (e.g., when the saliency of leaf and flower color cues are more similar). We tested this hypothesis via an associative learning assay in which bees were trained to associate a pollen reward with a given combination of leaf shape and petal color (each combination is a ‘target’). In one treatment, petals of different target types differed greatly in bee color space, which we predicted would be easier for bees to learn; in the other treatment, petals of different target types differed little in bee color space, which we predicted would be harder for bees to learn. Leaf shape differed among target types in the same way across treatments. Finally, we predicted that to reduce uncertainty when the petal color learning task was more difficult, bees would be more likely to learn specific combinations of leaf shape and petal color associated with the floral reward.

## METHODS

### Experiment outline

We conducted an experiment with two treatments, both of which involved an initial assessment of responses by flower-naïve bees to four types of pollen-rewarding artificial targets (‘flowers’ comprising combinations of petal color and leaf shape; Figure 2). In the simple learning treatment, flower-naïve bees were presented with artificial flowers that differed greatly in terms of petal color (blue versus yellow) in a bee color vision model. In the difficult learning treatment, flower-naïve bees received flowers that differed little in terms of petal color (blue versus purple). For both treatments, both sets of flowers differed in the same way in terms of leaf shape (Figure 2). Following the initial assessment of preference, these bees were given experience with only two of their four flower types in a training phase (Figure 2a; in the simple learning treatment: blue petals, short leaves vs yellow petals, long leaves; Figure 2b; in the difficult learning treatment: blue petals, short leaves vs purple petals, long leaves), in which one flower type was rewarding and the other was unrewarding, alternated with every trial. In a subsequent unrewarded test phase, the trained bees were allowed to visit any of the four flower types that they were exposed to in their initial preference assay. The initial preference assay evaluates each bee’s naïve preference for each leaf shape, petal color, and their combination. The training assay trained a bee to associate a particular petal color and leaf shape with a reward and allows us to assess whether the rate of learning differed by treatment. The test assay evaluates how bee preference for a specific petal color and/or leaf shape changed after training relative to the initial preference assay. Details of system and protocol follow.

**Figure 2.**
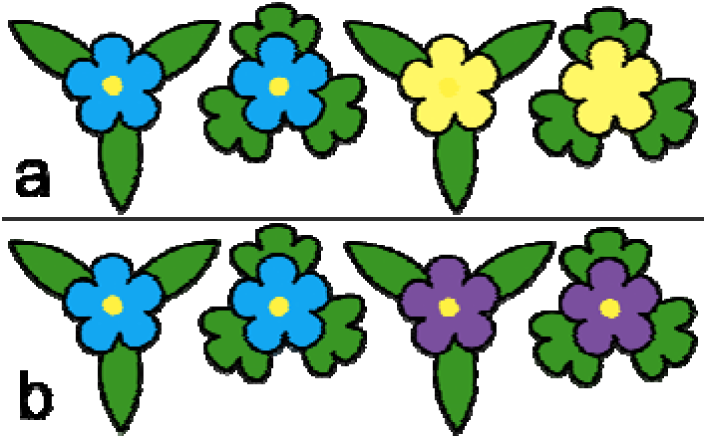
Artificial flower types used in the **(a)** simple and **(b)** difficult treatments. Treatments differed in terms of how similar the petal colors are in bee color space. Leaf shapes differed in the same way for both treatments. The artificial anther is depicted as the yellow dot in the center of all of the flowers. Flower types from left to right are, in panel **(a)** blue petal, long leaves (abbreviated as BL); blue petal, short leaves (BS); yellow petal, long leaves (YL); yellow petal, short leaves (YS); and in panel **(b)** blue petal, long leaves (BL); blue petal, short leaves (BS); purple petal, long leaves (PL); purple petal, short leaves (PS).

### Bees and housing

We used 79 workers from four captive-bred and commercially obtained (Biobest USA Inc., Romulus, MI, U.S.A.) colonies of *Bombus impatiens* bumble bees, maintained following Russell et al. (2017). To summarize, bees were permitted to forage on artificial feeders providing nectar solution (2M sucrose) and pulverized honey bee collected pollen (Plant Products Inc.) ad libitum. Bees were contained within a plywood arena (L × W × H 82 × 60 × 60 cm) painted grey on the floor and sides with a clear acrylic ceiling, lit from above and set to a 14:10 light:dark cycle.

### Artificial flowers

To precisely manipulate petal color and leaf shape, we cut plastic (Polypropylene Heavy Duty Plastic Folders, Amazon Basics) into the shape of petals and leaves and hot-glued to microcentrifuge tubes, with leaves offset two cm behind the petals. Petal shape was identical across all treatments and the two types of leaf shapes had identical total surface area (measured using ImageJ). We mounted a disposable artificial anther (a uniformly sized yellow chenille stem) to the center of each artificial flower’s corolla (‘petal’). We used chenille stems as artificial anthers following Russell & Papaj (2016), as they can hold and present precise amounts of pollen to foraging bees. To prevent bees from visually or olfactorily assessing pollen presence or absence, pollen and artificial anthers were a similar hue of yellow and unrewarding anthers were pollen scented following Muth et al. (2016).

To create treatments that differed in the difficulty of the color learning task, we selected two pairs of petal colors that were closer and further apart in bee color space, respectively (Figure 2a, b; Figure 3a). We closely matched the color of the artificial plastic leaves to live leaves collected from the Missouri State University campus (*Amelanchier sp.*, *Philodendron sp.*, *Tulipa sp.*) (Figure 3b).

**Figure 3.**
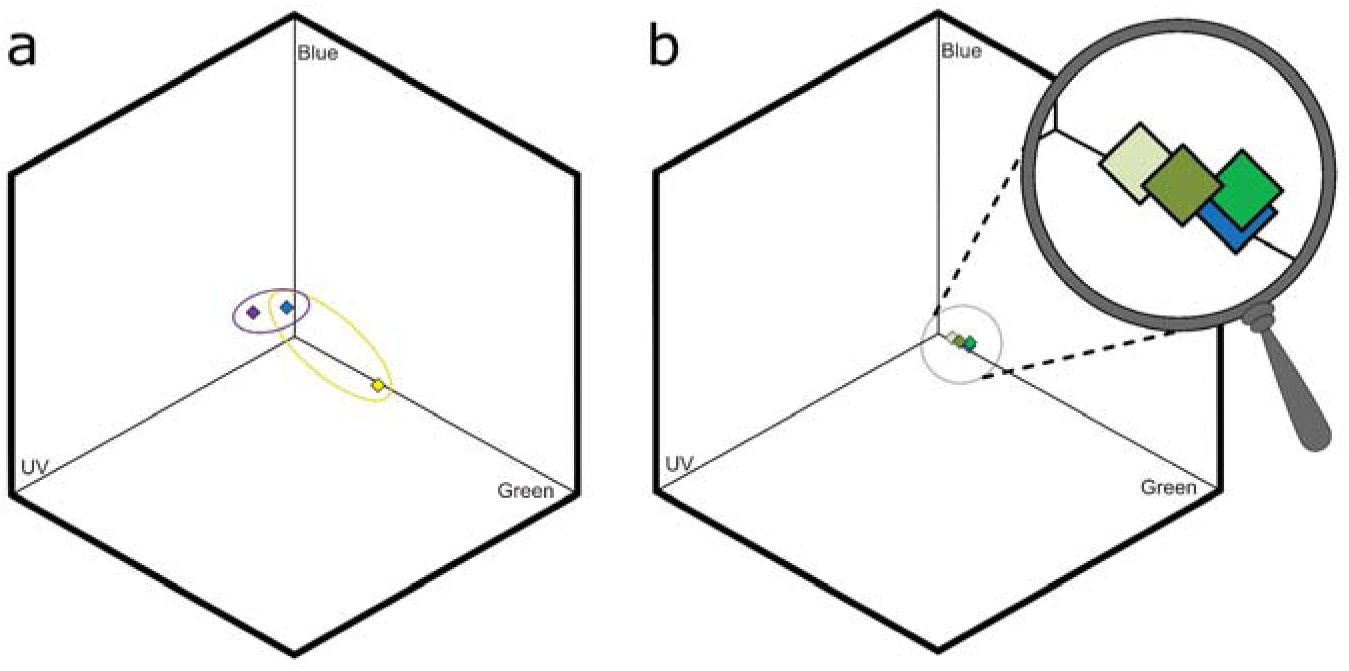
The loci in *Bombus impatiens* color space of **(a)** artificial blue (blue diamond), purple (purple diamond), and yellow (yellow diamond) petals and **(b)** live (*Amelanchier sp.*, *Philodendron sp.*, *Tulipa sp.*) and artificial leaves against the test arena background. Petal colors used in the simple and difficult treatments are encircled in yellow and purple, respectively. Blue and purple petals differed by 0.10 color units and from the background by 0.10 and 0.15 color units, respectively. Blue and yellow petals differed by 0.41 color units and the yellow differed from the background by 0.32 units. Artificial leaves differed from live leaves by 0.037 ± 0.019 (mean ± SE) color units and the green plastic differed from the background by 0.11 color units.

### Spectral analyses

Following Russell et al. (2021), we measured the color reflectance of the artificial petals and leaves (Amazon Basics) and live leaves with a UV-VIS spectrometer (USB2000, Ocean Optics, University, FL, USA), tungsten-deuterium light source (DH2000-BAL, Ocean Optics, University, FL, USA), and a fluoropolymer white standard (WS-1-SL Spectralon, Ocean Optics, University, FL, USA). We used an RPH reflectance probe (Ocean Optics, University, FL, USA), shielded from extraneous light, and held at a constant height and 45° angle above the samples.

Reflectance measurements were all taken using a 5 ms integration time averaging 500 ms in the same session. We measured irradiance within the flight arena at the center of the flower array using a 50 ms integration time and 50 ms averaging, CC-3-UV-S cosine-corrected (180 degrees) irradiance probe (Ocean Optics, University, FL, USA), Q400-7-SR UV/VIS optical fiber (Ocean Optics, University, FL, USA), and a tungsten-deuterium calibration light source (DH-3P-CAL, Ocean Optics, University, FL, USA). Each reflectance spectrum consisted of the mean of 10 measurements, taken from different petals, leaves, or parts of the arena wall background.

To characterize what bees perceived, we used our reflectance and irradiance measurements to plot artificial petals (Figure 3a) and leaves (Figure 3b) within a color space (e.g., the color hexagon) for *Bombus impatiens* following Russell et al. (2017). We crafted a color space diagram using UV-blue-green trichromat receptor sensitivities following Chittka (1992) and Skorupski & Chittka (2010), then transformed to spectral sensitivity curves following Stavenga et al. (1993). We used the grey-painted test arena wall, where the flowers were displayed, as the background stimulus for the color hexagon, and we used the irradiance of the overhead arena lights to calculate the receptor excitation values.

We selected the colors for our artificial flower petals and leaves by how they differed in bee color vision. Bumble bees have difficulty discriminating colors that are close in color space and fail to discriminate colors that are less than 0.04 color units apart (Dyer & Chittka 2004). Previous research notes that the difficulty of the color learning task is a reliable way to quantify bee perceptual difficulty of a visual task (Dyer & Garcia 2014). Therefore, petal colors selected for the simple treatment (blue versus yellow, which differ by 0.41 color units) should be more easily discriminable than petal colors selected for the difficult treatment (blue versus purple, which differ by 0.10 color units). Artificial leaves were a close match to live leaves in bee color space and differed on average by 0.037 color units. Colors of artificial petals and leaves differed from the background by at least 0.10 color units.

### Experimental Protocol

Behavioral trials were conducted in a cleaned and grey-painted plywood test arena (L × W × H 82 × 60 × 60 cm) with a clear acrylic ceiling, lit from above, following Russell & Ashman (2019). To summarize, artificial flowers were spaced 7 cm apart in a Cartesian grid design attached to the arena wall opposite a marking surrounding an artificial nest entrance. We systematically alternated flower positions for each trial.

To initiate a trial, we first set up flowers vertically on the arena wall and allowed one flower-naïve worker bee (hereafter ‘naïve bee’) into the arena. Each bee visited nearly all of the flowers in its trial at least once. To ensure trials were comparable, trials were terminated after the naïve bee met our predetermined criterion, did not approach any flower for a period of 5 mins, or attempted to leave the arena via the nest entrance, whichever came first. We recorded flower-visiting behavior following Muth et al. (2016). To summarize, we recorded flower ‘visits,’ which we defined as a bee touching the scented or pollen-covered anther. When bees collected pollen from rewarding flowers (scraped from anthers by ‘scrabbling’; see Russell & Papaj 2016), the visit was recorded as ‘rewarded’. When bees visited unrewarding flowers, the visit was recorded as ‘unrewarded’. In the rare cases when bees visited, but did not collect pollen from rewarding flowers, we excluded the visit from analyses because we could not be sure whether the visit reinforced or inhibited learning. Bees never depleted rewarding flowers of pollen and rarely filled their pollen baskets during trials. After a bee completed its trials, the bee was euthanized. After each trial, we discarded the artificial anthers and thoroughly cleaned the artificial flowers with 70% ethanol to be reused following Russell & Ashman (2019), as bees leave chemical footprints when foraging (Roselino et al. 2016). At the conclusion of each trial, and before starting again, the testing arena floor was cleaned.

Each of 79 naïve bees from 4 colonies was assigned to 1 of 2 treatments. For both treatments, we first tested the initial petal color and leaf shape preference of a bee using a 4 × 4 array of 4 types of pollen-rewarding flowers and systematically alternated flower position in the grid. The 4 types of flowers (the 4 flower petal and leaf combinations) present in the array depended on the assigned treatment (see Figure 2a vs Figure 2b). Each bee was permitted to make up to 40 pollen-collecting visits in its single trial. The artificial anther of each rewarding flower was loaded with 5.5 mg of cherry pollen (*Prunus avium*, Antles Pollen Inc., Wenatchee, WA, USA). Once a trial was complete, the bee was re-captured, marked with a unique paint color using non-toxic oil-based markers (Sharpie, CA, USA), and returned to its colony.

After 24-48 hours, marked bees were individually differentially conditioned (‘trained’) to the specific rewarding leaf and petal combination within their assigned treatment (see Giurfa 2007) (Figure 2). Bees were allowed to forage individually on a 5 × 4 training assay of 20 flowers with one rewarding flower type of the respective treatment (S+) and one unrewarding flower type (S-), with flower types alternated by position. Anthers of rewarding flowers were loaded with 5.5 mg of cherry pollen and anthers of unrewarding flowers were pollen-scented. We alternated which of the two training flower types were rewarding across trials, thus creating two sub-treatments per treatment (Figure 2, 6). We permitted bees to forage until they had made 8 of their last 10 rewarding visits to the rewarding flower type, which we considered evidence of the bee having learned. Bees that did not meet this learning criterion in their first training treatment (i.e., did not visit any flower for a period of 5 mins, or made several consecutive nest entrance visits) were returned to their colony to unload pollen and subsequently given a second training trial (16 of the 38 successful bees). Of 79 bees, 41 did not meet the learning criterion even after two trainings and were subsequently euthanized. One bee that met the learning criterion did not complete a retention test.

To test whether trained bees retained the learned preference, each was given a single retention test one hour after training. Each bee was individually tested and presented with unrewarding, pollen-scented flowers in a 4 × 4 array with the same flower types as in that bee’s initial preference assay. We allowed each bee in the retention test to make up to 40 flower visits.

### Data analyses

We analyzed how rate and strength of learning was affected by treatment and how preferences for the trained rewarding petal color, leaf shape, and/or the combination changed with experience. All data were analyzed using R statistical software, using R v.4.2.2 (R Development Core Team 2022) and RStudio 2023.09.1+494 (Ushey et al. 2022).

### Do bees learn more quickly when the learning task is easier?

To determine whether bees made more visits to the rewarding flowers with successive visits (i.e., learned which flower type was rewarding) and whether the pattern differed by treatment, we analyzed only bees that met our learning criterion and used a binomial generalized linear mixed effects model (GLMM) similar to Russell & Ashman (2019). We fit a model using the glmmTMB() function from the glmmTMB package (Brooks et al. 2017), specifying type II Wald chi-square (χ^2^) tests via the Anova() function in the car package (Fox & Weisberg 2019), and checked model assumptions using the DHARMa (Hartig & Lohse 2022) and sjPlot (Lüdecke 2024) packages. We specified ‘visit type’ (rewarded or unrewarded) as the response variable and ‘treatment’ (simple or difficult) and ‘visit number’ as the explanatory variables. We included ‘visit number’ as a repeated measure, nested within ‘bee ID’, nested within ‘colony ID’ as our random effects.

### Do bees learn the leaf-petal combination regardless of the difficulty of the learning task?

To analyze differences in preference across the four flower types for each group of bees in each treatment (Figure 2, 6), before and after learning, we used a hierarchical Bayesian model (BayesPref package; Fordyce et al. 2011) for multinomial count data, following Russell et al. (2018). Detailed information regarding the advantages of this model is in Fordyce et al. (2011); Forister & Scholl (2012); and Gompert & Buerkle (2012). For each analysis, we ran MCMC for 40,000 generations and discarded the first 10,000 runs as burn-in. We confirmed even mixing for all posteriors using the ‘plot’ function. To identify significant differences among estimates of preference for each of the four flower types in each treatment, we used pairwise comparisons of posterior probabilities (noted as ‘PP’) (BayesPref package; Fordyce et al. 2011). When preference for a particular flower type is greater than preference for another flower type – or when preference for a particular flower type after experience (i.e., the retention test) is greater than the initial preference for that flower type – in more than 95% of the sampled MCMC steps, the preference estimates are considered to be significantly different (Fordyce et al. 2011). Though we use a Bayesian approach for examining pairwise differences, we can interpret posterior probability values similarly to the frequentist *P* - α (where α = 0.05). While pairwise comparisons offer complementary values for both choice A over B and B over A, we report only the smaller value to align with a frequentist interpretation. See Russell et al. (2018) for a complete explanation.

### Is associative learning mediated equivalently by petal color and leaf shape?

To determine if treatment influenced whether bees learned to associate petal color or leaf shape with the pollen reward, we first calculated the mean percentage of each bees’ visits to the trained petal color or leaf shape in the initial preference assay and retention test. We compared initial to test preference for petal color or leaf shape via GLMMs as above. The response variable was ‘percent preference’ (for the rewarded petal color or leaf shape) and ‘experiment phase’ (initial vs test) was the explanatory variable. For these models, we specified ‘bee ID’, nested within ‘colony ID’ as our random effects.

## RESULTS

### Bees learned more quickly when the learning task was easier

Bumble bees (*B. impatiens*) visiting artificial targets (‘flowers’) rapidly learned via differential conditioning whether a given flower type was associated with the pollen reward. Bees in both training treatments made more flower visits across consecutive visits to the rewarding flower type (GLMM: visit number effect: χ^2^_1_ = 5.579, *P* = 0.019; treatment effect: χ^2^_1_ = 2.472, *P* = 0.116; effect size = 0.305). However, when the petal colors of the two artificial flower types were perceptually more dissimilar (simple treatment), bees learned significantly more quickly than when the petal colors of the two flower types were perceptually more similar (difficult treatment) (Figure 4; GLMM: visit number × treatment interaction: χ^2^_1_ = 6.016, *P* = 0.015). This difference corresponds to a 41% better performance in the mean number of flower visits to reach the learning criterion (mean no. visits ± SE: trained in simple treatment: 48.9 ± 5.0, *N* = 17 bees; trained in difficult treatment: 68.9 ± 8.3, *N* = 21 bees).

**Figure 4.**
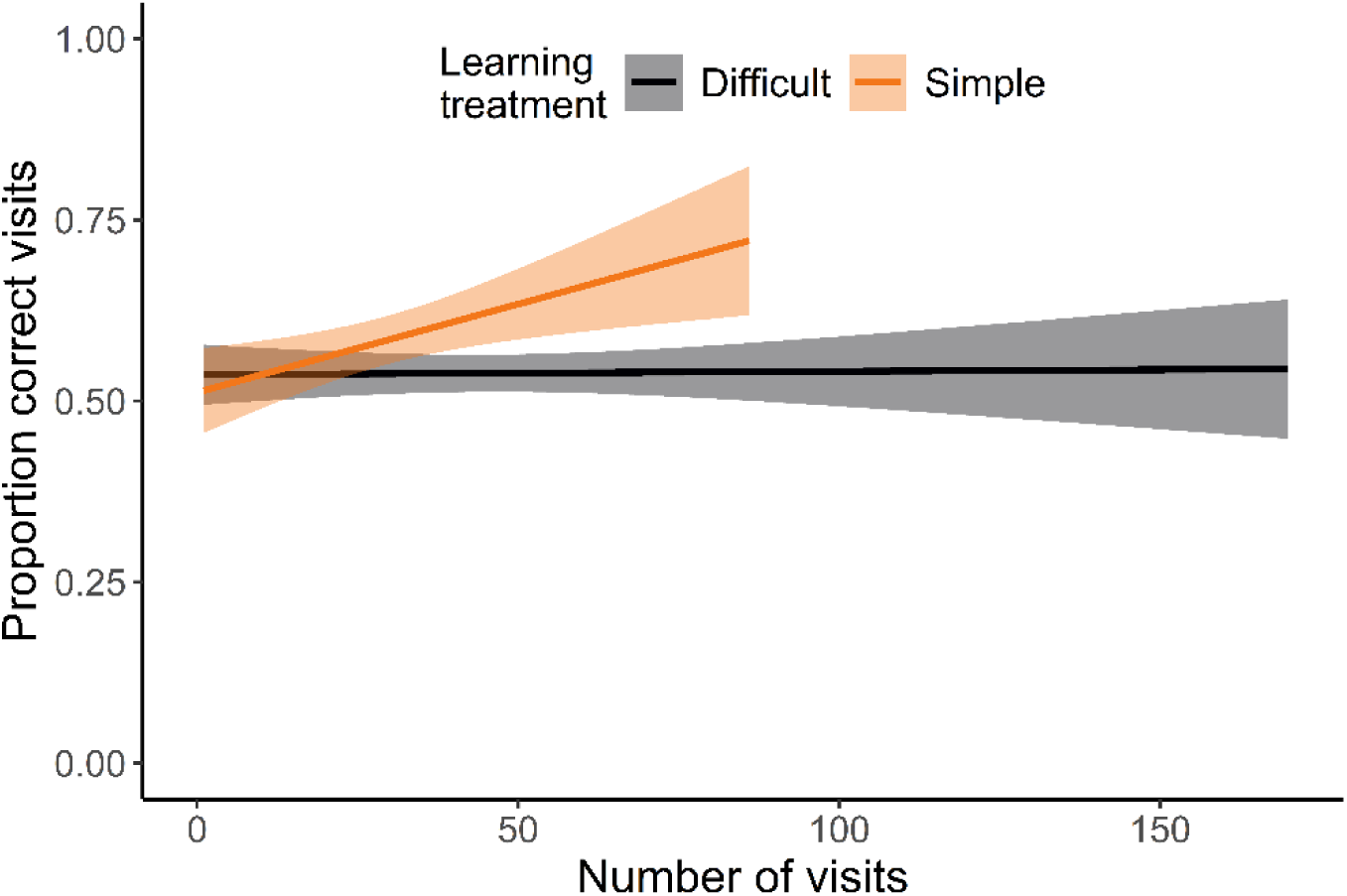
Mean proportion of correct (rewarded) visits to the rewarding flower type during the training phase. *N* = 17 and 21 bees in the simple and difficult treatment, respectively. Plotted lines indicate means and shaded regions indicate 95% confidence intervals.

### Bees learned the leaf-petal combination when the learning task was easier

Naïve bees showed no preference for any one flower type over any other flower type, regardless of treatment (Figure 5; Table 1-2). Conversely, relative to their initial preference, experienced bees in unrewarded retention tests for the simple treatment significantly preferred the flower type that had been rewarded (hereafter ‘CPCL’; flower types with ‘correct petal’ and ‘correct leaf’), avoided flower types with the incorrect petal color (‘IPCL’ or ‘IPIL’, flower types with incorrect petal color), and did not alter their preference when only leaf shape was incorrect (‘CPIL’) (Figure 5; Bayesian analysis: initial vs test preference for each flower type; Table 1; simple treatment: CPCL, *PP* = 0.007; CPIL, *PP* = 0.084; IPCL, *PP* = 0.020; IPIL, *PP* = 0.022). Experienced bees also significantly preferred flowers with the correct petal color (‘CP’) over flowers with the incorrect petal color (‘IP’), regardless of leaf shape (‘CL’ or ‘IL’) (Figure 5; Table 1; Bayesian analysis: differences in preference among flower types; CPIL vs IPIL, *PP* = 0.007; CPIL vs IPCL, *PP* = 0.013; CPCL vs IPIL, *PP* = 0.009; CPCL vs IPCL, *PP* = 0.003; CPIL vs CPCL, *PP* = 0.262; IPIL vs IPCL, *PP* = 0.470).

**Figure 5.** Initial and unrewarded retention test preference for each flower type, for bees in the **(a)** simple treatment or **(b)** difficult treatment. *N* = 17, 20 bees for simple and difficult treatments, respectively. Asterisks indicate pairwise differences for each flower type for the initial versus retention test at posterior probabilities <0.05. Dashed line at 25% indicates random expectation for an assay with four choices. In reference to the trained rewarding flower target, C = correct, I = incorrect, P = petal, L = leaf; therefore CPCL = correct petal correct leaf, CPIL = correct petal incorrect leaf, IPCL = incorrect petal correct leaf, IPIL = incorrect petal incorrect leaf.

**Table 1.** Differences in posterior probabilities via a Bayesian analysis when assigned to the simple treatment.

|  | Initial preference | Test preference | Initial vs Test |
| --- | --- | --- | --- |
| CPIL vs CPCL | 0.400 | 0.262 |  |
| CPIL vs IPIL | 0.345 | 0.007 |  |
| CPIL vs IPCL | 0.287 | 0.013 |  |
| CPCL vs IPIL | 0.264 | 0.009 |  |
| CPCL vs IPCL | 0.208 | 0.003 |  |
| IPIL vs IPCL | 0.432 | 0.470 |  |
| CPIL |  |  | 0.084 |
| CPCL |  |  | 0.007 |
| IPIL |  |  | 0.022 |
| IPCL |  |  | 0.020 |

In contrast, bees in the difficult treatment did not alter their preference for any flower type with experience (Figure 5; Bayesian analysis: initial vs test preference for each flower type; Table 2; difficult treatment: CPCL, *PP* = 0.227; CPIL, *PP* = 0.181; IPCL, *PP* = 0.277; IPIL, *PP* = 0.132). For preferences divided among sub-treatments for both treatments, see supplementary material Tables S1-S4 and Figure S1.

**Table 2.** Differences in posterior probabilities via a Bayesian analysis when assigned to the difficult treatment.

|  | Initial preference | Test preference | Initial vs Test |
| --- | --- | --- | --- |
| CPIL vs CPCL | 0.424 | 0.364 |  |
| CPIL vs IPIL | 0.353 | 0.079 |  |
| CPIL vs IPCL | 0.292 | 0.214 |  |
| CPCL vs IPIL | 0.299 | 0.150 |  |
| CPCL vs IPCL | 0.232 | 0.338 |  |
| IPIL vs IPCL | 0.422 | 0.257 |  |
| CPIL |  |  | 0.181 |
| CPCL |  |  | 0.227 |
| IPIL |  |  | 0.132 |
| IPCL |  |  | 0.277 |

### Associative learning was primarily mediated by petal color

Learned preferences for petal color, but not leaf shape, were retained for at least 1 hour by bees assigned to the simple treatment. Relative to their initial petal color preference and independent of leaf shape, these experienced bees significantly preferred the correct petal color (Figure 6a; GLMM: χ^2^_1_ = 11.743, *P* = 0.0007, effect size = 1.18). Conversely, experienced bees in the difficult treatment did not similarly retain their petal color preference (Figure 6a; GLMM: χ^2^_1_ = 1.095, *P* = 0.295, effect size = 0.349). However, in one of the two sub-treatments there was a trend for experienced bees to recall the correct petal color (GLMM: trained to PL flowers: χ^2^_1_ = 3.785, *P* = 0.052, effect size = 0.973; trained to BS flowers: χ^2^_1_ = 0.005, P = 0.945, effect size = 0.031). In contrast, regardless of treatment or sub-treatment, experienced bees did not retain their preference for leaf shape independent of flower color (Figure 6b; GLMMs: simple treatment: χ^2^_1_ = 1.803, *P* = 0.180; difficult treatment: χ^2^_1_ = 0.033, *P* = 0.857; effect size = 0.060). For further detail regarding bee preferences by sub-treatment, see supplementary material (Tables S1-S4 and Figure S1).

**Figure 6.**
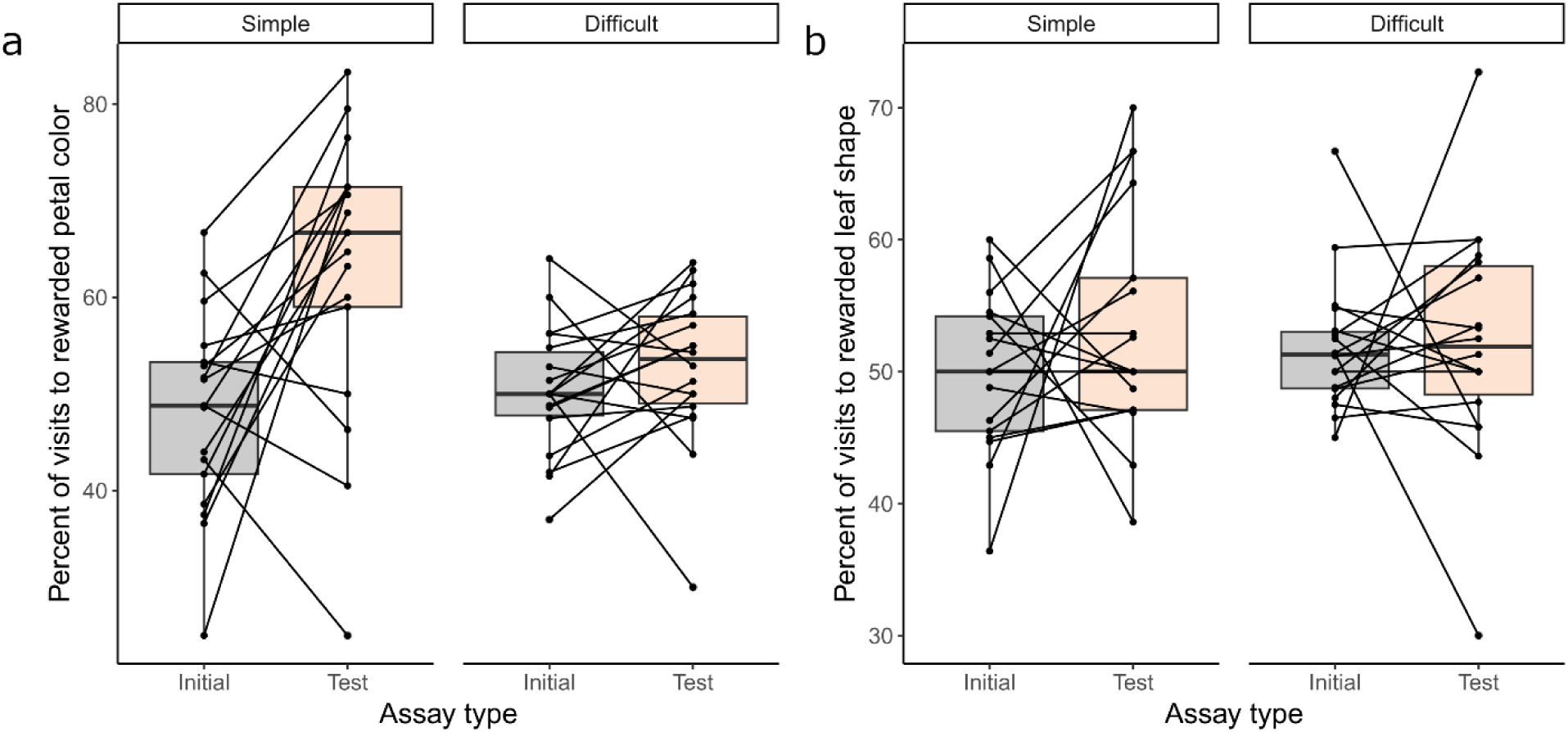
Initial and retention test preference for the trained flower type in the simple and difficult treatment (same raw data as in Figure 5), pooled by **(a)** petal color or **(b)** leaf shape. Plotted as box plots with raw data indicated via dots joined by lines showing how preference of individual bees changed with experience. *N* = 17 and 21 bees in the simple and difficult treatment, respectively.

## DISCUSSION

Our study sheds light on the role a common non-floral plant cue plays in pollinator learning. We found that generalist bumble bees can learn to associate leaf shape with a pollen food reward, but whether bees learned leaf shape depended on how readily the simultaneously available floral color cue was learned. However, our prediction for when bees would learn leaf shape did not align with our results. We hypothesized that bees would learn to discriminate flower types more slowly when presented with a difficult flower color perceptual learning task (see Niggebru gge et al. 2009). We also hypothesized that when the petal color learning task was more difficult, bees would readily learn the other available cue, leaf shape, alone or in combination with petal color, to facilitate learning. When the color perceptual learning task was difficult, bees indeed learned to discriminate flower types 29% more slowly, but they did not remember leaf shape alone or in combination with petal color. Instead, only when the color perceptual learning task was easier did bees remember the correct combination of petal color and leaf shape. Altogether, while foraging bees can attend to a non-floral cue like leaf shape, at least when compared to a flower color cue, leaf shape is less important for associative learning.

Compound floral stimuli facilitate significantly stronger learning and memory for bees (Kunze & Gumbert 2001; Stach et al. 2004; Leonard et al. 2011a; Leonard et al. 2011b; Rusch et al. 2017; Mansur et al. 2018). Our results extend prior research by demonstrating that bees can also be conditioned to a compound of flower and leaf cues. However, this compound stimulus was forgotten just one hour later by bees in our difficult color perceptual treatment. One possible explanation is that bees attempted to attend primarily to petal color, even when the color perceptual task was difficult. Consistent with this, bees in the difficult color perceptual treatment did not remember the correct leaf shape, but still showed modest recall of petal color. This bias could have occurred if the stimulus saliency of leaf shape was always significantly weaker than that of petal color in our experiment, thus resulting in weaker recall of leaf shape (see Gil-Guevara et al. 2022). Indeed, even in the simpler color perceptual learning treatment, bees learned petal color independently of leaf shape, but did not learn leaf shape independently of petal color. Altogether, our results seem to provide further evidence that differences in stimulus saliency affect learning processes and memory of compound stimuli (Katzenberger et al. 2013).

Our results also suggest that petal color overshadows learning of leaf shape. If bees only learned compound stimuli, they should have remembered only the correct leaf shape and petal color combination and avoided all other combinations. Instead, bees remembered the correct petal color and leaf shape combination, avoided flowers with the incorrect petal color (regardless of leaf shape), and showed no change in preference when petal color, but not leaf shape, matched the correct combination. This overshadowing could be a result of a bias in learning, in which bees always learn color before shape, regardless of which stimulus is more salient (e.g., Lehrer & Campan 2005; Rusch et al. 2017). Indeed, learning of flower shape is generally more difficult than learning of flower color (Menzel 1967; Lehrer et al. 1985; Lehrer 1990; Lehrer 1993), suggesting a bias in learning (Dexheimer & Dunlap 2025). Given that depending on the bee’s angle of approach to the flower, a shape cue changes substantially more than a non-iridescent color cue and thus may be a less reliable cue, a bias to learn color before shape could be functional for the bee (Gould 1993). Additionally, our experimental design potentially facilitated bees to attend to color over shape. Scent can prepare pollinators to learn a flower color cue (Leonard et al. 2011a; Leonard et al. 2011b; Leonard & Masek 2014; Mansur et al. 2018). Both rewarding and unrewarding flowers in this study were pollen-scented, which may have prepared bees to learn petal color first even when the shape learning task was potentially less perceptually difficult. Finally, perhaps flower cues generally overshadow leaf cues, given that, for instance, leaf cues will not always be reliably associated with the presence of flowers or their rewards. Future work will be required to tease apart the context and causes of overshadowing of shape cues.

How broadly representative are our results? In this study we standardized leaf surface area so that we could test whether bees specifically learned leaf shape of artificial targets. However, both leaf shape and surface area frequently differ between clades of plant species (Kidner & Umbreen 2010), which could further facilitate associative learning. Yet learning of leaf shape may also be more difficult than our findings suggest. Leaves of closely related plant species are often similar in shape and there may even be variation in leaf shape within a species (Kidner & Umbreen 2010; Richards et al. 2019), which could make learning this cue more difficult. Additionally, green leaves are often perceived by pollinators against a green background (e.g., the plant’s own foliage or the foliage of other plant species), and green achromatic contrast is especially important in associative learning by bees (Dyer & Spaethe 2008; Del Valle et al. 2025). Nonetheless, given that bees can learn to discriminate very similar flower shapes (e.g., Symington & Glover 2024) and that other insects can learn leaf shapes (e.g., Degen & Städler 1996; Wäckers & Lewis 1999; Dell’Aglio et al. 2016), learning relatively small differences in leaf shape in the context of floral foraging may also be possible.

In conclusion, given that multimodal floral cues are essential in enabling pollinators to discriminate flowers (Leonard et al. 2011a; Leonard et al. 2011b; Rusch et al. 2017; Mansur et al. 2018), the multimodal sensory billboard (Raguso 2004) may also comprise leaf cues, including potentially leaf scent (see Dufaÿ et al. 2003; Caissard et al. 2024). Whether these cues are also learned in combination with flower cues will rely on future study. In addition, the capacity of a generalist pollinator to learn a leaf cue has significant implications for plant-pollinator interactions. Pollinator cognition is broadly recognized to drive the evolution of floral cues that are more easily learned and remembered, as such cues facilitate plant reproduction (Schiestl & Johnson 2013; Russell et al. 2016; Gervasi & Schiestl 2017). Our results provide rare evidence that pollinator-mediated selection may also directly drive the evolution of leaf traits (see also Dufaÿ et al. 2003; Simon et al. 2011; Caissard et al. 2024). In particular, we expect pollinator-mediated selection on leaf cues to be strongest when co-occurring floral cues are easily learned. However, assuming that learning of flower cues generally overshadows learning of leaf cues such as shape, we would expect reduced selection on leaf cues when flower types are relatively hard to discriminate. Such a pattern might explain why, in floral mimicry systems, leaf shape (and even petal shape) often differ enough between model and mimic to be discriminable (e.g., Figure 1).

## Supporting information

R script and data files

## ACKNOWLEDGEMENTS

We thank Biobest for donating bee colonies and Antles Pollen for the pollen used in experiments. We live and work on unceded traditional territory of the Kiikaapoi, Sioux, and Osage and recognize acknowledging provenance of the land is the minimum and working to ‘land back’ is the goal.

## DECLARATION OF INTEREST

None

## CONFLICTS OF INTEREST

Not applicable

## ETHICS APPROVAL

All bumble bee experimentation was carried out in accordance with the legal and ethical standards of the USA.

## CONSENT TO PARTICIPATE

Not applicable

## CONSENT FOR PUBLICATION

Not applicable

## DATA ACCESSSIBILITY

The datasets supporting this article are available as electronic supplementary material.

## FUNDING

This research was supported by a Society for Integrative and Comparative Biology Grant in Aid of Research, a Missouri State University Graduate College Thesis Funding Grant, and a Sigma Xi Grant in Aid of Research.

## Notes

### Competing Interest Statement

The authors have declared no competing interest.

