## Supplementary material for "Take it or leaf it: bees learn leaf shape as a cue when flower color is easily learned": R script and data files: LeafLearningStatsCodes_5-16-2025.Rmd

````
---
title: "Leaf Learning Stats"
author: "MTC"
date: "`r Sys.Date()`"
output:
  pdf_document: default
  word_document: default
editor_options:
  chunk_output_type: console
---
## List of files needed to run all analyses

FILENAME1 <- "STATS_SuccessesLearningAnalysis_12-22-22_MC.csv" #Correct proportion of decisions over time
FILENAME2 <- "STATS_Visits-to-learn-by-treatment_2-20-23_MC.csv" #Number of visits to learn by treatment

Proportions of visits made to the trained rewarding cue by treatment and accompanying figures
FILENAME1 <- "PropVisits_DIFF_InitVSTest_Petal_3-30-23_MC.csv" 
FILENAME2 <- "PropVisits_DIFF_InitVSTest_Leaf_3-30-23_MC.csv"
FILENAME3 <- "PropVisits_SIMP_InitVSTest_Petal_3-30-23_MC.csv"
FILENAME4 <- "PropVisits_SIMP_InitVSTest_Leaf_3-30-23_MC.csv"
FILENAME5 <- "Analysis_Graph_Initial-vs-Final-petal-preference_7-7-2023.csv"
FILENAME6 <- "Analysis_Graph_Initial-vs-Final-leaf-preference_7-7-2023.csv"

Bar plots (visit percentages by phase and subtreatment)
FILENAME1 <- "STATS_BarplotCounts_Simp_InitialPhase-YL_12-21-23_MC.csv" # YL trained bees
FILENAME2 <- "STATS_BarplotCounts_Simp_InitialPhase-BS_12-21-23_MC.csv" # BS trained bees -simp
FILENAME3 <- "STATS_BarplotCounts_Diff_InitialPhase-PL_12-21-23_MC.csv" # PL trained bees
FILENAME4 <- "STATS_BarplotCounts_Diff_InitialPhase-BS_12-21-23_MC.csv" # BS trained bees -diff
FILENAME5 <- "STATS_BarplotCounts_Simp_TestPhase-YL_12-21-23_MC.csv" # YL trained bees
FILENAME6 <- "STATS_BarplotCounts_Simp_TestPhase-BS_12-21-23_MC.csv" # BS trained bees -simp
FILENAME7 <- "STATS_BarplotCounts_Diff_TestPhase-PL_12-21-23_MC.csv" # PL trained bees
FILENAME8 <- "STATS_BarplotCounts_Diff_TestPhase-BS_12-21-23_MC.csv" # BS trained bees -diff


Bayesian analysis of visits to each flower type by subtreatment
FILENAME1 <- "STATS_Counts_Simp_InitialPhaseAnalysis-YL_12-19-23_MC.csv" # YL trained bees
FILENAME2 <- "STATS_Counts_Simp_InitialPhaseAnalysis-BS_12-19-23_MC.csv" # BS trained bees -simp
FILENAME3 <- "STATS_Counts_Diff_InitialPhaseAnalysis-PL_12-20-23_MC.csv" # PL trained bees
FILENAME4 <- "STATS_Counts_Diff_InitialPhaseAnalysis-BS_12-20-23_MC.csv" # BS trained bees -diff
FILENAME5 <- "STATS_Counts_Simp_TestPhaseAnalysis-YL_12-20-23_MC.csv" # YL trained bees
FILENAME6 <- "STATS_Counts_Simp_TestPhaseAnalysis-BS_12-20-23_MC.csv" # BS trained bees -simp
FILENAME7 <- "STATS_Counts_Diff_TestPhaseAnalysis-PL_12-19-23_MC.csv" # PL trained bees
FILENAME8 <- "STATS_Counts_Diff_TestPhaseAnalysis-BS_12-19-23_MC.csv" # BS trained bees -diff


## CORRECT DECISION PROPORTION OVER TIME AND NUMBER OF VISITS TO LEARN BY TREATMENT

This chunk analyzes whether the bees' proportion of correct choices increases with experience, for ONLY bees that learned, by looking at the effect of experience and treatment on correct choice. This includes the random effects (colony, beeID, visit number) - nested under each other. Used GLMM and Type 2 Wald chi square tests for these. 

Data is plotted with proportion of correct visits by number of visits, with separate lines by treatment.

Also includes t-tests to analyze if the number of visits to reach learning criterion by treatment is significant

```{r}
# Changelog
# 11-3-2022 File created
# 11-15-2022 With corrected data (successes inc. tabulated) now included
# 11-15-2022 With correct rejections and correct detection data
# 12-22-2022 Updated files with missing bee data and removed data recorded from after the bee learned
# 12-22-2022 ALR updated test results for the visits to learn and checked that other stats running
# 1-5-2022 ALR updated test results for visits to learn with log transformation
# 12-19-2023 MC and ALR revised, renamed file from "Stats_learning flower and leaves analysis_12-19-2023 MC" to "Stats_CorrectDecisionsANDVisitsToLearn_12-19-2023 MC" then to "Stats_CorrectDecisionsANDVisitsToLearn_12-19-2023 MC ALR" after correcting plot to show only collected values rather than predicted
# 12-20-2023 MC corrected plot axis, cleaned up, and moved to an R markdown file

# LOAD EVERYTHING AFTER THIS

#Load the libraries:
library("cli")
library("mgcv")
library("lme4")
library("lsmeans")
library("multcomp")
library("lmerTest")
library("car")
library("effects")
library("DHARMa")
library("ggplot2")
library("DescTools")
library("sjPlot")
library("glmmTMB")

#Sets working directory automatically to where R script is saved
setwd(dirname(rstudioapi::getActiveDocumentContext()$path))

# Set the directory your plots will be in:
PLOTDIR <- dirname(rstudioapi::getActiveDocumentContext()$path)

#Name of data file:
FILENAME1 <- "STATS_SuccessesLearningAnalysis_12-22-22_MC.csv"
FILENAME2 <- "STATS_Visits-to-learn-by-treatment_2-20-23_MC.csv"

#Load data:
setwd(dirname(rstudioapi::getActiveDocumentContext()$path)); dataset1 <- read.table(FILENAME1, sep=",", header=TRUE)
setwd(dirname(rstudioapi::getActiveDocumentContext()$path)); dataset2 <- read.table(FILENAME2, sep=",", header=TRUE)

# Sanity check to make sure CSV file has correct data
head(dataset1)
head(dataset2)

#### Analysis of whether proportion correct choices increases with experience for ONLY bees that learned ####
# VisitNo treated as a repeated measure within BeeID
CorrectDecisions <- glmmTMB(CorrectChoice ~ Treatment * VisitNo + (1 | Colony / BeeID / VisitNo), family = binomial, data = dataset1)
summary(CorrectDecisions) 

# Effect size calculation
emm <- emmeans(CorrectDecisions, "Treatment")
eff_size(emm, sigma = sigma(CorrectDecisions), edf = df.residual(CorrectDecisions))

# contrast           effect.size    SE  df asymp.LCL asymp.UCL
# DIFFICULT - SIMPLE      -0.305 0.134 Inf    -0.567   -0.0421


#where it marks as significant is comparing the simple visit to difficult visit - not important to use for us rn (doesn't give info we need)
#effect of experience and treatment on correct choice, including the random effects (colony, beeID, visit number) - nested under each other
#family is binomial because correctchoice (dependent variable) is between two options (correct or incorrect)

#Type 2 sums of squares, for Wald Test reporting. As Z(<p are just approximations; Not useful with multiple factors)
#Type 2 Wald chi square tests
#Note the capital Anova. Part of the car package, runs as a Wald test
Anova(CorrectDecisions, type=2)

#### RESULTS (ones that we'd report in manuscript and presentations) ####
# Response: CorrectChoice
# Chisq Df Pr(>Chisq)  
# Treatment         2.4717  1    0.11591   #can't predict correct choice by treatment alone
# VisitNo           5.5786  1    0.01818 * #there is an effect of experience - more correct visits with experience
# Treatment:VisitNo 6.0163  1    0.01417 * #bees in one treatment are learning faster (making more correct detections) than the other treatment - effect of treatment depends on experience
#   ---
#   Signif. codes:  0 ‘***’ 0.001 ‘**’ 0.01 ‘*’ 0.05 ‘.’ 0.1 ‘ ’ 1

# Checking model assumptions via DHARMa
simulationOutput1 <- simulateResiduals(fittedModel = CorrectDecisions, n = 10000)
plot(simulationOutput1) # Plot of QQ plot good
testUniformity(simulationOutput = simulationOutput1) # No significant deviation
testDispersion(simulationOutput1) # Dispersion good
testZeroInflation(simulationOutput1) # Zero inflation good

# Model assumptions via sjPlot
# Normality of residuals and outliers
plot_model(CorrectDecisions, type = "diag") # Looks normal

# Sjplot effect sizes, data, and predicted values
# Does not fully work
plot_model(CorrectDecisions) #not super useful for us, might not even work properly for a binomial model
plot_model(CorrectDecisions, type="pred", terms=c("Treatment"), show.data = T) #
# plot_model(CorrectDecisions, type="pred", terms=c("Treatment", "VisitNo"), show.data = T) # What does this mean? Avery doesn't know...

#### Plot model without predictions ####
# Rename the figure legends
dataset1$Treatment = plyr::mapvalues(dataset1$Treatment, from = c("SIMPLE", "DIFFICULT"), to = c("Simple", "Difficult"))

#Plot model with SE
Figure1 <- ggplot(dataset1, aes(x=VisitNo, y=CorrectChoice, color=Treatment, fill=Treatment)) + 
  geom_smooth(method = "lm", se = TRUE) + 
  scale_linetype_manual(values=c("solid","solid")) + 
  ylab("Proportion correct visits") + xlab("Number of visits") + theme_classic() + 
  theme(text=element_text(size=14)) +
  scale_color_viridis_d(name = "Learning \ntreatment", end = 0.7, option = "inferno") + 
  scale_fill_viridis_d(name = "Learning \ntreatment", end =0.7, option = 'inferno') +
  theme(legend.position  =c(0.5, 0.9), legend.direction = "horizontal", 
        legend.background = element_rect(fill="transparent", colour = NA),
        strip.background = element_blank(),
        legend.title = element_text(size = 12), legend.text = element_text(size = 12),
        panel.spacing.x = unit(1, "lines"), 
        plot.margin = margin(c(0.2,0.51, 0.2, 0.2), unit = "cm")) + 
  xlim(0,170) + ylim(0,1)
plot(Figure1) #To see the figure in R

# Set date
DATE <- "`r Sys.Date()`"

tiff(filename=paste("Fig_CorrectChoiceModel_", DATE, ".tif", sep=""), units="mm", width=150, height=100, res=1500)

# Close the plotting device
setwd(PLOTDIR); 
dev.off()

#### Analysis of number of visits to reach learning criterion in the learning phase ####
#how quickly they learned by treatment

# Run t-test
t.test(dataset2$Log_DIFFICULT,dataset2$Log_SIMPLE,paired=FALSE)
#t.test(dataset5$DIFFICULT,dataset5$SIMPLE,paired=FALSE)

# Results
# t = 1.64, df = 35.757, p-value = 0.1098


# Model assumptions
# Equal variance of the two populations
# Run model var.test
var.test(dataset2$Log_DIFFICULT,dataset2$Log_SIMPLE)
#var.test(dataset5$DIFFICULT,dataset5$SIMPLE)
# Comes back as significantly different w/o the log transformation

# Test of normality for each population
shapiro.test(dataset2$Log_DIFFICULT)
# W = 0.92655, p-value = 0.1174 #

shapiro.test(dataset2$Log_SIMPLE)
# W = 0.96222, p-value = 0.6736 # 


# Since t-test violates assumptions, use wilcoxon signed rank test
wilcox.test(dataset2$SIMPLE,dataset2$DIFFICULT,paired=FALSE)

# Results
# W = 126, p-value = 0.1268 # 

#non-significant effect here, bees did not make more visits in one treatment than the other to meet our criteria
#can either report this or not

```

##INITIAL VS TEST PREFERENCES FOR PETAL OR LEAF

Comparing pooled (trained rewarding) initial to test preferences by treatment with accompanying plots

Stats and plots for comparing how individual bee preferences changed from their trained rewarding flower (split by petal and leaf) in the initial and test phase and compared by treatment.

```{r}
# Changelog

# 11-3-2022   File created
# 11-15-2022 With corrected data (successes inc. tabulated) now included
# 11-15-2022 With correct rejections and correct detection data
# 12-22-2022 Updated files with missing bee data and removed data recorded from after the bee learned
# 12-22-2022 ALR updated test results for the visits to learn and checked that other stats running
# 1-5-2022 ALR updated test results for visits to learn with log transformation
# 3-8-2023 MC created modified file for initial vs test analysis
# 3-9-2023 MC made edits to run initial vs test analyses
# 3-29-2023 MC updated difficult and rewardingpercentchange datasets to remove outliers (B22 and B4)
# 3-30-2023 MC added violin plot code and ColonyID to datasheets
# 4-6-2023 MC edited violin plot axis names and spans
# 12-19-2023 MC and ALR cut redundant code (violin plots, combination cues) and files, fixed Colony stuff
# 12-20-2023 MC moved to R markdown file and added relevant plot code from "Graph-Initial-vs-Test-paired-data_7-7-2023"
#2-10-2024 MC added separate analyses for sub-treatment level preference changes

# LOAD EVERYTHING AFTER THIS

#Load the libraries:
library("mgcv")
library("lme4")
library("lsmeans")
library("multcomp")
library("lmerTest")
library("car")
library("effects")
library("DHARMa")
library("ggplot2")
library("DescTools")
library("sjPlot")
library("glmmTMB")
library("mclust")
library("RVAideMemoire")
library("tidyverse")
library("hrbrthemes")
library("viridis")
library("vioplot")

#Name of data file:
FILENAME1 <- "PropVisits_DIFF_InitVSTest_Petal_3-30-23_MC.csv"
FILENAME2 <- "PropVisits_DIFF_InitVSTest_Leaf_3-30-23_MC.csv"
FILENAME3 <- "PropVisits_SIMP_InitVSTest_Petal_3-30-23_MC.csv"
FILENAME4 <- "PropVisits_SIMP_InitVSTest_Leaf_3-30-23_MC.csv"
FILENAME5 <- "Analysis_Graph_Initial-vs-Final-petal-preference_7-7-2023.csv"
FILENAME6 <- "Analysis_Graph_Initial-vs-Final-leaf-preference_7-7-2023.csv"

#Sets working directory automatically to where R script is saved
setwd(dirname(rstudioapi::getActiveDocumentContext()$path))

# Set the directory your plots will be in:
PLOTDIR <- dirname(rstudioapi::getActiveDocumentContext()$path)

#Load data:
setwd(dirname(rstudioapi::getActiveDocumentContext()$path)); dataset1 <- read.table(FILENAME1, sep=",", header=TRUE)
setwd(dirname(rstudioapi::getActiveDocumentContext()$path)); dataset2 <- read.table(FILENAME2, sep=",", header=TRUE)
setwd(dirname(rstudioapi::getActiveDocumentContext()$path)); dataset3 <- read.table(FILENAME3, sep=",", header=TRUE)
setwd(dirname(rstudioapi::getActiveDocumentContext()$path)); dataset4 <- read.table(FILENAME4, sep=",", header=TRUE)
setwd(dirname(rstudioapi::getActiveDocumentContext()$path)); dataset5 <- read.table(FILENAME5, sep=",", header=TRUE)
setwd(dirname(rstudioapi::getActiveDocumentContext()$path)); dataset6 <- read.table(FILENAME6, sep=",", header=TRUE)

# Sanity check to make sure CSV file has correct data
head(dataset1)
head(dataset2)
head(dataset3)
head(dataset4)
head(dataset5)
head(dataset6)

#### Analysis of whether bee petal color preference differs by phase, difficult treatment ####

DifficultPetalProp <- glmmTMB(TrainedRewardingPetal ~ Phase + (1 | ColonyID / BeeID), data = dataset1)

summary(DifficultPetalProp)

# Effect size calculation
emm <- emmeans(DifficultPetalProp, "Phase")
eff_size(emm, sigma = sigma(DifficultPetalProp), edf = df.residual(DifficultPetalProp))

# contrast       effect.size    SE df lower.CL upper.CL
# Initial - Test      -0.349 0.336 32    -1.03    0.336

#Type 2 sums of squares, for Wald Test reporting. As Z(<p are just approximations; Not useful with multiple factors)
#Note the capital Anova.
Anova(DifficultPetalProp)

## RESULTS
# Response: TrainedRewardingPetal
#       Chisq Df   Pr(>Chisq)
# Phase 1.0948  1     0.2954

# Model assumptions via DHARMa
simulationOutput1 <- simulateResiduals(fittedModel = DifficultPetalProp, n = 10000)
plot(simulationOutput1) # Plot of QQ plot looks good
testUniformity(simulationOutput = simulationOutput1) # good
testDispersion(simulationOutput1) # Dispersion good
testZeroInflation(simulationOutput1) # no significant deviation, no problem

# Bees learned in the training phase, but this shows that they didn't remember the petal color in the difficult test phase

#### Analysis of whether bee petal color preference differs by phase, simple treatment ####
SimplePetalProp <- glmmTMB(TrainedRewardingPetal ~ Phase + (1 | ColonyID / BeeID), data = dataset3)
summary(SimplePetalProp)

# Effect size calculation
emm <- emmeans(SimplePetalProp, "Phase")
eff_size(emm, sigma = sigma(SimplePetalProp), edf = df.residual(SimplePetalProp))

# contrast       effect.size    SE df lower.CL upper.CL
# Initial - Test       -1.18 0.376 29    -1.94   -0.406

#Type 2 sums of squares, for Wald Test reporting. As Z(<p are just approximations; Not useful with multiple factors)
#Note the capital Anova.
Anova(SimplePetalProp)

## RESULTS
# Response: TrainedRewardingPetal
#        Chisq Df Pr(>Chisq)    
# Phase 11.743  1  0.0006107 ***
#  ---
#  Signif. codes:  0 ‘***’ 0.001 ‘**’ 0.01 ‘*’ 0.05 ‘.’ 0.1 ‘ ’ 1


# Model assumptions via DHARMa
simulationOutput3 <- simulateResiduals(fittedModel = SimplePetalProp, n = 10000)
plot(simulationOutput3) # Plot of QQ plot has significant deviation
testUniformity(simulationOutput = simulationOutput3) # (Uniformity) Deviation n.s
testDispersion(simulationOutput3) # Dispersion good
testZeroInflation(simulationOutput3) # Zero inflation ?

# bees did remember correct petal color for the simple treatment

#### Plotting petal color preference changes ####
# Figure with paired data
# Rename the figure legends
#dataset2$Phase = plyr::mapvalues(dataset2$Phase, from = c("Initial", "Test"), to = c("Naïve", "Experienced"))
dataset5$Treatment = plyr::mapvalues(dataset5$Treatment, from = c("Difficult", "Simple"), 
                                     to = c("Difficult","Simple"))

# Reorder the experience levels (treatment variables)
dataset5$Phase <- factor(dataset5$Phase, levels = c("Initial","Test"))

# Reorder the experience levels (treatment variables)
dataset5$Treatment <- factor(dataset5$Treatment, levels = c("Simple","Difficult"))

# Plot figure
FigurePetal <- ggplot(dataset5, aes(x = Phase, y = Preference)) + 
  geom_boxplot(aes(fill = Phase), alpha = .2) +
  geom_line(aes(group = BeeID)) + 
  geom_point(size = 1) + 
  facet_wrap(~ Treatment) +
  ylab("Percent of visits to rewarded petal color") + xlab("Assay type") + theme_classic() + 
  theme(text=element_text(size=14)) +
  scale_color_viridis_d(name = "Experience \nlevel", end = 0.7, option = "inferno") + 
  scale_fill_viridis_d(name = "Experience \nlevel", end =0.7, option = 'inferno') +
  theme(legend.position  = c(-1.0, -1.0), legend.direction = "horizontal", 
        legend.background = element_rect(fill="transparent", colour = NA),
        legend.title = element_text(size = 0), legend.text = element_text(size = 0),
        panel.spacing.x = unit(1, "lines"), 
        plot.margin = margin(c(0.2,0.2, 0.2, 0.2), unit = "cm"))
plot(FigurePetal) #To see the figure in R

# Set date
DATE <- "3-26-2025"

tiff(filename=paste("Fig_Initial-vs-Final-petal-Preference_", DATE, ".tif", sep=""), units="mm", width=85, height=85, res=1200)

# Close the plotting device
setwd(PLOTDIR); 
dev.off()


#### Analysis of whether bee leaf shape preference differs by phase, difficult treatment ####
#DifficultLeafProp <- glmmTMB(TrainedRewardingLeaf ~ Phase + (1 | BeeID), data = dataset2)
DifficultLeafProp <- glmmTMB(TrainedRewardingLeaf ~ Phase + (1 | ColonyID / BeeID), data = dataset2)
summary(DifficultLeafProp)

# Effect size calculation
emm <- emmeans(DifficultLeafProp, "Phase")
eff_size(emm, sigma = sigma(DifficultLeafProp), edf = df.residual(DifficultLeafProp))

# contrast       effect.size    SE df lower.CL upper.CL
# Initial - Test     -0.0602 0.333 31    -0.74     0.62


#Type 2 sums of squares, for Wald Test reporting. As Z(<p are just approximations; Not useful with multiple factors)
#Note the capital Anova.
Anova(DifficultLeafProp)

## RESULTS
# Response: TrainedRewardingLeaf
#        Chisq Df Pr(>Chisq)
#Phase 0.0326  1     0.8566

# Model assumptions via DHARMa
simulationOutput2 <- simulateResiduals(fittedModel = DifficultLeafProp, n = 10000)
plot(simulationOutput2) # Plot of QQ plot good
testUniformity(simulationOutput = simulationOutput2) # good
testDispersion(simulationOutput2) # Dispersion good
testZeroInflation(simulationOutput2) # Zero inflation good

# difficult bees did not remember rewarding leaf shape

#### Analysis of whether bee leaf shape preference differs by phase, simple treatment ####
#SimpleLeafProp <- glmmTMB(TrainedRewardingLeaf ~ Phase + (1 | BeeID), data = dataset4)
SimpleLeafProp <- glmmTMB(TrainedRewardingLeaf ~ Phase + (1 | ColonyID / BeeID), data = dataset4) #giving an error but seems fine?
summary(SimpleLeafProp)

#Type 2 sums of squares, for Wald Test reporting. As Z(<p are just approximations; Not useful with multiple factors)
#Note the capital Anova.
Anova(SimpleLeafProp)

## RESULTS
# Response: TrainedRewardingLeaf
#        Chisq Df Pr(>Chisq)
#Phase 1.8028  1     0.1794

# Model assumptions via DHARMa
simulationOutput4 <- simulateResiduals(fittedModel = SimpleLeafProp, n = 10000)
plot(simulationOutput4) # Plot of QQ plot has 
testUniformity(simulationOutput = simulationOutput4) # (Uniformity) Deviation
testDispersion(simulationOutput4) # Dispersion?
testZeroInflation(simulationOutput4) # Zero inflation ?

# bees didn't remember correct leaf shape for the simple treatment

#### Plotting leaf shape preference changes ####
# Figure with paired data
# Rename the figure legends
#dataset2$Phase = plyr::mapvalues(dataset2$Phase, from = c("Initial", "Test"), to = c("Naïve", "Experienced"))
dataset6$Treatment = plyr::mapvalues(dataset6$Treatment, from = c("Difficult", "Simple"), 
                                     to = c("Difficult","Simple"))

# Reorder the experience levels (treatment variables)
dataset6$Phase <- factor(dataset6$Phase, levels = c("Initial","Test"))

# Reorder the experience levels (treatment variables)
dataset6$Treatment <- factor(dataset6$Treatment, levels = c("Simple","Difficult"))

# Plot figure
FigureLeaf <- ggplot(dataset6, aes(x = Phase, y = Preference)) + 
  geom_boxplot(aes(fill = Phase), alpha = .2) +
  geom_line(aes(group = BeeID)) + 
  geom_point(size = 1) + 
  facet_wrap(~ Treatment) +
  ylab("Percent of visits to rewarded leaf shape") + xlab("Assay type") + theme_classic() + 
  theme(text=element_text(size=14)) +
  scale_color_viridis_d(name = "Experience \nlevel", end = 0.7, option = "inferno") + 
  scale_fill_viridis_d(name = "Experience \nlevel", end =0.7, option = 'inferno') +
  theme(legend.position  = c(-1.0, -1.0), legend.direction = "horizontal", 
        legend.background = element_rect(fill="transparent", colour = NA),
        legend.title = element_text(size = 0), legend.text = element_text(size = 0),
        panel.spacing.x = unit(1, "lines"), 
        plot.margin = margin(c(0.2,0.2, 0.2, 0.2), unit = "cm"))
plot(FigureLeaf) #To see the figure in R

# Set date
DATE <- "3-26-2025"

tiff(filename=paste("Fig_Initial-vs-Final-leaf-Preference_", DATE, ".tif", sep=""), units="mm", width=85, height=85, res=1200)

# Close the plotting device
setwd(PLOTDIR); 
dev.off()


####Re-runnning analyses for sub-treatment levels####

#Name of data file:
FILENAME7 <- "PropVisits_DIFF_InitVSTest_Petal-BS_2-10-24_MC.csv"
FILENAME8 <- "PropVisits_DIFF_InitVSTest_Leaf-BS_2-10-24_MC.csv"
FILENAME9 <- "PropVisits_DIFF_InitVSTest_Petal-PL_2-10-24_MC.csv"
FILENAME10 <- "PropVisits_DIFF_InitVSTest_Leaf-PL_2-10-24_MC.csv"
FILENAME11 <- "PropVisits_SIMP_InitVSTest_Petal-BS_2-10-24_MC.csv"
FILENAME12 <- "PropVisits_SIMP_InitVSTest_Leaf-BS_2-10-24_MC.csv"
FILENAME13 <- "PropVisits_SIMP_InitVSTest_Petal-YL_2-10-24_MC.csv"
FILENAME14 <- "PropVisits_SIMP_InitVSTest_Leaf-YL_2-10-24_MC.csv"
#FILENAME5 <- "Analysis_Graph_Initial-vs-Final-petal-preference_7-7-2023.csv"
#FILENAME6 <- "Analysis_Graph_Initial-vs-Final-leaf-preference_7-7-2023.csv"

#Sets working directory automatically to where R script is saved
setwd(dirname(rstudioapi::getActiveDocumentContext()$path))

# Set the directory your plots will be in:
PLOTDIR <- dirname(rstudioapi::getActiveDocumentContext()$path)

#Load data:
setwd(dirname(rstudioapi::getActiveDocumentContext()$path)); dataset7 <- read.table(FILENAME7, sep=",", header=TRUE)
setwd(dirname(rstudioapi::getActiveDocumentContext()$path)); dataset8 <- read.table(FILENAME8, sep=",", header=TRUE)
setwd(dirname(rstudioapi::getActiveDocumentContext()$path)); dataset9 <- read.table(FILENAME9, sep=",", header=TRUE)
setwd(dirname(rstudioapi::getActiveDocumentContext()$path)); dataset10 <- read.table(FILENAME10, sep=",", header=TRUE)
setwd(dirname(rstudioapi::getActiveDocumentContext()$path)); dataset11 <- read.table(FILENAME11, sep=",", header=TRUE)
setwd(dirname(rstudioapi::getActiveDocumentContext()$path)); dataset12 <- read.table(FILENAME12, sep=",", header=TRUE)
setwd(dirname(rstudioapi::getActiveDocumentContext()$path)); dataset13 <- read.table(FILENAME13, sep=",", header=TRUE)
setwd(dirname(rstudioapi::getActiveDocumentContext()$path)); dataset14 <- read.table(FILENAME14, sep=",", header=TRUE)


# Sanity check to make sure CSV file has correct data
head(dataset7)
head(dataset8)
head(dataset9)
head(dataset10)
head(dataset11)
head(dataset12)
head(dataset13)
head(dataset14)


#### Analysis of whether bee petal color preference differs by phase, difficult treatment BS ####

# DifficultPetalProp <- glmmTMB(TrainedRewardingPetal ~ Phase + (1 | BeeID), data = dataset1)
DifficultPetalPropBS <- glmmTMB(TrainedRewardingPetal ~ Phase + (1 | ColonyID / BeeID), data = dataset7)

summary(DifficultPetalPropBS)

# Effect size calculation
emm <- emmeans(DifficultPetalPropBS, "Phase")
eff_size(emm, sigma = sigma(DifficultPetalPropBS), edf = df.residual(DifficultPetalPropBS))

# contrast       effect.size    SE df lower.CL upper.CL
# Initial - Test     -0.0307 0.447 15   -0.984    0.923


#Type 2 sums of squares, for Wald Test reporting. As Z(<p are just approximations; Not useful with multiple factors)
#Note the capital Anova.
Anova(DifficultPetalPropBS)

## RESULTS
# Response: TrainedRewardingPetal
#       Chisq Df Pr(>Chisq)
# Phase 0.0047  1     0.9453

# Model assumptions via DHARMa
simulationOutput7 <- simulateResiduals(fittedModel = DifficultPetalPropBS, n = 10000)
plot(simulationOutput7) # Plot of QQ plot looks good
testUniformity(simulationOutput = simulationOutput7) # good
testDispersion(simulationOutput7) # Dispersion good
testZeroInflation(simulationOutput7) # no problem

# Model assumptions via sjPlot
# Normality of residuals and outliers
plot_model(DifficultPetalPropBS, type = "diag") # Looks interesting? Avery also doesn't know what it means

# Sjplot effect sizes, data, and predicted values
# Does not fully work
plot_model(DifficultPetalPropBS)
plot_model(DifficultPetalPropBS, type="pred", terms=c("Phase"), show.data = T) #
# plot_model(DifficultPetalProp, type="pred", terms=c("Phase", "TrainedRewardingPetal"), show.data = T) # What does this mean? Avery doesn't know...

# Bees learned in the training phase, but this shows that they didn't remember the petal color in the difficult test phase - BS


#### Analysis of whether bee petal color preference differs by phase, difficult treatment PL####
#DifficultPetalPropPL <- glmmTMB(TrainedRewardingPetal ~ Phase + (1 | ColonyID / BeeID), data = dataset9)
DifficultPetalPropPL <- glmmTMB(TrainedRewardingPetal ~ Phase + (1 | BeeID), data = dataset9)

summary(DifficultPetalPropPL)

# Effect size calculation
emm <- emmeans(DifficultPetalPropPL, "Phase")
eff_size(emm, sigma = sigma(DifficultPetalPropPL), edf = df.residual(DifficultPetalPropPL))

# contrast       effect.size    SE df lower.CL upper.CL
# Initial - Test     -0.0307 0.447 15   -0.984    0.923

#Type 2 sums of squares, for Wald Test reporting. As Z(<p are just approximations; Not useful with multiple factors)
#Note the capital Anova.
Anova(DifficultPetalPropPL)

## RESULTS
# Response: TrainedRewardingPetal
#        Chisq Df Pr(>Chisq)  
# Phase 3.7853  1    0.05171 .      

# Model assumptions via DHARMa
simulationOutput9 <- simulateResiduals(fittedModel = DifficultPetalPropPL, n = 10000)
plot(simulationOutput9) # Plot of QQ plot looks good
testUniformity(simulationOutput = simulationOutput9) # good
testDispersion(simulationOutput9) # Dispersion good
testZeroInflation(simulationOutput9) # no problem

# Model assumptions via sjPlot
# Normality of residuals and outliers
plot_model(DifficultPetalPropPL, type = "diag") # Looks interesting? Avery also doesn't know what it means

# Sjplot effect sizes, data, and predicted values
# Does not fully work
plot_model(DifficultPetalPropPL)
plot_model(DifficultPetalPropPL, type="pred", terms=c("Phase"), show.data = T) #
# plot_model(DifficultPetalProp, type="pred", terms=c("Phase", "TrainedRewardingPetal"), show.data = T) # What does this mean? Avery doesn't know...

# Bees learned in the training phase, but this shows that they didn't remember (just barely) the petal color in the difficult test phase - PL


#### Analysis of whether bee leaf shape preference differs by phase, difficult treatment BS ####
DifficultLeafPropBS <- glmmTMB(TrainedRewardingLeaf ~ Phase + (1 | BeeID), data = dataset8)

summary(DifficultLeafPropBS)

#Type 2 sums of squares, for Wald Test reporting. As Z(<p are just approximations; Not useful with multiple factors)
#Note the capital Anova.
Anova(DifficultLeafPropBS)

## RESULTS
# Response: TrainedRewardingLeaf
#       Chisq Df Pr(>Chisq)
# Phase 0.2914  1     0.5893

# Model assumptions via DHARMa
simulationOutput8 <- simulateResiduals(fittedModel = DifficultLeafPropBS, n = 10000)
plot(simulationOutput8) # Plot of QQ plot looks good
testUniformity(simulationOutput = simulationOutput8) # good
testDispersion(simulationOutput8) # Dispersion good
testZeroInflation(simulationOutput8) # no problem

# Model assumptions via sjPlot
# Normality of residuals and outliers
plot_model(DifficultLeafPropBS, type = "diag") # Looks interesting? Avery also doesn't know what it means

# Sjplot effect sizes, data, and predicted values
# Does not fully work
plot_model(DifficultLeafPropBS)
plot_model(DifficultLeafPropBS, type="pred", terms=c("Phase"), show.data = T) #
# plot_model(DifficultPetalProp, type="pred", terms=c("Phase", "TrainedRewardingPetal"), show.data = T) # What does this mean? Avery doesn't know...

# Bees learned in the training phase, but this shows that they didn't remember the leaf shape in the difficult test phase - BS


#### Analysis of whether bee leaf shape preference differs by phase, difficult treatment PL####
#DifficultPetalPropPL <- glmmTMB(TrainedRewardingPetal ~ Phase + (1 | ColonyID / BeeID), data = dataset9)
DifficultLeafPropPL <- glmmTMB(TrainedRewardingLeaf ~ Phase + (1 | BeeID), data = dataset10)

summary(DifficultLeafPropPL)

#Type 2 sums of squares, for Wald Test reporting. As Z(<p are just approximations; Not useful with multiple factors)
#Note the capital Anova.
Anova(DifficultLeafPropPL)

## RESULTS
# Response: TrainedRewardingLeaf
#        Chisq Df Pr(>Chisq)  
# Phase 0.1472  1     0.7012      

# Model assumptions via DHARMa
simulationOutput10 <- simulateResiduals(fittedModel = DifficultLeafPropPL, n = 10000)
plot(simulationOutput10) # Plot of QQ plot looks good
testUniformity(simulationOutput = simulationOutput10) # good
testDispersion(simulationOutput10) # Dispersion good
testZeroInflation(simulationOutput10) # no problem

# Model assumptions via sjPlot
# Normality of residuals and outliers
plot_model(DifficultLeafPropPL, type = "diag") # Looks interesting? Avery also doesn't know what it means

# Sjplot effect sizes, data, and predicted values
# Does not fully work
plot_model(DifficultLeafPropPL)
plot_model(DifficultLeafPropPL, type="pred", terms=c("Phase"), show.data = T) #
# plot_model(DifficultPetalProp, type="pred", terms=c("Phase", "TrainedRewardingPetal"), show.data = T) # What does this mean? Avery doesn't know...

# Bees learned in the training phase, but this shows that they didn't remember leaf shape in the difficult test phase - PL


#### Analysis of whether bee petal color preference differs by phase, simple treatment BS ####

SimplePetalPropBS <- glmmTMB(TrainedRewardingPetal ~ Phase + (1 | ColonyID / BeeID), data = dataset11)

summary(SimplePetalPropBS)

#Type 2 sums of squares, for Wald Test reporting. As Z(<p are just approximations; Not useful with multiple factors)
#Note the capital Anova.
Anova(SimplePetalPropBS)

## RESULTS
# Response: TrainedRewardingPetal
#       Chisq Df Pr(>Chisq)
# Phase 4.74  1    0.02947 *

# Model assumptions via DHARMa
simulationOutput11 <- simulateResiduals(fittedModel = SimplePetalPropBS, n = 10000)
plot(simulationOutput11) # Plot of QQ plot looks good
testUniformity(simulationOutput = simulationOutput11) # good
testDispersion(simulationOutput11) # Dispersion good
testZeroInflation(simulationOutput11) # no problem

# Model assumptions via sjPlot
# Normality of residuals and outliers
plot_model(SimplePetalPropBS, type = "diag") # Looks interesting? Avery also doesn't know what it means

# Sjplot effect sizes, data, and predicted values
# Does not fully work
plot_model(SimplePetalPropBS)
plot_model(SimplePetalPropBS, type="pred", terms=c("Phase"), show.data = T) #
# plot_model(DifficultPetalProp, type="pred", terms=c("Phase", "TrainedRewardingPetal"), show.data = T) # What does this mean? Avery doesn't know...

# Bees learned in the training phase, and did remember the petal color in the simple test phase - BS


#### Analysis of whether bee petal color preference differs by phase, simple treatment YL####
#DifficultPetalPropPL <- glmmTMB(TrainedRewardingPetal ~ Phase + (1 | ColonyID / BeeID), data = dataset9)
SimplePetalPropYL <- glmmTMB(TrainedRewardingPetal ~ Phase + (1 | ColonyID / BeeID), data = dataset13)

summary(SimplePetalPropYL)

#Type 2 sums of squares, for Wald Test reporting. As Z(<p are just approximations; Not useful with multiple factors)
#Note the capital Anova.
Anova(SimplePetalPropYL)

## RESULTS
# Response: TrainedRewardingPetal
#        Chisq Df Pr(>Chisq)  
# Phase 10.53  1   0.001174 **   

#w/ colony:
# Phase 14.067  1  0.0001764 ***

# Model assumptions via DHARMa
simulationOutput13 <- simulateResiduals(fittedModel = SimplePetalPropYL, n = 10000)
plot(simulationOutput13) # Plot of QQ plot looks good
testUniformity(simulationOutput = simulationOutput13) # good
testDispersion(simulationOutput13) # Dispersion good
testZeroInflation(simulationOutput13) # no problem

# Model assumptions via sjPlot
# Normality of residuals and outliers
plot_model(SimplePetalPropYL, type = "diag") # Looks interesting? Avery also doesn't know what it means

# Sjplot effect sizes, data, and predicted values
# Does not fully work
plot_model(SimplePetalPropYL)
plot_model(SimplePetalPropYL, type="pred", terms=c("Phase"), show.data = T) #
# plot_model(DifficultPetalProp, type="pred", terms=c("Phase", "TrainedRewardingPetal"), show.data = T) # What does this mean? Avery doesn't know...

# Bees learned in the training phase and did remember the petal color in the simple test phase - YL


#### Analysis of whether bee leaf shape preference differs by phase, simple treatment BS ####
SimpleLeafPropBS <- glmmTMB(TrainedRewardingLeaf ~ Phase + (1 | ColonyID / BeeID), data = dataset12)
#SimpleLeafPropBS <- glmmTMB(TrainedRewardingLeaf ~ Phase + (1 | BeeID), data = dataset12)

summary(SimpleLeafPropBS)

#Type 2 sums of squares, for Wald Test reporting. As Z(<p are just approximations; Not useful with multiple factors)
#Note the capital Anova.
Anova(SimpleLeafPropBS)

## RESULTS
# Response: TrainedRewardingLeaf
#       Chisq Df Pr(>Chisq)
# Phase 1.2849  1      0.257

# Model assumptions via DHARMa
simulationOutput12 <- simulateResiduals(fittedModel = SimpleLeafPropBS, n = 10000)
plot(simulationOutput12) # Plot of QQ plot looks good
testUniformity(simulationOutput = simulationOutput12) # good
testDispersion(simulationOutput12) # Dispersion good
testZeroInflation(simulationOutput12) # no problem

# Model assumptions via sjPlot
# Normality of residuals and outliers
plot_model(SimpleLeafPropBS, type = "diag") # Looks interesting? Avery also doesn't know what it means

# Sjplot effect sizes, data, and predicted values
# Does not fully work
plot_model(SimpleLeafPropBS)
plot_model(SimpleLeafPropBS, type="pred", terms=c("Phase"), show.data = T) #
# plot_model(DifficultPetalProp, type="pred", terms=c("Phase", "TrainedRewardingPetal"), show.data = T) # What does this mean? Avery doesn't know...

# Bees learned in the training phase, but this shows that they didn't remember the leaf shape in the simple test phase - BS


#### Analysis of whether bee leaf shape preference differs by phase, difficult treatment PL####
#DifficultPetalPropPL <- glmmTMB(TrainedRewardingPetal ~ Phase + (1 | ColonyID / BeeID), data = dataset9)
SimpleLeafPropYL <- glmmTMB(TrainedRewardingLeaf ~ Phase + (1 | ColonyID / BeeID), data = dataset14)

summary(SimpleLeafPropYL)

#Type 2 sums of squares, for Wald Test reporting. As Z(<p are just approximations; Not useful with multiple factors)
#Note the capital Anova.
Anova(SimpleLeafPropYL)

## RESULTS
# Response: TrainedRewardingLeaf
#        Chisq Df Pr(>Chisq)  
# Phase 0.5524  1     0.4574      

# Model assumptions via DHARMa
simulationOutput14 <- simulateResiduals(fittedModel = SimpleLeafPropYL, n = 10000)
plot(simulationOutput14) # Plot of QQ plot looks good
testUniformity(simulationOutput = simulationOutput14) # good
testDispersion(simulationOutput14) # Dispersion good
testZeroInflation(simulationOutput14) # no problem

# Model assumptions via sjPlot
# Normality of residuals and outliers
plot_model(SimpleLeafPropYL, type = "diag") # Looks interesting? Avery also doesn't know what it means

# Sjplot effect sizes, data, and predicted values
# Does not fully work
plot_model(SimpleLeafPropYL)
plot_model(SimpleLeafPropYL, type="pred", terms=c("Phase"), show.data = T) #
# plot_model(DifficultPetalProp, type="pred", terms=c("Phase", "TrainedRewardingPetal"), show.data = T) # What does this mean? Avery doesn't know...

# Bees learned in the training phase, but this shows that they didn't remember leaf shape in the simple test phase - YL


#### Plotting petal color preference changes by sub-treatment####

FILENAME15 <- "Analysis_Graph_Initial-vs-Final-petal-preference-SIMPLE_2-18-2024.csv"
FILENAME16 <- "Analysis_Graph_Initial-vs-Final-petal-preference-DIFF_2-18-2024.csv"
dataset15 <- read.csv("~/LL stats/Analysis_Graph_Initial-vs-Final-petal-preference-SIMPLE_2-18-2024.csv", sep=",", header=TRUE)
dataset16 <- read.csv("~/LL stats/Analysis_Graph_Initial-vs-Final-petal-preference-DIFF_2-18-2024.csv", sep=",", header=TRUE)

head(dataset15)
head(dataset16)


# Figure with paired data, simple sub-treatments
# Rename the figure legends
dataset15$Subtreatment = plyr::mapvalues(dataset15$Subtreatment, from = c("BS", "YL"), 
                                     to = c("BS","YL"))

# Reorder the experience levels (treatment variables)
dataset15$Phase <- factor(dataset15$Phase, levels = c("Initial","Test"))

# Reorder the experience levels (treatment variables)
dataset15$Subtreatment <- factor(dataset15$Subtreatment, levels = c("BS","YL"))

# Plot figure
FigurePetalSIMPLEsubs <- ggplot(dataset15, aes(x = Phase, y = Preference)) + 
  geom_boxplot(aes(fill = Phase), alpha = .2) +
  geom_line(aes(group = BeeID)) + 
  geom_point(size = 1) + 
  facet_wrap(~ Subtreatment) +
  ylab("Preference for Trained (S+) Petal Color") + xlab("Assay type") + theme_classic() + 
  theme(text=element_text(size=14)) +
  scale_color_viridis_d(name = "Experience \nlevel", end = 0.7, option = "inferno") + 
  scale_fill_viridis_d(name = "Experience \nlevel", end =0.7, option = 'inferno') +
  theme(legend.position  = c(-1.0, -1.0), legend.direction = "horizontal", 
        legend.background = element_rect(fill="transparent", colour = NA),
        legend.title = element_text(size = 0), legend.text = element_text(size = 0),
        panel.spacing.x = unit(1, "lines"), 
        plot.margin = margin(c(0,0, 0, 0), unit = "cm"))
plot(FigurePetalSIMPLEsubs) #To see the figure in R

# Set date
DATE <- "`r Sys.Date()`"

tiff(filename=paste("Fig_Initial-vs-Final-petal-Preference_SIMPLE-subs", DATE, ".tif", sep=""), units="mm", width=200, height=200, res=1200)

plot(FigurePetalSIMPLEsubs) #To export it

dev.off()


# Figure with paired data, difficult sub-treatments
# Rename the figure legends
dataset16$Subtreatment = plyr::mapvalues(dataset16$Subtreatment, from = c("BS", "PL"), 
                                     to = c("BS","PL"))

# Reorder the experience levels (treatment variables)
dataset16$Phase <- factor(dataset16$Phase, levels = c("Initial","Test"))

# Reorder the experience levels (treatment variables)
dataset16$Subtreatment <- factor(dataset16$Subtreatment, levels = c("BS","PL"))

# Plot figure
FigurePetalDIFFsubs <- ggplot(dataset16, aes(x = Phase, y = Preference)) + 
  geom_boxplot(aes(fill = Phase), alpha = .2) +
  geom_line(aes(group = BeeID)) + 
  geom_point(size = 1) + 
  facet_wrap(~ Subtreatment) +
  ylab("Preference for Trained (S+) Petal Color") + xlab("Assay type") + theme_classic() + 
  theme(text=element_text(size=14)) +
  scale_color_viridis_d(name = "Experience \nlevel", end = 0.7, option = "inferno") + 
  scale_fill_viridis_d(name = "Experience \nlevel", end =0.7, option = 'inferno') +
  theme(legend.position  = c(-1.0, -1.0), legend.direction = "horizontal", 
        legend.background = element_rect(fill="transparent", colour = NA),
        legend.title = element_text(size = 0), legend.text = element_text(size = 0),
        panel.spacing.x = unit(1, "lines"), 
        plot.margin = margin(c(0,0, 0, 0), unit = "cm"))
plot(FigurePetalDIFFsubs) #To see the figure in R

# Set date
DATE <- "`r Sys.Date()`"

tiff(filename=paste("Fig_Initial-vs-Final-petal-Preference_DIFF-subs", DATE, ".tif", sep=""), units="mm", width=200, height=200, res=1200)

plot(FigurePetalDIFFsubs) #To export it

dev.off()


#### Plotting leaf shape preference changes by sub-treatment####

FILENAME17 <- "Analysis_Graph_Initial-vs-Final-leaf-preference-SIMPLE_2-18-2024.csv"
FILENAME18 <- "Analysis_Graph_Initial-vs-Final-leaf-preference-DIFF_2-18-2024.csv"
dataset17 <- read.csv("~/LL stats/Analysis_Graph_Initial-vs-Final-leaf-preference-SIMPLE_2-18-2024.csv", sep=",", header=TRUE)
dataset18 <- read.csv("~/LL stats/Analysis_Graph_Initial-vs-Final-leaf-preference-DIFF_2-18-2024.csv", sep=",", header=TRUE)

head(dataset17)
head(dataset18)

# Figure with paired data - simple subtreatments
# Rename the figure legends
dataset17$Subtreatment = plyr::mapvalues(dataset17$Subtreatment, from = c("BS", "YL"), 
                                     to = c("BS","YL"))

# Reorder the experience levels (treatment variables)
dataset17$Phase <- factor(dataset17$Phase, levels = c("Initial","Test"))

# Reorder the experience levels (treatment variables)
dataset17$Subtreatment <- factor(dataset17$Subtreatment, levels = c("BS","YL"))

# Plot figure
FigureLeafSIMPLEsubs <- ggplot(dataset17, aes(x = Phase, y = Preference)) + 
  geom_boxplot(aes(fill = Phase), alpha = .2) +
  geom_line(aes(group = BeeID)) + 
  geom_point(size = 1) + 
  facet_wrap(~ Subtreatment) +
  ylab("Preference for Trained (S+) Leaf Shape") + xlab("Assay type") + theme_classic() + 
  theme(text=element_text(size=14)) +
  scale_color_viridis_d(name = "Experience \nlevel", end = 0.7, option = "inferno") + 
  scale_fill_viridis_d(name = "Experience \nlevel", end =0.7, option = 'inferno') +
  theme(legend.position  = c(-1.0, -1.0), legend.direction = "horizontal", 
        legend.background = element_rect(fill="transparent", colour = NA),
        legend.title = element_text(size = 0), legend.text = element_text(size = 0),
        panel.spacing.x = unit(1, "lines"), 
        plot.margin = margin(c(0,0, 0, 0), unit = "cm"))
plot(FigureLeafSIMPLEsubs) #To see the figure in R

# Set date
DATE <- "`r Sys.Date()`"

tiff(filename=paste("Fig_Initial-vs-Final-leaf-Preference_SIMPLE-subs", DATE, ".tif", sep=""), units="mm", width=200, height=200, res=1200)

plot(FigureLeafSIMPLEsubs) #To export it

dev.off()

# Figure with paired data - difficult subtreatments
# Rename the figure legends
dataset18$Subtreatment = plyr::mapvalues(dataset18$Subtreatment, from = c("BS", "PL"), 
                                     to = c("BS","PL"))

# Reorder the experience levels (treatment variables)
dataset18$Phase <- factor(dataset18$Phase, levels = c("Initial","Test"))

# Reorder the experience levels (treatment variables)
dataset18$Subtreatment <- factor(dataset18$Subtreatment, levels = c("BS","PL"))

# Plot figure
FigureLeafDIFFsubs <- ggplot(dataset18, aes(x = Phase, y = Preference)) + 
  geom_boxplot(aes(fill = Phase), alpha = .2) +
  geom_line(aes(group = BeeID)) + 
  geom_point(size = 1) + 
  facet_wrap(~ Subtreatment) +
  ylab("Preference for Trained (S+) Leaf Shape") + xlab("Assay type") + theme_classic() + 
  theme(text=element_text(size=14)) +
  scale_color_viridis_d(name = "Experience \nlevel", end = 0.7, option = "inferno") + 
  scale_fill_viridis_d(name = "Experience \nlevel", end =0.7, option = 'inferno') +
  theme(legend.position  = c(-1.0, -1.0), legend.direction = "horizontal", 
        legend.background = element_rect(fill="transparent", colour = NA),
        legend.title = element_text(size = 0), legend.text = element_text(size = 0),
        panel.spacing.x = unit(1, "lines"), 
        plot.margin = margin(c(0,0, 0, 0), unit = "cm"))
plot(FigureLeafDIFFsubs) #To see the figure in R

# Set date
DATE <- "`r Sys.Date()`"

tiff(filename=paste("Fig_Initial-vs-Final-leaf-Preference_DIFF-subs", DATE, ".tif", sep=""), units="mm", width=200, height=200, res=1200)

plot(FigureLeafDIFFsubs) #To export it

dev.off()


```


##Bar plots

```{r}
#File first made on 7-21-2015
#Modified to fit leaf learning project on 12-26-22
# reorganized and modified to compare initial to test 12-21-23
# modified file locations to work in markdown 12-21-23
# MC rewrote plots in ggplot 12-22-23 

#LOAD EVERYTHING AFTER THIS

# library
library(ggplot2)
library(ggsignif)
library(dplyr)
library(ggpubr)
library(grid)

## FOR TRAINED BEES - INITIAL
## load data
#Name of data file:
FILENAME1 <- "STATS_BarplotCounts_Simp_InitialPhase-YL_12-21-23_MC.csv" # YL trained bees
FILENAME2 <- "STATS_BarplotCounts_Simp_InitialPhase-BS_12-21-23_MC.csv" # BS trained bees -simp
FILENAME3 <- "STATS_BarplotCounts_Diff_InitialPhase-PL_12-21-23_MC.csv" # PL trained bees
FILENAME4 <- "STATS_BarplotCounts_Diff_InitialPhase-BS_12-21-23_MC.csv" # BS trained bees -diff

## FOR TRAINED BEES - TEST
FILENAME5 <- "STATS_BarplotCounts_Simp_TestPhase-YL_12-21-23_MC.csv" # YL trained bees
FILENAME6 <- "STATS_BarplotCounts_Simp_TestPhase-BS_12-21-23_MC.csv" # BS trained bees -simp
FILENAME7 <- "STATS_BarplotCounts_Diff_TestPhase-PL_12-21-23_MC.csv" # PL trained bees
FILENAME8 <- "STATS_BarplotCounts_Diff_TestPhase-BS_12-21-23_MC.csv" # BS trained bees -diff

simp.pref.YL <- read.csv("~/Leaf Learning Stats/bayesian/barplot/STATS_BarplotCounts_Simp_InitialPhase-YL_12-21-23_MC.csv", sep=",", header=TRUE)
simp.pref.BS <- read.csv("~/Leaf Learning Stats/bayesian/barplot/STATS_BarplotCounts_Simp_InitialPhase-BS_12-21-23_MC.csv", sep=",", header=TRUE)
diff.pref.PL <- read.csv("~/Leaf Learning Stats/bayesian/barplot/STATS_BarplotCounts_Diff_InitialPhase-PL_12-21-23_MC.csv", sep=",", header=TRUE)
diff.pref.BS <- read.csv("~/Leaf Learning Stats/bayesian/barplot/STATS_BarplotCounts_Diff_InitialPhase-BS_12-21-23_MC.csv", sep=",", header=TRUE)

#Load data - test:
simp.test.YL <- read.csv("~/Leaf Learning Stats/bayesian/barplot/STATS_BarplotCounts_Simp_TestPhase-YL_12-21-23_MC.csv", sep=",", header=TRUE)
simp.test.BS <- read.csv("~/Leaf Learning Stats/bayesian/barplot/STATS_BarplotCounts_Simp_TestPhase-BS_12-21-23_MC.csv", sep=",", header=TRUE)
diff.test.PL <- read.csv("~/Leaf Learning Stats/bayesian/barplot/STATS_BarplotCounts_Diff_TestPhase-PL_12-21-23_MC.csv", sep=",", header=TRUE)
diff.test.BS <- read.csv("~/Leaf Learning Stats/bayesian/barplot/STATS_BarplotCounts_Diff_TestPhase-BS_12-21-23_MC.csv", sep=",", header=TRUE)

# Look at data #
# INITIAL
head(simp.pref.YL)
head(simp.pref.BS)
head(diff.pref.PL)
head(diff.pref.BS)
# TEST
head(simp.test.YL)
head(simp.test.BS)
head(diff.test.PL)
head(diff.test.BS)

#### GRAPH OF SIMPLE INITIAL AND TEST PHASE, BEES TRAINED TO YL VISIT PERCENTAGES FOR ALL FOUR TYPES OF TARGETS ####

# Calculate means and standard errors for each treatment
#   Means
means.yl.s <- c(mean(simp.pref.YL$VisitPercent[simp.pref.YL$Target == "BL"]), mean(simp.test.YL$VisitPercent[simp.test.YL$Target == "BL"]),
           mean(simp.pref.YL$VisitPercent[simp.pref.YL$Target == "BS"]), mean(simp.test.YL$VisitPercent[simp.test.YL$Target == "BS"]),
           mean(simp.pref.YL$VisitPercent[simp.pref.YL$Target == "YS"]), mean(simp.test.YL$VisitPercent[simp.test.YL$Target == "YL"]),
           mean(simp.pref.YL$VisitPercent[simp.pref.YL$Target == "YL"]), mean(simp.test.YL$VisitPercent[simp.test.YL$Target == "YS"]))

#   SEs
sds.yl.s <- c(sd(simp.pref.YL$VisitPercent[simp.pref.YL$Target == "BL"]), sd(simp.test.YL$VisitPercent[simp.test.YL$Target == "BL"]), 
         sd(simp.pref.YL$VisitPercent[simp.pref.YL$Target == "BS"]), sd(simp.test.YL$VisitPercent[simp.test.YL$Target == "BS"]), 
         sd(simp.pref.YL$VisitPercent[simp.pref.YL$Target == "YL"]), sd(simp.test.YL$VisitPercent[simp.test.YL$Target == "YL"]),
         sd(simp.pref.YL$VisitPercent[simp.pref.YL$Target == "YS"]), sd(simp.test.YL$VisitPercent[simp.test.YL$Target == "YS"]))
n.yl.s <- c(length(simp.pref.YL$VisitPercent[simp.pref.YL$Target == "BL"]), length(simp.test.YL$VisitPercent[simp.test.YL$Target == "BL"]), 
       length(simp.pref.YL$VisitPercent[simp.pref.YL$Target == "BS"]), length(simp.test.YL$VisitPercent[simp.test.YL$Target == "BS"]),
       length(simp.pref.YL$VisitPercent[simp.pref.YL$Target == "YL"]), length(simp.test.YL$VisitPercent[simp.test.YL$Target == "YL"]),
       length(simp.pref.YL$VisitPercent[simp.pref.YL$Target == "YS"]), length(simp.test.YL$VisitPercent[simp.test.YL$Target == "YS"]))
ses.yl.s <- sds.yl.s/sqrt(n.yl.s)

# Make the plot

Flower_type.s <- c(rep("BL" , 2) , rep("BS" , 2) , rep("YL" , 2) , rep("YS" , 2))
Phase <- rep(c("Initial" , "Test") , 4)
visits.yl.s <- means.yl.s
errorbars.yl.s <- ses.yl.s

df.yl.s <- data.frame(Flower_type.s, Phase, visits.yl.s, errorbars.yl.s)  
df.yl.s$significant <- dplyr::case_when(
  grepl("BL", df.yl.s$Flower_type.s) & grepl("Initial", df$Phase) ~ TRUE, 
  grepl("BS", df.yl.s$Flower_type.s) & grepl("Initial", df$Phase) ~ TRUE,
  grepl("YL", df.yl.s$Flower_type.s) & grepl("Test", df$Phase) ~ FALSE,
  grepl("YS", df.yl.s$Flower_type.s) & grepl("Test", df$Phase) ~ TRUE,
  TRUE ~ FALSE
)

plot.yl.s <- ggplot(df.yl.s, aes(y=visits.yl.s, x=Flower_type.s, fill = Phase)) +
  geom_hline(yintercept=25, linetype="dashed", color="red", size=0.5) +
  geom_col( position="dodge", width=0.5, alpha = 0.7, color = "black", size = 0.1) +
  geom_errorbar(aes(ymin = visits.yl.s-ses.yl.s, ymax = visits.yl.s+ses.yl.s),
                position =  position_dodge(width = 0.5), width = 0.2) +
  geom_text(aes(label = ifelse(significant, "*", ""), group = Flower_type.s), 
            position = position_dodge(width = .9), vjust = -1.7, size = 6) +
  geom_text(aes(label = ifelse(significant, "___", ""), group = Flower_type.s), 
            position = position_dodge(width = .9), vjust = -2.2, size = 6) +
  scale_fill_manual(values = c("yellow3","yellow4")) +
  scale_y_continuous(expand=expansion(0),
                     limits = c(0,60)) +
  labs(title = "c", x = "Flower type", y = "Percent of visits (mean + SE)") 

plot(plot.yl.s)


#### GRAPH OF SIMPLE INITIAL AND TEST PHASE, BEES TRAINED TO BS VISIT PERCENTAGES FOR ALL FOUR TYPES OF TARGETS ####

# Calculate means and standard errors for each treatment
#   Means
means.bs.s <- c(mean(simp.pref.BS$VisitPercent[simp.pref.BS$Target == "BL"]), mean(simp.test.BS$VisitPercent[simp.test.BS$Target == "BL"]),
           mean(simp.pref.BS$VisitPercent[simp.pref.BS$Target == "BS"]), mean(simp.test.BS$VisitPercent[simp.test.BS$Target == "BS"]),
           mean(simp.pref.BS$VisitPercent[simp.pref.BS$Target == "YS"]), mean(simp.test.BS$VisitPercent[simp.test.BS$Target == "YL"]),
           mean(simp.pref.BS$VisitPercent[simp.pref.BS$Target == "YL"]), mean(simp.test.BS$VisitPercent[simp.test.BS$Target == "YS"]))

#   SEs
sds.bs.s <- c(sd(simp.pref.BS$VisitPercent[simp.pref.BS$Target == "BL"]), sd(simp.test.BS$VisitPercent[simp.test.BS$Target == "BL"]), 
         sd(simp.pref.BS$VisitPercent[simp.pref.BS$Target == "BS"]), sd(simp.test.BS$VisitPercent[simp.test.BS$Target == "BS"]), 
         sd(simp.pref.BS$VisitPercent[simp.pref.BS$Target == "YL"]), sd(simp.test.BS$VisitPercent[simp.test.BS$Target == "YL"]),
         sd(simp.pref.BS$VisitPercent[simp.pref.BS$Target == "YS"]), sd(simp.test.BS$VisitPercent[simp.test.BS$Target == "YS"]))
n.bs.s <- c(length(simp.pref.BS$VisitPercent[simp.pref.BS$Target == "BL"]), length(simp.test.BS$VisitPercent[simp.test.BS$Target == "BL"]), 
       length(simp.pref.BS$VisitPercent[simp.pref.BS$Target == "BS"]), length(simp.test.BS$VisitPercent[simp.test.BS$Target == "BS"]),
       length(simp.pref.BS$VisitPercent[simp.pref.BS$Target == "YL"]), length(simp.test.BS$VisitPercent[simp.test.BS$Target == "YL"]),
       length(simp.pref.BS$VisitPercent[simp.pref.BS$Target == "YS"]), length(simp.test.BS$VisitPercent[simp.test.BS$Target == "YS"]))
ses.bs.s <- sds.bs.s/sqrt(n.bs.s)

Flower_type.s <- c(rep("BL" , 2) , rep("BS" , 2) , rep("YL" , 2) , rep("YS" , 2))
Phase <- rep(c("Initial" , "Test") , 4)
visits.bs.s <- means.bs.s
errorbars.bs.s <- ses.bs.s

df.bs.s <- data.frame(Flower_type.s, Phase, visits.bs.s, errorbars.bs.s)  
df.bs.s$significant <- dplyr::case_when(
  grepl("BL", df.bs.s$Flower_type) ~ FALSE,
  grepl("BS", df.bs.s$Flower_type) & grepl("Test", df$Phase) ~ TRUE,
  grepl("YL", df.bs.s$Flower_type) & grepl("Initial", df$Phase) ~ FALSE,
  grepl("YS", df.bs.s$Flower_type) ~ FALSE,
  TRUE ~ FALSE
)

plot.bs.s <- ggplot(df.bs.s, aes(y=visits.bs.s, x=Flower_type.s, fill = Phase)) +
  geom_hline(yintercept=25, linetype="dashed", color="red", size=0.5) +
  geom_col( position="dodge", width=0.5, alpha = 0.7, color = "black", size = 0.1) +
  geom_errorbar(aes(ymin = visits.bs.s-ses.bs.s, ymax = visits.bs.s+ses.bs.s),
                position =  position_dodge(width = 0.5), width = 0.2) +
  geom_text(aes(label = ifelse(significant, "*", ""), group = Flower_type.s), 
            position = position_dodge(width = .9), vjust = -1.5, size = 18 / .pt) +
  geom_text(aes(label = ifelse(significant, "___", ""), group = Flower_type.s), 
            position = position_dodge(width = .9), vjust = -2.2, size = 6) +
  scale_fill_manual(values = c("deepskyblue2","deepskyblue4")) +
  scale_y_continuous(expand=expansion(0),
                     limits = c(0,60)) +
  labs(title = "a", x = "Flower type", y = "Percent of visits (mean + SE)") 

plot(plot.bs.s)

#### GRAPH OF DIFFICULT INITIAL AND TEST PHASE, BEES TRAINED TO PL VISIT PERCENTAGES FOR ALL FOUR TYPES OF TARGETS ####

# Calculate means and standard errors for each treatment
#   Means
means.pl.d <- c(mean(diff.pref.PL$VisitPercent[diff.pref.PL$Target == "BL"]), mean(diff.test.PL$VisitPercent[diff.test.PL$Target == "BL"]),
           mean(diff.pref.PL$VisitPercent[diff.pref.PL$Target == "BS"]), mean(diff.test.PL$VisitPercent[diff.test.PL$Target == "BS"]),
           mean(diff.pref.PL$VisitPercent[diff.pref.PL$Target == "PS"]), mean(diff.test.PL$VisitPercent[diff.test.PL$Target == "PL"]),
           mean(diff.pref.PL$VisitPercent[diff.pref.PL$Target == "PL"]), mean(diff.test.PL$VisitPercent[diff.test.PL$Target == "PS"]))

#   SEs
sds.pl.d <- c(sd(diff.pref.PL$VisitPercent[diff.pref.PL$Target == "BL"]), sd(diff.test.PL$VisitPercent[diff.test.PL$Target == "BL"]), 
         sd(diff.pref.PL$VisitPercent[diff.pref.PL$Target == "BS"]), sd(diff.test.PL$VisitPercent[diff.test.PL$Target == "BS"]), 
         sd(diff.pref.PL$VisitPercent[diff.pref.PL$Target == "PL"]), sd(diff.test.PL$VisitPercent[diff.test.PL$Target == "PL"]),
         sd(diff.pref.PL$VisitPercent[diff.pref.PL$Target == "PS"]), sd(diff.test.PL$VisitPercent[diff.test.PL$Target == "PS"]))
n.pl.d <- c(length(diff.pref.PL$VisitPercent[diff.pref.PL$Target == "BL"]), length(diff.test.PL$VisitPercent[diff.test.PL$Target == "BL"]), 
       length(diff.pref.PL$VisitPercent[diff.pref.PL$Target == "BS"]), length(diff.test.PL$VisitPercent[diff.test.PL$Target == "BS"]),
       length(diff.pref.PL$VisitPercent[diff.pref.PL$Target == "PL"]), length(diff.test.PL$VisitPercent[diff.test.PL$Target == "PL"]),
       length(diff.pref.PL$VisitPercent[diff.pref.PL$Target == "PS"]), length(diff.test.PL$VisitPercent[diff.test.PL$Target == "PS"]))
ses.pl.d <- sds.pl.d/sqrt(n.pl.d)

Flower_type.d <- c(rep("BL" , 2) , rep("BS" , 2) , rep("PL" , 2) , rep("PS" , 2))
Phase <- rep(c("Initial" , "Test") , 4)
visits.pl.d <- means.pl.d
errorbars.pl.d <- ses.pl.d

df.pl.d <- data.frame(Flower_type.d, Phase, visits.pl.d, errorbars.pl.d)  
df.pl.d$significant <- dplyr::case_when(
  grepl("BL", df.pl.d$Flower_type) ~ FALSE,
  grepl("BS", df.pl.d$Flower_type) ~ FALSE,
  grepl("PL", df.pl.d$Flower_type) ~ FALSE,
  grepl("PS", df.pl.d$Flower_type) ~ FALSE,
  TRUE ~ FALSE
)

plot.pl.d <- ggplot(df.pl.d, aes(y=visits.pl.d, x=Flower_type.d, fill = Phase)) +
  geom_hline(yintercept=25, linetype="dashed", color="red", size=0.5) +
  geom_col( position="dodge", width=0.5, alpha = 0.7, color = "black", size = 0.1) +
  geom_errorbar(aes(ymin = visits.pl.d-ses.pl.d, ymax = visits.pl.d+ses.pl.d),
                position =  position_dodge(width = 0.5), width = 0.2) +
  geom_text(aes(label = ifelse(significant, "*", ""), group = Flower_type.d), 
            position = position_dodge(width = .9), vjust = -1.5, size = 18 / .pt) +
  geom_text(aes(label = ifelse(significant, "___", ""), group = Flower_type.d), 
            position = position_dodge(width = .9), vjust = -2.2, size = 6) +
  scale_fill_manual(values = c("darkorchid3","darkorchid4")) +
  scale_y_continuous(expand=expansion(0),
                     limits = c(0,60)) +
  labs(title = "d", x = "Flower type", y = "Percent of visits (mean + SE)") 

plot(plot.pl.d)


#### GRAPH OF DIFFICULT INITIAL AND TEST PHASE, BEES TRAINED TO BS VISIT PERCENTAGES FOR ALL FOUR TYPES OF TARGETS ####

# Calculate means and standard errors for each treatment
#   Means
means.bs.d <- c(mean(diff.pref.BS$VisitPercent[diff.pref.BS$Target == "BL"]), mean(diff.test.BS$VisitPercent[diff.test.BS$Target == "BL"]),
           mean(diff.pref.BS$VisitPercent[diff.pref.BS$Target == "BS"]), mean(diff.test.BS$VisitPercent[diff.test.BS$Target == "BS"]),
           mean(diff.pref.BS$VisitPercent[diff.pref.BS$Target == "PS"]), mean(diff.test.BS$VisitPercent[diff.test.BS$Target == "PL"]),
           mean(diff.pref.BS$VisitPercent[diff.pref.BS$Target == "PL"]), mean(diff.test.BS$VisitPercent[diff.test.BS$Target == "PS"]))

#   SEs
sds.bs.d <- c(sd(diff.pref.BS$VisitPercent[diff.pref.BS$Target == "BL"]), sd(diff.test.BS$VisitPercent[diff.test.BS$Target == "BL"]), 
         sd(diff.pref.BS$VisitPercent[diff.pref.BS$Target == "BS"]), sd(diff.test.BS$VisitPercent[diff.test.BS$Target == "BS"]), 
         sd(diff.pref.BS$VisitPercent[diff.pref.BS$Target == "PL"]), sd(diff.test.BS$VisitPercent[diff.test.BS$Target == "PL"]),
         sd(diff.pref.BS$VisitPercent[diff.pref.BS$Target == "PS"]), sd(diff.test.BS$VisitPercent[diff.test.BS$Target == "PS"]))
n.bs.d <- c(length(diff.pref.BS$VisitPercent[diff.pref.BS$Target == "BL"]), length(diff.test.BS$VisitPercent[diff.test.BS$Target == "BL"]), 
       length(diff.pref.BS$VisitPercent[diff.pref.BS$Target == "BS"]), length(diff.test.BS$VisitPercent[diff.test.BS$Target == "BS"]),
       length(diff.pref.BS$VisitPercent[diff.pref.BS$Target == "PL"]), length(diff.test.BS$VisitPercent[diff.test.BS$Target == "PL"]),
       length(diff.pref.BS$VisitPercent[diff.pref.BS$Target == "PS"]), length(diff.test.BS$VisitPercent[diff.test.BS$Target == "PS"]))
ses.bs.d <- sds.bs.d/sqrt(n.bs.d)

Flower_type.d <- c(rep("BL" , 2) , rep("BS" , 2) , rep("PL" , 2) , rep("PS" , 2))
Phase <- rep(c("Initial" , "Test") , 4)
visits.bs.d <- means.bs.d
errorbars.bs.d <- ses.bs.d

df.bs.d <- data.frame(Flower_type.d, Phase, visits.bs.d, errorbars.bs.d)  
df.bs.d$significant <- dplyr::case_when(
  grepl("BL", df.bs.d$Flower_type) & grepl("Test", df$Phase) ~ FALSE,
  grepl("BS", df.bs.d$Flower_type) ~ FALSE,
  grepl("PL", df.bs.d$Flower_type) & grepl("Initial", df$Phase) ~ FALSE,
  grepl("PS", df.bs.d$Flower_type) ~ FALSE,
  TRUE ~ FALSE
)

plot.bs.d <- ggplot(df.bs.d, aes(y=visits.bs.d, x=Flower_type.d, fill = Phase)) +
  geom_hline(yintercept=25, linetype="dashed", color="red", size=0.5) +
  geom_col( position="dodge", width=0.5, alpha = 0.7, color = "black", size = 0.1) +
  geom_errorbar(aes(ymin = visits.bs.d-ses.bs.d, ymax = visits.bs.d+ses.bs.d),
                position =  position_dodge(width = 0.5), width = 0.2) +
  geom_text(aes(label = ifelse(significant, "*", " "), group = Flower_type.d), 
            position = position_dodge(width = .9), vjust = -1.5, size = 18 / .pt) +
  geom_text(aes(label = ifelse(significant, "___", ""), group = Flower_type.d), 
            position = position_dodge(width = .9), vjust = -2.2, size = 6) +
  scale_fill_manual(values = c("deepskyblue2","deepskyblue4")) +
  scale_y_continuous(expand=expansion(0),
                     limits = c(0,60)) + 
  labs(title = "b", x = "Flower type", y = "Percent of visits (mean + SE)") 

plot(plot.bs.d)

#### Plotting all together ####

combinedfigure <- ggarrange(plot.bs.s, plot.bs.d, plot.yl.s, plot.pl.d,  
                    plot.bs.s + rremove("xlab"), 
                    plot.bs.d + rremove("ylab") + rremove("xlab"), 
                    plot.yl.s, 
                    plot.pl.d + rremove("ylab"), 
                    font.label = list(size = 14, color = "black", face = "bold", family = NULL, position = "right"),
                    labels = c("                  Simple", 
                               "                 Difficult"), 
                    ncol = 2, nrow = 2)
combinedfigure

DATE <- "`r Sys.Date()`"

tiff(filename=paste("Fig_CombinedBarplots_", DATE, ".tif", sep=""), units="mm", width=250, height=200, res=1200)

combinedfigure

dev.off()


#### Replotting to show the simple sub-treatments as one and the difficult sub-treatments as one ####

# library
library(ggplot2)
library(ggsignif)
library(dplyr)
library(ggpubr)
library(grid)


## load data
#Name of data file:
FILENAME9 <- "STATS_BarplotCounts_Simp_InitialPhase_2-15-24_MC.csv" # simp bees initial
FILENAME10 <- "STATS_BarplotCounts_Simp_TestPhase_2-15-24_MC.csv" # simp test
FILENAME11 <- "STATS_BarplotCounts_Diff_InitialPhase_2-15-24_MC.csv" # diff bees initial
FILENAME12 <- "STATS_BarplotCounts_Diff_TestPhase_2-15-24_MC.csv" # diff test

simp.pref <- read.csv("~/LL stats/STATS_BarplotCounts_Simp_InitialPhase_2-15-24_MC.csv", sep=",", header=TRUE)
simp.test <- read.csv("~/LL stats/STATS_BarplotCounts_Simp_TestPhase_2-15-24_MC.csv", sep=",", header=TRUE)
diff.pref <- read.csv("~/LL stats/STATS_BarplotCounts_Diff_InitialPhase_2-15-24_MC.csv", sep=",", header=TRUE)
diff.test <- read.csv("~/LL stats/STATS_BarplotCounts_Diff_TestPhase_2-15-24_MC.csv", sep=",", header=TRUE)

# Look at data #
head(simp.pref)
head(simp.test)
head(diff.pref)
head(diff.test)


#### GRAPH OF SIMPLE INITIAL AND TEST PHASE ####

# Calculate means and standard errors for each treatment
#   Means
means.s <- c(mean(simp.pref$VisitPercent[simp.pref$Target == "CPIL"]), mean(simp.test$VisitPercent[simp.test$Target == "CPIL"]),
           mean(simp.pref$VisitPercent[simp.pref$Target == "CPCL"]), mean(simp.test$VisitPercent[simp.test$Target == "CPCL"]),
           mean(simp.pref$VisitPercent[simp.pref$Target == "IPIL"]), mean(simp.test$VisitPercent[simp.test$Target == "IPIL"]),
           mean(simp.pref$VisitPercent[simp.pref$Target == "IPCL"]), mean(simp.test$VisitPercent[simp.test$Target == "IPCL"]))

#   SEs
sds.s <- c(sd(simp.pref$VisitPercent[simp.pref$Target == "CPIL"]), sd(simp.test$VisitPercent[simp.test$Target == "CPIL"]), 
         sd(simp.pref$VisitPercent[simp.pref$Target == "CPCL"]), sd(simp.test$VisitPercent[simp.test$Target == "CPCL"]), 
         sd(simp.pref$VisitPercent[simp.pref$Target == "IPIL"]), sd(simp.test$VisitPercent[simp.test$Target == "IPIL"]),
         sd(simp.pref$VisitPercent[simp.pref$Target == "IPCL"]), sd(simp.test$VisitPercent[simp.test$Target == "IPCL"]))

n.s <- c(length(simp.pref$VisitPercent[simp.pref$Target == "CPIL"]), length(simp.test$VisitPercent[simp.test$Target == "CPIL"]), 
       length(simp.pref$VisitPercent[simp.pref$Target == "CPCL"]), length(simp.test$VisitPercent[simp.test$Target == "CPCL"]),
       length(simp.pref$VisitPercent[simp.pref$Target == "IPIL"]), length(simp.test$VisitPercent[simp.test$Target == "IPIL"]),
       length(simp.pref$VisitPercent[simp.pref$Target == "IPCL"]), length(simp.test$VisitPercent[simp.test$Target == "IPCL"]))
ses.s <- sds.s/sqrt(n.s)

Flower_type.s <- c(rep("CPIL" , 2) , rep("CPCL" , 2) , rep("IPIL" , 2) , rep("IPCL" , 2))
Phase <- rep(c("Initial" , "Test") , 4)
visits.s <- means.s
errorbars.s <- ses.s

df.s <- data.frame(Flower_type.s, Phase, visits.s, errorbars.s)  
df.s$significant <- dplyr::case_when(
  grepl("CPIL", df.s$Flower_type) ~ FALSE,
  grepl("CPCL", df.s$Flower_type) & grepl("Test", df.s$Phase) ~ TRUE,
  grepl("IPIL", df.s$Flower_type) & grepl("Initial", df.s$Phase) ~ TRUE,
  grepl("IPCL", df.s$Flower_type) & grepl("Initial", df.s$Phase) ~ TRUE,
  TRUE ~ FALSE
)

plot.s <- ggplot(df.s, aes(y=visits.s, x=Flower_type.s, fill = Phase)) +
  geom_hline(yintercept=25, linetype="dashed", color="red", size=0.5) +
  geom_col( position="dodge", width=0.5, alpha = 0.7, color = "black", size = 0.1) +
  geom_errorbar(aes(ymin = visits.s-ses.s, ymax = visits.s+ses.s),
                position =  position_dodge(width = 0.5), width = 0.2) +
  geom_text(aes(label = ifelse(significant, "*", ""), group = Flower_type.s), 
            position = position_dodge(width = .9), vjust = -1.7, size = 18 / .pt) +
  geom_text(aes(label = ifelse(significant, "___", ""), group = Flower_type.s), 
            position = position_dodge(width = .9), vjust = -2.4, size = 6) +
  scale_fill_manual(values = c("yellow3","yellow4")) +
  scale_y_continuous(expand=expansion(0),
                     limits = c(0,60)) +
  labs(title = "a", x = "Flower type", y = "Percent of visits (mean + SE)") 

plot(plot.s)

#### GRAPH OF DIFFICULT INITIAL AND TEST PHASE, BEES TRAINED TO PL VISIT PERCENTAGES FOR ALL FOUR TYPES OF TARGETS ####

# Calculate means and standard errors for each treatment
#   Means
means.d <- c(mean(diff.pref$VisitPercent[diff.pref$Target == "CPIL"]), mean(diff.test$VisitPercent[diff.test$Target == "CPIL"]),
           mean(diff.pref$VisitPercent[diff.pref$Target == "CPCL"]), mean(diff.test$VisitPercent[diff.test$Target == "CPCL"]),
           mean(diff.pref$VisitPercent[diff.pref$Target == "IPIL"]), mean(diff.test$VisitPercent[diff.test$Target == "IPIL"]),
           mean(diff.pref$VisitPercent[diff.pref$Target == "IPCL"]), mean(diff.test$VisitPercent[diff.test$Target == "IPCL"]))

#   SEs
sds.d <- c(sd(diff.pref$VisitPercent[diff.pref$Target == "CPIL"]), sd(diff.test$VisitPercent[diff.test$Target == "CPIL"]), 
         sd(diff.pref$VisitPercent[diff.pref$Target == "CPCL"]), sd(diff.test$VisitPercent[diff.test$Target == "CPCL"]), 
         sd(diff.pref$VisitPercent[diff.pref$Target == "IPIL"]), sd(diff.test$VisitPercent[diff.test$Target == "IPIL"]),
         sd(diff.pref$VisitPercent[diff.pref$Target == "IPCL"]), sd(diff.test$VisitPercent[diff.test$Target == "IPCL"]))

n.d <- c(length(diff.pref$VisitPercent[diff.pref$Target == "CPIL"]), length(diff.test$VisitPercent[diff.test$Target == "CPIL"]), 
       length(diff.pref$VisitPercent[diff.pref$Target == "CPCL"]), length(diff.test$VisitPercent[diff.test$Target == "CPCL"]),
       length(diff.pref$VisitPercent[diff.pref$Target == "IPIL"]), length(diff.test$VisitPercent[diff.test$Target == "IPIL"]),
       length(diff.pref$VisitPercent[diff.pref$Target == "IPCL"]), length(diff.test$VisitPercent[diff.test$Target == "IPCL"]))
ses.d <- sds.d/sqrt(n.d)

Flower_type.d <- c(rep("CPIL" , 2) , rep("CPCL" , 2) , rep("IPIL" , 2) , rep("IPCL" , 2))
Phase <- rep(c("Initial" , "Test") , 4)
visits.d <- means.d
errorbars.d <- ses.d

df.d <- data.frame(Flower_type.d, Phase, visits.d, errorbars.d)  
df.d$significant <- dplyr::case_when(
  grepl("CPIL", df.d$Flower_type) ~ FALSE,
  grepl("CPCL", df.d$Flower_type) & grepl("Test", df.d$Phase) ~ FALSE,
  grepl("IPIL", df.d$Flower_type) & grepl("Initial", df.d$Phase) ~ FALSE,
  grepl("IPCL", df.d$Flower_type) & grepl("Initial", df.d$Phase) ~ FALSE,
  TRUE ~ FALSE
)

plot.d <- ggplot(df.d, aes(y=visits.d, x=Flower_type.d, fill = Phase)) +
  geom_hline(yintercept=25, linetype="dashed", color="red", size=0.5) +
  geom_col( position="dodge", width=0.5, alpha = 0.7, color = "black", size = 0.1) +
  geom_errorbar(aes(ymin = visits.d-ses.d, ymax = visits.d+ses.d),
                position =  position_dodge(width = 0.5), width = 0.2) +
  geom_text(aes(label = ifelse(significant, "*", ""), group = Flower_type.d), 
            position = position_dodge(width = .9), vjust = -1.7, size = 18 / .pt) +
  geom_text(aes(label = ifelse(significant, "___", ""), group = Flower_type.d), 
            position = position_dodge(width = .9), vjust = -2.4, size = 6) +
  scale_fill_manual(values = c("darkorchid3","darkorchid4")) +
  scale_y_continuous(expand=expansion(0),
                     limits = c(0,60)) +
  labs(title = "b", x = "Flower type", y = "Percent of visits (mean + SE)") 

plot(plot.d)


#### Plotting simp and diff together ####

combinedfigure2 <- ggarrange(plot.s, plot.d,  
                    plot.s, 
                    plot.d + rremove("ylab"), 
                    font.label = list(size = 14, color = "black", face = "bold", family = NULL, position = "right"),
                    labels = c("               Simple", 
                               "               Difficult"), 
                    ncol = 2, nrow = 1)
combinedfigure2

DATE <- "`r Sys.Date()`"

tiff(filename=paste("Fig_CombinedBarplots_", DATE, ".tif", sep=""), units="mm", width=220, height=100, res=3000)

combinedfigure2

dev.off()


```


##BAYESIAN STUFF

Bayesian analysis of how preferences for each combinatory cue changed from the initial to test phase for each treatment, split by subtreatment.

```{r}
# Changelog
# 12-6-2022 original file creation and adaption to leaf learning data
# 12-16-2022 fixed some errors
# 12-19-2023 Reviewed by MC and ALR
# 12-20-2023 Complete reorganization and revision by MC
# 12-21-2023 Ran bayesian, results recorded in R, and moved code to markdown doc

#### SETUP ####
#Load the library that bayespref() is in:
library(bayespref)

#### IMPORT ALL REQUIRED FILES ####
## Example with one population (named "1") and the four choices.
## Note that populations do need to be numbered.

## FOR TRAINED BEES - INITIAL
## load data
#Name of data file:
FILENAME1 <- "STATS_Counts_Simp_InitialPhaseAnalysis-YL_12-19-23_MC.csv" # YL trained bees
FILENAME2 <- "STATS_Counts_Simp_InitialPhaseAnalysis-BS_12-19-23_MC.csv" # BS trained bees -simp
FILENAME3 <- "STATS_Counts_Diff_InitialPhaseAnalysis-PL_12-20-23_MC.csv" # PL trained bees
FILENAME4 <- "STATS_Counts_Diff_InitialPhaseAnalysis-BS_12-20-23_MC.csv" # BS trained bees -diff

## FOR TRAINED BEES - TEST
FILENAME5 <- "STATS_Counts_Simp_TestPhaseAnalysis-YL_12-20-23_MC.csv" # YL trained bees
FILENAME6 <- "STATS_Counts_Simp_TestPhaseAnalysis-BS_12-20-23_MC.csv" # BS trained bees -simp
FILENAME7 <- "STATS_Counts_Diff_TestPhaseAnalysis-PL_12-19-23_MC.csv" # PL trained bees
FILENAME8 <- "STATS_Counts_Diff_TestPhaseAnalysis-BS_12-19-23_MC.csv" # BS trained bees -diff

## FOR SUB-TREATMENTS COMBINED - INITIAL AND TEST
FILENAME9 <- "STATS_Counts_Diff_InitialPhaseAnalysis_2-14-24_MC.csv" #difficult initial
FILENAME10 <- "STATS_Counts_Diff_TestPhaseAnalysis_2-14-24_MC.csv" #difficult test
FILENAME11 <- "STATS_Counts_Simp_InitialPhaseAnalysis_2-14-24_MC.csv" #simple initial
FILENAME12 <- "STATS_Counts_Simp_TestPhaseAnalysis_2-14-24_MC.csv" #simple test

#Load data - initial:
simp.pref.YL <- read.csv("~/Leaf Learning Stats/bayesian/STATS_Counts_Simp_InitialPhaseAnalysis-YL_12-19-23_MC.csv", sep=",", header=TRUE)
simp.pref.BS <- read.csv("~/Leaf Learning Stats/bayesian/STATS_Counts_Simp_InitialPhaseAnalysis-BS_12-19-23_MC.csv", sep=",", header=TRUE)
diff.pref.PL <- read.csv("~/Leaf Learning Stats/bayesian/STATS_Counts_Diff_InitialPhaseAnalysis-PL_12-20-23_MC.csv", sep=",", header=TRUE)
diff.pref.BS <- read.csv("~/Leaf Learning Stats/bayesian/STATS_Counts_Diff_InitialPhaseAnalysis-BS_12-20-23_MC.csv", sep=",", header=TRUE)

#Load data - test:
simp.test.YL <- read.csv("~/Leaf Learning Stats/bayesian/STATS_Counts_Simp_TestPhaseAnalysis-YL_12-20-23_MC.csv", sep=",", header=TRUE)
simp.test.BS <- read.csv("~/Leaf Learning Stats/bayesian/STATS_Counts_Simp_TestPhaseAnalysis-BS_12-20-23_MC.csv", sep=",", header=TRUE)
diff.test.PL <- read.csv("~/Leaf Learning Stats/bayesian/STATS_Counts_Diff_TestPhaseAnalysis-PL_12-19-23_MC.csv", sep=",", header=TRUE)
diff.test.BS <- read.csv("~/Leaf Learning Stats/bayesian/STATS_Counts_Diff_TestPhaseAnalysis-BS_12-19-23_MC.csv", sep=",", header=TRUE)

#load data - no sub-treatments
## FOR SUB-TREATMENTS COMBINED - INITIAL AND TEST
diffpref <- read.csv("~/LL stats/STATS_Counts_Diff_InitialPhaseAnalysis_2-14-24_MC.csv", sep=",", header=TRUE,check.names=FALSE) 
difftest <- read.csv("~/LL stats/STATS_Counts_Diff_TestPhaseAnalysis_2-14-24_MC.csv", sep=",", header=TRUE,check.names=FALSE)
simppref <- read.csv("~/LL stats/STATS_Counts_Simp_InitialPhaseAnalysis_2-14-24_MC.csv", sep=",", header=TRUE,check.names=FALSE)
simptest <- read.csv("~/LL stats/STATS_Counts_Simp_TestPhaseAnalysis_2-14-24_MC.csv", sep=",", header=TRUE,check.names=FALSE)

# Look at data #
# INITIAL
head(simp.pref.YL)
head(simp.pref.BS)
head(diff.pref.PL)
head(diff.pref.BS)
# TEST
head(simp.test.YL)
head(simp.test.BS)
head(diff.test.PL)
head(diff.test.BS)
# W/O SUB-TREATMENTS
head(diffpref)
head(difftest)
head(simppref)
head(simptest)

#### FOR SIMPLE PETAL CHOICE TRAINED BEES - INITIAL BS ####
FILENAME2 <- "STATS_Counts_Simp_InitialPhaseAnalysis-BS_12-19-23_MC.csv" 
head(simp.pref.BS)

## all bees in one group ('pop')
data3 <- subset(simp.pref.BS,Pop==1,select=-c(BeeID))
data3 <- droplevels(data3)
data3 <- as.matrix(data3)
head(data3)

# run the model
SimpPrefBS <- bayesPref(pData=data3,mcmcL=40000,pops=TRUE,dicburn=10000,dirvar=9)
#dic =  61.09866 

# examine mixing, which is not great at 1000 reps
plot(SimpPrefBS[[1]]$PopPref[1,],xlab="MCMC step",ylab="PopPref",cex=0.5)

## simple plot
prefPlot(SimpPrefBS[[1]],ymax=35)

## means and credible intervals
## this next command will get you means and credible intervals returned in the $PopPref element
## the means are in the middle column, and the credible intervals are on either side
credibleIntervals(prefres = SimpPrefBS[[1]],burn=10000,interval=0.95)  

#$IndPref
#, , 1
#
#[,1]      [,2]      [,3]      [,4]
#[1,] 0.1490548 0.1084565 0.1945512 0.1921895
#[2,] 0.1555752 0.1048530 0.1236695 0.2124786
#[3,] 0.1942530 0.1039827 0.1661182 0.1526793
#[4,] 0.1187521 0.1312272 0.1635993 0.1754813
#[5,] 0.1559317 0.1659385 0.1199347 0.1918012
#[6,] 0.1485870 0.1287431 0.1529351 0.2129212
#[7,] 0.1437734 0.1520178 0.1904558 0.1476190
#[8,] 0.1347610 0.1143463 0.2036483 0.1810794
#[9,] 0.1561448 0.1437846 0.1495492 0.1585336
#
#, , 2
#
#[,1]      [,2]      [,3]      [,4]
#[1,] 0.2337827 0.1849374 0.2871228 0.2855904
#[2,] 0.2552377 0.1929908 0.2177929 0.3225767
#[3,] 0.2945099 0.1858509 0.2626761 0.2477275
#[4,] 0.2126439 0.2259654 0.2678241 0.2818976
#[5,] 0.2444461 0.2563175 0.2021135 0.2880107
#[6,] 0.2343391 0.2088109 0.2400246 0.3088049
#[7,] 0.2298693 0.2411757 0.2851833 0.2359313
#[8,] 0.2196237 0.1941079 0.3017546 0.2755319
#[9,] 0.2514210 0.2352421 0.2441995 0.2580083
#
#, , 3
#
#[,1]      [,2]      [,3]      [,4]
#[1,] 0.3361180 0.2806397 0.3930593 0.3925858
#[2,] 0.3780252 0.3062918 0.3356780 0.4486956
#[3,] 0.4090534 0.2886780 0.3761269 0.3604488
#[4,] 0.3303153 0.3473472 0.3882696 0.4061389
#[5,] 0.3510128 0.3629823 0.3043323 0.3970610
#[6,] 0.3366603 0.3079409 0.3424458 0.4164231
#[7,] 0.3319855 0.3462912 0.3930013 0.3383752
#[8,] 0.3218208 0.2958522 0.4128891 0.3850848
#[9,] 0.3694168 0.3502802 0.3596994 0.3753940
#
#
#$PopPref
#[,1]      [,2]      [,3]
#[1,] 0.1846826 0.2442231 0.3068095
#[2,] 0.1652866 0.2169795 0.2788869
#[3,] 0.2002507 0.2573087 0.3208441
#[4,] 0.2178453 0.2778781 0.3437703
#
#$PopVar
#2.5%      50%    97.5% 
#27.45676 43.21658 49.71138 


## It might also be handy to extract the pop level preference results, so they can be graphed
## or whatever.  Here I extract those two vectors for the two choices and find means manually
## which match the means found above:
## first extract the population preference element from the main result list
popPrefMCMC.BS.S <- SimpPrefBS[[1]]$PopPref
dim(popPrefMCMC.BS.S)
# [1]     4 40000


## extract preferences for the three choices and find mean
prefBL.BS.S <- popPrefMCMC.BS.S[1,] 
mean(prefBL.BS.S)
# [1] 0.2451105

prefBS.BS.S <- popPrefMCMC.BS.S[2,]
mean(prefBS.BS.S)
# [1] 0.2182591

prefYL.BS.S <- popPrefMCMC.BS.S[3,]
mean(prefYL.BS.S)
# [1] 0.257376

prefYS.BS.S <- popPrefMCMC.BS.S[4,]
mean(prefYS.BS.S)
# [1] 0.2792544

## pairwise probabilities
## in this case, you interpret the result such that
## you have 100% confidence (or almost) that preference for large (purple) is greater than preference for small (green),
## and that you have 0% confidence that preference for small is greater than preference for large.
pairwiseProb(SimpPrefBS[[1]],burn=10000)

#          [,1]      [,2]      [,3]      [,4]
#[1,] 0.0000000 0.7143333 0.3982667 0.2624000 #BL
#[2,] 0.2856667 0.0000000 0.2164000 0.1185000 #BS
#[3,] 0.6017333 0.7836000 0.0000000 0.3497667 #YL
#[4,] 0.7376000 0.8815000 0.6502333 0.0000000 #YS


#### FOR SIMPLE PETAL CHOICE TRAINED BEES - TEST BS ####
FILENAME6 <- "STATS_Counts_Simp_TestPhaseAnalysis-BS_12-20-23_MC.csv" # BS trained bees -simp
head(simp.test.BS)

## For simple petal choice yellow long (YL) trained bees

## all bees in one group ('pop')
data4 <- subset(simp.test.BS,Pop==2,select=-c(BeeID))
data4 <- droplevels(data4)
## change pop label to 1, which is what bayesPref will expect for single analysis
data4$Pop <- rep(1,dim(data4)[1]) #called pop 1, but it's actually pop2. Just a quirk
data4 <- as.matrix(data4)
head(data4)

# run the model
SimpTestBS<-bayesPref(pData=data4,mcmcL=40000,pops=TRUE,dicburn=10000,dirvar=9)
#dic =  44.40121

# examine mixing, which is not great at 1000 reps
plot(SimpTestBS[[1]]$PopPref[1,],xlab="MCMC step",ylab="PopPref",cex=0.5)

## simple plot
prefPlot(SimpTestBS[[1]],ymax=35)

## means and credible intervals
## this next command will get you means and credible intervals returned in the $PopPref element
## the means are in the middle column, and the credible intervals are on either side
credibleIntervals(prefres = SimpTestBS[[1]],burn=10000,interval=0.95)  

#$IndPref
#, , 1
#
#[,1]      [,2]       [,3]       [,4]
#[1,] 0.15457965 0.1890124 0.09560877 0.10363794
#[2,] 0.10513987 0.1934386 0.08888190 0.07879275
#[3,] 0.09072731 0.2146783 0.09900384 0.08841780
#[4,] 0.14064726 0.1650015 0.08736386 0.07690585
#[5,] 0.17903821 0.1939756 0.12299960 0.11660533
#[6,] 0.10992040 0.1338568 0.09619026 0.10422954
#[7,] 0.12558316 0.1626032 0.13901452 0.18672444
#[8,] 0.15946827 0.2170645 0.09946660 0.05814326
#[9,] 0.09214727 0.2104879 0.15988955 0.07276294
#
#, , 2
#
#[,1]      [,2]      [,3]      [,4]
#[1,] 0.2701460 0.3136218 0.1981142 0.2045285
#[2,] 0.2307185 0.3417881 0.2113622 0.1963493
#[3,] 0.2105344 0.3554843 0.2147244 0.2008312
#[4,] 0.2747377 0.3077636 0.2062589 0.1911065
#[5,] 0.2776347 0.2963840 0.2120810 0.2039021
#[6,] 0.2423524 0.2796305 0.2226366 0.2328836
#[7,] 0.2150486 0.2593916 0.2286507 0.2864415
#[8,] 0.2791895 0.3467576 0.2054484 0.1539166
#[9,] 0.1975008 0.3387356 0.2775836 0.1703169
#
#, , 3
#
#[,1]      [,2]      [,3]      [,4]
#[1,] 0.4112393 0.4570570 0.3345067 0.3389529
#[2,] 0.3988172 0.5170555 0.3789975 0.3553723
#[3,] 0.3686553 0.5243166 0.3716156 0.3491306
#[4,] 0.4510736 0.4763825 0.3696066 0.3477115
#[5,] 0.3967262 0.4130956 0.3244932 0.3105539
#[6,] 0.4203097 0.4503831 0.4000179 0.4072240
#[7,] 0.3244995 0.3712012 0.3412619 0.4044975
#[8,] 0.4286609 0.4963378 0.3441395 0.2824838
#[9,] 0.3374790 0.4867229 0.4309844 0.2991568
#
#
#$PopPref
#[,1]      [,2]      [,3]
#[1,] 0.1749446 0.2438577 0.3884704
#[2,] 0.2391733 0.3176157 0.3962379
#[3,] 0.1498390 0.2237345 0.3203563
#[4,] 0.1293304 0.2059088 0.2908542

#$PopVar
#2.5%      50%    97.5% 
#16.89398 38.25675 49.54529 


## It might also be handy to extract the pop level preference results, so they can be graphed
## or whatever.  Here I extract those two vectors for the two choices and find means manually
## which match the means found above:
## first extract the population preference element from the main result list
popPrefTMCMC2.BS.S <- SimpTestBS[[1]]$PopPref
dim(popPrefTMCMC2.BS.S)
# [1]     4 40000

##FOR Experienced bees
prefBLt.BS.S <- popPrefTMCMC2.BS.S[1,] 
mean(prefBLt.BS.S)
# [1] 0.2502021

prefBSt.BS.S <- popPrefTMCMC2.BS.S[2,]
mean(prefBSt.BS.S)
# [1] 0.3174744

prefYLt.BS.S <- popPrefTMCMC2.BS.S[3,]
mean(prefYLt.BS.S)
# [1] 0.2251897

prefYSt.BS.S <- popPrefTMCMC2.BS.S[4,]
mean(prefYSt.BS.S)
# [1] 0.2071338

## pairwise probabilities
## in this case, you interpret the result such that
## you have 100% confidence (or almost) that preference for large (purple) is greater than preference for small (green),
## and that you have 0% confidence that preference for small is greater than preference for large.
pairwiseProb(SimpTestBS[[1]],burn=10000)

#          [,1]       [,2]      [,3]      [,4]
#[1,] 0.0000000 0.16016667 0.6243333 0.7335667 #BL
#[2,] 0.8398333 0.00000000 0.9048667 0.9474000 #BS
#[3,] 0.3756667 0.09513333 0.0000000 0.5815333 #YL
#[4,] 0.2664333 0.05260000 0.4184667 0.0000000 #YS


#### COMPARISONS BETWEEN THE TWO PHASES FOR EACH TARGET FLOWER TYPE - BS ####

pop1Full.BS.S <- SimpPrefBS[[1]]$PopPref[1,] # Pull out BL for the Naive bees
pop1Chopped.BS.S <- pop1Full.BS.S[10001:40000] # saves the steps after the burn in
#then for the other population
pop2Full.BS.S <- SimpTestBS[[1]]$PopPref[1,] # Pull out BL for the Experienced bees
pop2Chopped.BS.S <- pop2Full.BS.S[10001:40000] #Post burn in
dif <- pop1Chopped.BS.S - pop2Chopped.BS.S #How many times in the posterior 1 is bigger than the other
BLprob.BS.s <- length(dif[dif>0]) / 30000 # should be the amount post burn in throw out
BLprob.BS.s
# [1] 0.4912667

pop1Full.BS.S <- SimpPrefBS[[1]]$PopPref[2,] # Pull out BS for the Naive bees
pop1Chopped.BS.S <- pop1Full.BS.S[10001:40000] # saves the steps after the burn in
#then for the other population
pop2Full.BS.S <- SimpTestBS[[1]]$PopPref[2,] # Pull out BS for the Experienced bees
pop2Chopped.BS.S <- pop2Full.BS.S[10001:40000] #Post burn in
dif <- pop1Chopped.BS.S - pop2Chopped.BS.S #How many times in the posterior 1 is bigger than the other
BSprob.BS.s <- length(dif[dif>0]) / 30000 # should be the amount post burn in throw out
BSprob.BS.s
# [1] 0.0259

pop1Full.BS.S <- SimpPrefBS[[1]]$PopPref[3,] # Pull out YL for the Naive bees
pop1Chopped.BS.S <- pop1Full.BS.S[10001:40000] # saves the steps after the burn in
#then for the other population
pop2Full.BS.S <- SimpTestBS[[1]]$PopPref[3,] # Pull out YL for the Experienced bees
pop2Chopped.BS.S <- pop2Full.BS.S[10001:40000] #Post burn in
dif <- pop1Chopped.BS.S - pop2Chopped.BS.S #How many times in the posterior 1 is bigger than the other
YLprob.BS.s <- length(dif[dif>0]) / 30000 # should be the amount post burn in throw out
YLprob.BS.s
#[1] 0.7258

pop1Full.BS.S <- SimpPrefBS[[1]]$PopPref[4,] # Pull out YS for the Naive bees
pop1Chopped.BS.S <- pop1Full.BS.S[10001:40000] # saves the steps after the burn in
#then for the other population
pop2Full.BS.S <- SimpTestBS[[1]]$PopPref[4,] # Pull out YS for the Experienced bees
pop2Chopped.BS.S <- pop2Full.BS.S[10001:40000] #Post burn in
dif <- pop1Chopped.BS.S - pop2Chopped.BS.S #How many times in the posterior 1 is bigger than the other
YSprob.BS.s <- length(dif[dif>0]) / 30000 # should be the amount post burn in throw out
YSprob.BS.s
#[1] 0.9048


#### FOR DIFFICULT PETAL CHOICE TRAINED BEES - INITIAL PL ####
FILENAME3 <- "STATS_Counts_Diff_InitialPhaseAnalysis-PL_12-20-23_MC.csv" # PL trained bees
head(diff.pref.PL)

## all bees in one group ('pop')
data5 <- subset(diff.pref.PL,Pop==1,select=-c(BeeID))
data5 <- droplevels(data5)
data5 <- as.matrix(data5)
head(data5)

# run the model
DiffPrefPL <- bayesPref(pData=data5,mcmcL=40000,pops=TRUE,dicburn=10000,dirvar=9)
# dic =  61.37362

# examine mixing, which is not great at 1000 reps
plot(DiffPrefPL[[1]]$PopPref[1,],xlab="MCMC step",ylab="PopPref",cex=0.5)

## simple plot
prefPlot(DiffPrefPL[[1]],ymax=35)

## means and credible intervals
## this next command will get you means and credible intervals returned in the $PopPref element
## the means are in the middle column, and the credible intervals are on either side
credibleIntervals(prefres = DiffPrefPL[[1]],burn=10000,interval=0.95)  

#$IndPref
#, , 1
#
#[,1]       [,2]      [,3]      [,4]
#[1,] 0.1436288 0.14105289 0.1773340 0.1735982
#[2,] 0.1876495 0.15222675 0.1265656 0.1652017
#[3,] 0.1888726 0.12252661 0.1378226 0.1860856
#[4,] 0.1032856 0.08602595 0.1516208 0.2112112
#[5,] 0.1068227 0.12989553 0.1857885 0.1697777
#[6,] 0.1647493 0.13984489 0.1557991 0.1634338
#[7,] 0.1525857 0.11631513 0.1664144 0.1746641
#[8,] 0.1959285 0.11150075 0.1455173 0.1850434
#[9,] 0.1582438 0.15628466 0.1692840 0.1466639
#
#, , 2
#
#[,1]      [,2]      [,3]      [,4]
#[1,] 0.2308768 0.2268363 0.2684781 0.2660747
#[2,] 0.2826266 0.2415017 0.2104502 0.2568472
#[3,] 0.2817448 0.2052134 0.2228363 0.2813899
#[4,] 0.2071084 0.1814379 0.2640511 0.3349535
#[5,] 0.1990640 0.2248621 0.2913510 0.2741286
#[6,] 0.2582498 0.2284142 0.2465703 0.2577321
#[7,] 0.2482789 0.2022882 0.2641903 0.2748946
#[8,] 0.2917016 0.1903095 0.2316699 0.2777714
#[9,] 0.2490319 0.2452872 0.2621474 0.2352396
#
#
#, , 3
#
#[,1]      [,2]      [,3]      [,4]
#[1,] 0.3344235 0.3279578 0.3746102 0.3726730
#[2,] 0.3930010 0.3453561 0.3135981 0.3653615
#[3,] 0.3913893 0.3038064 0.3253281 0.3924570
#[4,] 0.3359091 0.3070553 0.4026078 0.4787465
#[5,] 0.3155497 0.3397222 0.4154730 0.3967024
#[6,] 0.3681318 0.3347682 0.3545159 0.3665135
#[7,] 0.3615779 0.3102223 0.3793277 0.3908760
#[8,] 0.3982382 0.2897433 0.3371576 0.3852550
#[9,] 0.3555419 0.3514196 0.3696513 0.3403754
#
#
#$PopPref
#[,1]      [,2]      [,3]
#[1,] 0.1899111 0.2518982 0.3157007
#[2,] 0.1613974 0.2188533 0.2796243
#[3,] 0.1937004 0.2539285 0.3192476
#[4,] 0.2076019 0.2746910 0.3380967
#
#$PopVar
#2.5%      50%    97.5% 
#27.96279 43.28543 49.74412 


## It might also be handy to extract the pop level preference results, so they can be graphed
## or whatever.  Here I extract those two vectors for the two choices and find means manually
## which match the means found above:
## first extract the population preference element from the main result list
popPrefMCMC.PL <- DiffPrefPL[[1]]$PopPref
dim(popPrefMCMC.PL)
# [1]     4 40000


## extract preferences for the three choices and find mean
prefBL.PL <- popPrefMCMC.PL[1,] 
mean(prefBL.PL)
# [1] 0.2515624

prefBS.PL <- popPrefMCMC.PL[2,]
mean(prefBS.PL)
# [1] 0.2194767

prefPL.PL <- popPrefMCMC.PL[3,]
mean(prefPL.PL)
# [1] 0.2546885

prefPS.PL <- popPrefMCMC.PL[4,]
mean(prefPS.PL)
# [1] 0.2742724


## pairwise probabilities
## in this case, you interpret the result such that
## you have 100% confidence (or almost) that preference for large (purple) is greater than preference for small (green),
## and that you have 0% confidence that preference for small is greater than preference for large.
pairwiseProb(DiffPrefPL[[1]],burn=10000)

#         [,1]      [,2]      [,3]      [,4]
#[1,] 0.0000000 0.7452000 0.4939667 0.3453000
#[2,] 0.2548000 0.0000000 0.2492000 0.1512333
#[3,] 0.5060333 0.7508000 0.0000000 0.3413667
#[4,] 0.6547000 0.8487667 0.6586333 0.0000000


#### FOR DIFFICULT PETAL CHOICE TRAINED BEES - TEST PL ####
FILENAME7 <- "STATS_Counts_Diff_TestPhaseAnalysis-PL_12-19-23_MC.csv" # PL trained bees
head(diff.test.PL)

## all bees in one group ('pop')
data6 <- subset(diff.test.PL,Pop==2,select=-c(BeeID))
data6 <- droplevels(data6)
## change pop label to 1, which is what bayesPref will expect for single analysis
data6$Pop <- rep(1,dim(data6)[1]) #called pop 1, but it's actually pop2. Just a quirk
data6 <- as.matrix(data6)
head(data6)

# run the model
DiffTestPL<-bayesPref(pData=data6,mcmcL=40000,pops=TRUE,dicburn=10000,dirvar=9)
#dic =  56.5414

# examine mixing, which is not great at 1000 reps
plot(DiffTestPL[[1]]$PopPref[1,],xlab="MCMC step",ylab="PopPref",cex=0.5)

## simple plot
prefPlot(DiffTestPL[[1]],ymax=35)

## means and credible intervals
## this next command will get you means and credible intervals returned in the $PopPref element
## the means are in the middle column, and the credible intervals are on either side
credibleIntervals(prefres = DiffTestPL[[1]],burn=10000,interval=0.95)  

#$IndPref
#, , 1
#
#[,1]       [,2]      [,3]      [,4]
#[1,] 0.11669763 0.09558066 0.1622178 0.1617261
#[2,] 0.12906482 0.10550186 0.2020832 0.2021770
#[3,] 0.13373472 0.13106067 0.1875277 0.1762066
#[4,] 0.09058898 0.14551362 0.1952685 0.1830665
#[5,] 0.16902269 0.09811838 0.1672899 0.1801072
#[6,] 0.09993335 0.13660562 0.1241993 0.2103626
#[7,] 0.11731692 0.12435019 0.1996440 0.1982264
#[8,] 0.10842093 0.10580759 0.1686103 0.1683950
#[9,] 0.16733939 0.12182562 0.1138798 0.2234141
#
#, , 2
#
#[,1]      [,2]      [,3]      [,4]
#[1,] 0.2234180 0.1971090 0.2827140 0.2823923
#[2,] 0.2125523 0.1832723 0.2981674 0.2974029
#[3,] 0.2207764 0.2151576 0.2838930 0.2714398
#[4,] 0.1726295 0.2360368 0.2974705 0.2833750
#[5,] 0.2667258 0.1776349 0.2657746 0.2791137
#[6,] 0.1932744 0.2370691 0.2281137 0.3289779
#[7,] 0.1977015 0.2052945 0.2945781 0.2941234
#[8,] 0.2115504 0.2046868 0.2848714 0.2854017
#[9,] 0.2612288 0.2054745 0.1982919 0.3257390
#
#, , 3
#
#[,1]      [,2]      [,3]      [,4]
#[1,] 0.3611188 0.3305907 0.4278699 0.4244827
#[2,] 0.3144410 0.2811586 0.4069618 0.4061714
#[3,] 0.3242738 0.3201530 0.3955872 0.3809931
#[4,] 0.2769191 0.3502452 0.4166261 0.4008129
#[5,] 0.3833116 0.2829006 0.3821081 0.3944578
#[6,] 0.3153049 0.3653450 0.3557273 0.4638210
#[7,] 0.2960611 0.3048579 0.4028667 0.4036407
#[8,] 0.3415227 0.3351283 0.4246069 0.4238005
#[9,] 0.3730162 0.3102724 0.3041172 0.4411916
#
#
#$PopPref
#[,1]      [,2]      [,3]
#[1,] 0.1592283 0.2199621 0.2889761
#[2,] 0.1548150 0.2091560 0.2786477
#[3,] 0.2053081 0.2694228 0.3447802
#[4,] 0.2288811 0.2963468 0.3674410
#
#$PopVar
#2.5%      50%    97.5% 
#23.53856 42.15044 49.65450 


## It might also be handy to extract the pop level preference results, so they can be graphed
## or whatever.  Here I extract those two vectors for the two choices and find means manually
## which match the means found above:
## first extract the population preference element from the main result list
popPrefTMCMC2.PL <- DiffTestPL[[1]]$PopPref
dim(popPrefTMCMC2.PL)
# [1]     4 40000

##FOR Experienced bees
prefBLt.PL <- popPrefTMCMC2.PL[1,] 
mean(prefBLt.PL)
# [1] 0.2205243

prefBSt.PL <- popPrefTMCMC2.PL[2,]
mean(prefBSt.PL)
# [1] 0.2109441

prefPLt.PL <- popPrefTMCMC2.PL[3,]
mean(prefPLt.PL)
# [1] 0.2721271

prefPSt.PL <- popPrefTMCMC2.PL[4,]
mean(prefPSt.PL)
# [1] 0.2964045

## pairwise probabilities
## in this case, you interpret the result such that
## you have 100% confidence (or almost) that preference for large (purple) is greater than preference for small (green),
## and that you have 0% confidence that preference for small is greater than preference for large.
pairwiseProb(DiffTestPL[[1]],burn=10000)

#          [,1]      [,2]      [,3]       [,4]
#[1,] 0.0000000 0.5866667 0.1764000 0.08870000
#[2,] 0.4133333 0.0000000 0.1389000 0.05543333
#[3,] 0.8236000 0.8611000 0.0000000 0.34466667
#[4,] 0.9113000 0.9445667 0.6553333 0.00000000


#### COMPARISONS BETWEEN THE TWO PHASES FOR EACH TARGET FLOWER TYPE - PL ####

pop1Full.PL <- DiffPrefPL[[1]]$PopPref[1,] # Pull out BL for the Naive bees
pop1Chopped.PL <- pop1Full.PL[10001:40000] # saves the steps after the burn in
#then for the other population
pop2Full.PL <- DiffTestPL[[1]]$PopPref[1,] # Pull out BL for the Experienced bees
pop2Chopped.PL <- pop2Full.PL[10001:40000] #Post burn in
dif <- pop1Chopped.PL - pop2Chopped.PL #How many times in the posterior 1 is bigger than the other
BLprob.PL <- length(dif[dif>0]) / 30000 # should be the amount post burn in throw out
BLprob.PL
#[1] 0.757

pop1Full.PL <- DiffPrefPL[[1]]$PopPref[2,] # Pull out BS for the Naive bees
pop1Chopped.PL <- pop1Full.PL[10001:40000] # saves the steps after the burn in
#then for the other population
pop2Full.PL <- DiffTestPL[[1]]$PopPref[2,] # Pull out BS for the Experienced bees
pop2Chopped.PL <- pop2Full.PL[10001:40000] #Post burn in
dif <- pop1Chopped.PL - pop2Chopped.PL #How many times in the posterior 1 is bigger than the other
BSprob.PL <- length(dif[dif>0]) / 30000 # should be the amount post burn in throw out
BSprob.PL
#[1] 0.5819667

pop1Full.PL <- DiffPrefPL[[1]]$PopPref[3,] # Pull out PL for the Naive bees
pop1Chopped.PL <- pop1Full.PL[10001:40000] # saves the steps after the burn in
#then for the other population
pop2Full.PL <- DiffTestPL[[1]]$PopPref[3,] # Pull out PL for the Experienced bees
pop2Chopped.PL <- pop2Full.PL[10001:40000] #Post burn in
dif <- pop1Chopped.PL - pop2Chopped.PL #How many times in the posterior 1 is bigger than the other
PLprob.PL <- length(dif[dif>0]) / 30000 # should be the amount post burn in throw out
PLprob.PL
#[1] 0.3560667

pop1Full.PL <- DiffPrefPL[[1]]$PopPref[4,] # Pull out PS for the Naive bees
pop1Chopped.PL <- pop1Full.PL[10001:40000] # saves the steps after the burn in
#then for the other population
pop2Full.PL <- DiffTestPL[[1]]$PopPref[4,] # Pull out PS for the Experienced bees
pop2Chopped.PL <- pop2Full.PL[10001:40000] #Post burn in
dif <- pop1Chopped.PL - pop2Chopped.PL #How many times in the posterior 1 is bigger than the other
PSprob.PL <- length(dif[dif>0]) / 30000 # should be the amount post burn in throw out
PSprob.PL
#[1] 0.3237333


#### Rerunning stuff that didn't mix with dirvar=5 ####

#### FOR SIMPLE PETAL CHOICE TRAINED BEES - INITIAL YL ####
FILENAME1 <- "STATS_Counts_Simp_InitialPhaseAnalysis-YL_12-19-23_MC.csv" # YL trained bees
head(simp.pref.YL)

## all bees in one group ('pop')
data1 <- subset(simp.pref.YL,Pop==1,select=-c(BeeID))
data1 <- droplevels(data1)
data1 <- as.matrix(data1)
head(data1)

# run the model
#SimpPrefYL <- bayesPref(pData=data1,mcmcL=400,pops=TRUE,dicburn=100,dirvar=9) #shortened for testing the code faster, not for use in analysis
SimpPrefYL <- bayesPref(pData=data1,mcmcL=40000,pops=TRUE,dicburn=10000,dirvar=5)

# Results, 40000 steps
# dic =  62.65259   
# reran to check 12/26/23 - dic = 61.96356 

# examine mixing, which is not great at 1000 reps
plot(SimpPrefYL[[1]]$PopPref[1,],xlab="MCMC step",ylab="PopPref",cex=0.5)
#mixing looks good

## simple plot
prefPlot(SimpPrefYL[[1]],ymax=35)

## means and credible intervals
## this next command will get you means and credible intervals returned in the $PopPref element
## the means are in the middle column, and the credible intervals are on either side
#credibleIntervals(prefres = SimpPrefYL[[1]],burn=100,interval=0.95) #shortened for testing the code faster, not for use in analysis
credibleIntervals(prefres = SimpPrefYL[[1]],burn=10000,interval=0.95)  

#$IndPref
#, , 1
#
#[,1]      [,2]      [,3]       [,4]
#[1,] 0.1342693 0.1597313 0.1850912 0.13094885
#[2,] 0.1821970 0.1094103 0.1504162 0.15272996
#[3,] 0.1120209 0.1634124 0.1864606 0.18010508
#[4,] 0.1561108 0.1572492 0.1597223 0.14061705
#[5,] 0.1097429 0.1256644 0.1563682 0.17632271
#[6,] 0.1748872 0.1996283 0.1358352 0.11829787
#[7,] 0.1342075 0.1250651 0.1826037 0.18538328
#[8,] 0.2108466 0.2126443 0.1167353 0.08739416
#
#, , 2
#
#[,1]      [,2]      [,3]      [,4]
#[1,] 0.2255455 0.2562009 0.2861971 0.2223302
#[2,] 0.2869433 0.2008507 0.2486175 0.2531921
#[3,] 0.1918439 0.2520133 0.2773886 0.2703638
#[4,] 0.2494658 0.2530585 0.2545547 0.2329395
#[5,] 0.2075896 0.2279975 0.2650576 0.2874452
#[6,] 0.2687549 0.2962032 0.2232832 0.2024227
#[7,] 0.2212404 0.2116607 0.2769238 0.2811972
#[8,] 0.3100095 0.3133166 0.2005754 0.1661949
#
#, , 3
#
#[,1]      [,2]      [,3]      [,4]
#[1,] 0.3366441 0.3733434 0.4046058 0.3327317
#[2,] 0.4104118 0.3142967 0.3679028 0.3739004
#[3,] 0.2888844 0.3562305 0.3850759 0.3775162
#[4,] 0.3635989 0.3669611 0.3696174 0.3441512
#[5,] 0.3296854 0.3539597 0.3962116 0.4216395
#[6,] 0.3775708 0.4095077 0.3302299 0.3055355
#[7,] 0.3265500 0.3179047 0.3893921 0.3933757
#[8,] 0.4259796 0.4297648 0.3046270 0.2683089
#
#
#$PopPref
#[,1]      [,2]      [,3]
#[1,] 0.1818944 0.2463896 0.3146009
#[2,] 0.1873703 0.2527900 0.3246620
#[3,] 0.1952355 0.2566454 0.3317895
#[4,] 0.1733843 0.2401051 0.3134870
#
#$PopVar
#2.5%      50%    97.5% 
#21.05232 41.18602 49.63456

## It might also be handy to extract the pop level preference results, so they can be graphed
## or whatever.  Here I extract those two vectors for the two choices and find means manually
## which match the means found above:
## first extract the population preference element from the main result list
popPrefMCMC.YL <- SimpPrefYL[[1]]$PopPref
dim(popPrefMCMC.YL)
# [1]     4 40000


## extract preferences for the three choices and find mean
prefBL.YL <- popPrefMCMC.YL[1,] 
mean(prefBL.YL)
# [1] 0.2469298
# v2 [1] 0.2468545

prefBS.YL <- popPrefMCMC.YL[2,]
mean(prefBS.YL)
# [1] 0.2536451
#v2 [1] 0.2549079

prefYL.YL <- popPrefMCMC.YL[3,]
mean(prefYL.YL)
# [1] 0.2576055
# v2 [1] 0.2562121

prefYS.YL <- popPrefMCMC.YL[4,]
mean(prefYS.YL)
# [1] 0.2418196
# v2 [1] 0.2420255

## pairwise probabilities
## in this case, you interpret the result such that
## you have 100% confidence (or almost) that preference for large (purple) is greater than preference for small (green),
## and that you have 0% confidence that preference for small is greater than preference for large.
#pairwiseProb(SimpPrefYL[[1]],burn=100) #shortened for testing the code faster, not for use in analysis
pairwiseProb(SimpPrefYL[[1]],burn=10000)

#          [,1]      [,2]      [,3]      [,4]
#[1,] 0.0000000 0.4513333 0.4306333 0.5497667
#[2,] 0.5486667 0.0000000 0.4664333 0.5852667
#[3,] 0.5693667 0.5335667 0.0000000 0.6223333
#[4,] 0.4502333 0.4147333 0.3776667 0.0000000

#v2 12-16-23
#          [,1]      [,2]      [,3]      [,4]
#[1,] 0.0000000 0.4467667 0.4238333 0.5327333
#[2,] 0.5532333 0.0000000 0.4872333 0.5932333
#[3,] 0.5761667 0.5127667 0.0000000 0.6072333
#[4,] 0.4672667 0.4067667 0.3927667 0.0000000


#### FOR SIMPLE PETAL CHOICE TRAINED BEES - TEST YL ####
FILENAME5 <- "STATS_Counts_Simp_TestPhaseAnalysis-YL_12-20-23_MC.csv" # YL trained bees
head(simp.test.YL)

## For simple petal choice yellow long (YL) trained bees

## all bees in one group ('pop')
data2 <- subset(simp.test.YL,Pop==2,select=-c(BeeID))
data2 <- droplevels(data2)
## change pop label to 1, which is what bayesPref will expect for single analysis
data2$Pop <- rep(1,dim(data2)[1]) #called pop 1, but it's actually pop2. Just a quirk
data2 <- as.matrix(data2)
head(data2)

# run the model
#SimpTestYL<-bayesPref(pData=data2,mcmcL=400,pops=TRUE,dicburn=100,dirvar=9) #shortened for testing the code faster, not for use in analysis
SimpTestYL<-bayesPref(pData=data2,mcmcL=40000,pops=TRUE,dicburn=10000,dirvar=5)
#dic =  47.38347
#v2 12/26/23 dic =  47.51576 

# examine mixing, which is not great at 1000 reps
plot(SimpTestYL[[1]]$PopPref[1,],xlab="MCMC step",ylab="PopPref",cex=0.5)
#mixing good

## simple plot
prefPlot(SimpTestYL[[1]],ymax=35)

## means and credible intervals
## this next command will get you means and credible intervals returned in the $PopPref element
## the means are in the middle column, and the credible intervals are on either side
#credibleIntervals(prefres = SimpTestYL[[1]],burn=100,interval=0.95) #shortened for testing the code faster, not for use in analysis
credibleIntervals(prefres = SimpTestYL[[1]],burn=10000,interval=0.95)  

#$IndPref
#, , 1
#
#[,1]       [,2]      [,3]      [,4]
#[1,] 0.08984883 0.05655206 0.1918629 0.2086919
#[2,] 0.06761988 0.05536386 0.1993806 0.3240778
#[3,] 0.06116812 0.07088118 0.2086744 0.2085657
#[4,] 0.06617294 0.06156336 0.2410410 0.1675741
#[5,] 0.04237889 0.05767138 0.2271790 0.1626496
#[6,] 0.05749606 0.09944887 0.2377429 0.2113601
#[7,] 0.15460301 0.10750988 0.1843029 0.1713801
#[8,] 0.09946646 0.05728518 0.2160319 0.2626009
#
#, , 2
#
#[,1]      [,2]      [,3]      [,4]
#[1,] 0.1866311 0.1409483 0.3193385 0.3389474
#[2,] 0.1357960 0.1191203 0.2989903 0.4370023
#[3,] 0.1486920 0.1599706 0.3378963 0.3394729
#[4,] 0.1576063 0.1499759 0.3781788 0.2985220
#[5,] 0.1392797 0.1546545 0.3779397 0.3089354
#[6,] 0.1298391 0.1842620 0.3525545 0.3231201
#[7,] 0.2454525 0.1887917 0.2842834 0.2718842
#[8,] 0.1781184 0.1222818 0.3189295 0.3713322
#
#, , 3
#
#[,1]      [,2]      [,3]      [,4]
#[1,] 0.3186446 0.2613294 0.4646715 0.4848934
#[2,] 0.2291087 0.2080451 0.4113294 0.5574107
#[3,] 0.2726830 0.2871990 0.4836980 0.4861102
#[4,] 0.2857085 0.2796080 0.5334350 0.4502547
#[5,] 0.2810056 0.3020633 0.5535709 0.4763088
#[6,] 0.2312930 0.2956646 0.4775653 0.4487626
#[7,] 0.3583091 0.2936086 0.3982529 0.3859506
#[8,] 0.2787887 0.2135646 0.4344365 0.4893789
#
#
#$PopPref
#[,1]      [,2]      [,3]
#[1,] 0.10484255 0.1693734 0.2391087
#[2,] 0.09790725 0.1589747 0.2282741
#[3,] 0.25493362 0.3348631 0.4123561
#[4,] 0.25124890 0.3348111 0.4216239
#
#$PopVar
#2.5%      50%    97.5% 
#17.81985 38.66696 49.44479 


## It might also be handy to extract the pop level preference results, so they can be graphed
## or whatever.  Here I extract those two vectors for the two choices and find means manually
## which match the means found above:
## first extract the population preference element from the main result list
popPrefTMCMC2.YL <- SimpTestYL[[1]]$PopPref
dim(popPrefTMCMC2.YL)
# [1]     4 40000

##FOR Experienced bees
prefBLt.YL <- popPrefTMCMC2.YL[1,] 
mean(prefBLt.YL)
# [1] 0.1717428
# v2 [1] 0.1710027

prefBSt.YL <- popPrefTMCMC2.YL[2,]
mean(prefBSt.YL)
# [1] 0.1602178
#v2 [1] 0.159027

prefYLt.YL <- popPrefTMCMC2.YL[3,]
mean(prefYLt.YL)
# [1] 0.3342451
#v2 [1] 0.3357445

prefYSt.YL <- popPrefTMCMC2.YL[4,]
mean(prefYSt.YL)
# [1] 0.3337944
#v2 [1] 0.3342259

## pairwise probabilities
## in this case, you interpret the result such that
## you have 100% confidence (or almost) that preference for large (purple) is greater than preference for small (green),
## and that you have 0% confidence that preference for small is greater than preference for large.
#pairwiseProb(SimpTestYL[[1]],burn=100) #shortened for testing the code faster, not for use in analysis
pairwiseProb(SimpTestYL[[1]],burn=10000)
#          [,1]      [,2]        [,3]        [,4]
#[1,] 0.0000000 0.5893667 0.003800000 0.008300000
#[2,] 0.4106333 0.0000000 0.001066667 0.002466667
#[3,] 0.9962000 0.9989333 0.000000000 0.495233333
#[4,] 0.9917000 0.9975333 0.504766667 0.000000000\

#v2 12-26-23
#          [,1]      [,2]        [,3]        [,4]
#[1,] 0.0000000 0.5862333 0.003966667 0.006200000
#[2,] 0.4137667 0.0000000 0.001833333 0.004666667
#[3,] 0.9960333 0.9981667 0.000000000 0.505366667
#[4,] 0.9938000 0.9953333 0.494633333 0.000000000


#### COMPARISONS BETWEEN THE TWO PHASES FOR EACH TARGET FLOWER TYPE - YL ####

pop1Full.YL <- SimpPrefYL[[1]]$PopPref[1,] # Pull out BL for the Naive bees
pop1Chopped.YL <- pop1Full.YL[10001:40000] # saves the steps after the burn in
#then for the other population
pop2Full.YL <- SimpTestYL[[1]]$PopPref[1,] # Pull out BL for the Experienced bees
pop2Chopped.YL <- pop2Full.YL[10001:40000] #Post burn in
dif <- pop1Chopped.YL - pop2Chopped.YL #How many times in the posterior 1 is bigger than the other
BLprob.YL <- length(dif[dif>0]) / 30000 # should be the amount post burn in throw out
BLprob.YL
# [1] 0.9498667
#v2 [1] 0.9398333

pop1Full.YL <- SimpPrefYL[[1]]$PopPref[2,] # Pull out BS for the Naive bees
pop1Chopped.YL <- pop1Full.YL[10001:40000] # saves the steps after the burn in
#then for the other population
pop2Full.YL <- SimpTestYL[[1]]$PopPref[2,] # Pull out BS for the Experienced bees
pop2Chopped.YL <- pop2Full.YL[10001:40000] #Post burn in
dif <- pop1Chopped.YL - pop2Chopped.YL #How many times in the posterior 1 is bigger than the other
BSprob.YL <- length(dif[dif>0]) / 30000 # should be the amount post burn in throw out
BSprob.YL
# [1] 0.9729
#v2 [1] 0.9761667

pop1Full.YL <- SimpPrefYL[[1]]$PopPref[3,] # Pull out YL for the Naive bees
pop1Chopped.YL <- pop1Full.YL[10001:40000] # saves the steps after the burn in
#then for the other population
pop2Full.YL <- SimpTestYL[[1]]$PopPref[3,] # Pull out YL for the Experienced bees
pop2Chopped.YL <- pop2Full.YL[10001:40000] #Post burn in
dif <- pop1Chopped.YL - pop2Chopped.YL #How many times in the posterior 1 is bigger than the other
YLprob.YL <- length(dif[dif>0]) / 30000 # should be the amount post burn in throw out
YLprob.YL
# [1] 0.07756667
# v2 [1] 0.07326667

pop1Full.YL <- SimpPrefYL[[1]]$PopPref[4,] # Pull out YS for the Naive bees
pop1Chopped.YL <- pop1Full.YL[10001:40000] # saves the steps after the burn in
#then for the other population
pop2Full.YL <- SimpTestYL[[1]]$PopPref[4,] # Pull out YS for the Experienced bees
pop2Chopped.YL <- pop2Full.YL[10001:40000] #Post burn in
dif <- pop1Chopped.YL - pop2Chopped.YL #How many times in the posterior 1 is bigger than the other
YSprob.YL <- length(dif[dif>0]) / 30000 # should be the amount post burn in throw out
YSprob.YL
# [1] 0.04413333
#v2 [1] 0.04526667


#### FOR DIFFICULT PETAL CHOICE TRAINED BEES - INITIAL BS ####
FILENAME4 <- "STATS_Counts_Diff_InitialPhaseAnalysis-BS_12-20-23_MC.csv" # BS trained bees
head(diff.pref.BS)

## all bees in one group ('pop')
data7 <- subset(diff.pref.BS,Pop==1,select=-c(BeeID))
data7 <- droplevels(data7)
data7 <- as.matrix(data7)
head(data7)

# run the model
DiffPrefBS <- bayesPref(pData=data7,mcmcL=40000,pops=TRUE,dicburn=10000,dirvar=5)
# dic =  70.39279  

# examine mixing, which is not great at 1000 reps
plot(DiffPrefBS[[1]]$PopPref[1,],xlab="MCMC step",ylab="PopPref",cex=0.5)

## simple plot
prefPlot(DiffPrefBS[[1]],ymax=35)

## means and credible intervals
## this next command will get you means and credible intervals returned in the $PopPref element
## the means are in the middle column, and the credible intervals are on either side
credibleIntervals(prefres = DiffPrefBS[[1]],burn=10000,interval=0.95) 

#$IndPref
#, , 1
#
#[,1]      [,2]      [,3]      [,4]
#[1,] 0.12902655 0.1975271 0.1357178 0.1559646
#[2,] 0.14476872 0.1663881 0.1501331 0.1595721
#[3,] 0.08510626 0.1713867 0.1630842 0.1473235
#[4,] 0.14024589 0.1401078 0.1873506 0.1751589
#[5,] 0.16773100 0.1452922 0.1622031 0.1606589
#[6,] 0.12594039 0.1800449 0.1758324 0.1399261
#[7,] 0.09090748 0.1635863 0.1450035 0.2047820
#[8,] 0.16713732 0.1877993 0.1524439 0.1303222
#[9,] 0.13677239 0.1472769 0.1762540 0.1728783
#[10,] 0.13239185 0.1868495 0.1603631 0.1465348
#[11,] 0.11547045 0.1968823 0.1744074 0.1486126
#
#, , 2
#
#[,1]      [,2]      [,3]      [,4]
#[1,] 0.2159826 0.2985873 0.2257021 0.2496959
#[2,] 0.2340221 0.2612319 0.2426752 0.2526513
#[3,] 0.1750382 0.2829698 0.2756121 0.2538115
#[4,] 0.2235238 0.2243722 0.2790459 0.2642343
#[5,] 0.2568474 0.2329764 0.2523570 0.2494565
#[6,] 0.2106531 0.2775288 0.2725141 0.2291343
#[7,] 0.1737923 0.2621417 0.2419903 0.3115488
#[8,] 0.2576511 0.2817807 0.2392542 0.2126550
#[9,] 0.2222998 0.2358197 0.2677476 0.2656346
#[10,] 0.2189360 0.2834009 0.2517814 0.2365270
#[11,] 0.1957213 0.2938048 0.2644503 0.2376116
#
#, , 3
#
#[,1]      [,2]      [,3]      [,4]
#[1,] 0.3249187 0.4151732 0.3347567 0.3633755
#[2,] 0.3421955 0.3715554 0.3513461 0.3643108
#[3,] 0.2962067 0.4163728 0.4072194 0.3838539
#[4,] 0.3244633 0.3247602 0.3846123 0.3688966
#[5,] 0.3620937 0.3366448 0.3567591 0.3557868
#[6,] 0.3179363 0.3902790 0.3837889 0.3380736
#[7,] 0.2825466 0.3804031 0.3584061 0.4339464
#[8,] 0.3636419 0.3894993 0.3435874 0.3162153
#[9,] 0.3271389 0.3394817 0.3756578 0.3748772
#[10,] 0.3256034 0.3956897 0.3619726 0.3448350
#[11,] 0.2969003 0.4032371 0.3710018 0.3432167
#
#
#$PopPref
#[,1]      [,2]      [,3]
#[1,] 0.1653558 0.2188858 0.2768763
#[2,] 0.2131232 0.2682979 0.3287119
#[3,] 0.2039633 0.2583113 0.3153740
#[4,] 0.1979577 0.2528889 0.3129451
#
#$PopVar
#2.5%      50%    97.5% 
#28.43453 43.91616 49.76007 


## It might also be handy to extract the pop level preference results, so they can be graphed
## or whatever.  Here I extract those two vectors for the two choices and find means manually
## which match the means found above:
## first extract the population preference element from the main result list
popPrefMCMC.BS.D <- DiffPrefBS[[1]]$PopPref
dim(popPrefMCMC.BS.D)
# [1]     4 40000


## extract preferences for the three choices and find mean
prefBL.BS.D <- popPrefMCMC.BS.D[1,] 
mean(prefBL.BS.D)
# [1] 0.2195958

prefBS.BS.D <- popPrefMCMC.BS.D[2,]
mean(prefBS.BS.D)
# [1] 0.2681205

prefPL.BS.D <- popPrefMCMC.BS.D[3,]
mean(prefPL.BS.D)
# [1] 0.2587532

prefPS.BS.D <- popPrefMCMC.BS.D[4,]
mean(prefPS.BS.D)
# [1] 0.2535304

## pairwise probabilities
## in this case, you interpret the result such that
## you have 100% confidence (or almost) that preference for large (purple) is greater than preference for small (green),
## and that you have 0% confidence that preference for small is greater than preference for large.
pairwiseProb(DiffPrefBS[[1]],burn=10000)
#          [,1]      [,2]      [,3]      [,4]
#[1,] 0.0000000 0.1463000 0.2023667 0.2308000
#[2,] 0.8537000 0.0000000 0.5879000 0.6109333
#[3,] 0.7976333 0.4121000 0.0000000 0.5394000
#[4,] 0.7692000 0.3890667 0.4606000 0.0000000


#### FOR DIFFICULT PETAL CHOICE TRAINED BEES - TEST BS (not rerun) ####
FILENAME8 <- "STATS_Counts_Diff_TestPhaseAnalysis-PL_12-19-23_MC.csv" # BS trained bees
head(diff.test.BS)

## all bees in one group ('pop')
data8 <- subset(diff.test.BS,Pop==2,select=-c(BeeID))
data8 <- droplevels(data8)
## change pop label to 1, which is what bayesPref will expect for single analysis
data8$Pop <- rep(1,dim(data8)[1]) #called pop 1, but it's actually pop2. Just a quirk
data8 <- as.matrix(data8)
head(data8)

# run the model
DiffTestBS<-bayesPref(pData=data8,mcmcL=40000,pops=TRUE,dicburn=10000,dirvar=9)
#dic =  55.28344

# examine mixing, which is not great at 1000 reps
plot(DiffTestBS[[1]]$PopPref[1,],xlab="MCMC step",ylab="PopPref",cex=0.5)

## simple plot
prefPlot(DiffTestBS[[1]],ymax=35)

## means and credible intervals
## this next command will get you means and credible intervals returned in the $PopPref element
## the means are in the middle column, and the credible intervals are on either side
credibleIntervals(prefres = DiffTestBS[[1]],burn=10000,interval=0.95)  

#$IndPref
#, , 1
#
#[,1]       [,2]      [,3]      [,4]
#[1,] 0.11609214 0.09720214 0.1626570 0.1620770
#[2,] 0.12941794 0.10666376 0.2020966 0.2015211
#[3,] 0.13450771 0.13167471 0.1880912 0.1768029
#[4,] 0.09257473 0.14553127 0.1944699 0.1832076
#[5,] 0.16905480 0.09886911 0.1689871 0.1806106
#[6,] 0.09990983 0.13724387 0.1270971 0.2114412
#[7,] 0.11893046 0.12524010 0.1986128 0.1982663
#[8,] 0.11099580 0.10640209 0.1693641 0.1697712
#[9,] 0.16773413 0.12343709 0.1135311 0.2255946
#
#, , 2
#
#[,1]      [,2]      [,3]      [,4]
#[1,] 0.2237423 0.1974974 0.2818896 0.2826071
#[2,] 0.2125465 0.1843441 0.2966283 0.2976216
#[3,] 0.2201146 0.2168055 0.2834580 0.2706542
#[4,] 0.1729254 0.2361647 0.2972659 0.2838014
#[5,] 0.2648991 0.1791171 0.2662764 0.2794202
#[6,] 0.1944902 0.2383003 0.2281252 0.3276338
#[7,] 0.1977981 0.2059692 0.2940581 0.2939605
#[8,] 0.2108139 0.2056372 0.2851557 0.2847144
#[9,] 0.2604479 0.2063499 0.1985791 0.3258394
#
#, , 3
#
#[,1]      [,2]      [,3]      [,4]
#[1,] 0.3598272 0.3314755 0.4253340 0.4235905
#[2,] 0.3132272 0.2828149 0.4060645 0.4050435
#[3,] 0.3231235 0.3200055 0.3943084 0.3804358
#[4,] 0.2771518 0.3480928 0.4142465 0.4000576
#[5,] 0.3792371 0.2835399 0.3792327 0.3936073
#[6,] 0.3128689 0.3662569 0.3539525 0.4614965
#[7,] 0.2952488 0.3054727 0.4023239 0.4023060
#[8,] 0.3390452 0.3346372 0.4229279 0.4221485
#[9,] 0.3693924 0.3109251 0.3015840 0.4388818
#
#
#$PopPref
#[,1]      [,2]      [,3]
#[1,] 0.1627072 0.2207894 0.2853410
#[2,] 0.1557442 0.2118598 0.2769183
#[3,] 0.2060152 0.2704243 0.3390170
#[4,] 0.2338089 0.2959371 0.3631001
#
#$PopVar
#2.5%      50%    97.5% 
#25.34823 43.33841 49.74835 


## It might also be handy to extract the pop level preference results, so they can be graphed
## or whatever.  Here I extract those two vectors for the two choices and find means manually
## which match the means found above:
## first extract the population preference element from the main result list
popPrefTMCMC2.BS.D <- DiffTestBS[[1]]$PopPref
dim(popPrefTMCMC2.BS.D)
# [1]     4 40000

##FOR Experienced bees
prefBLt.BS.D <- popPrefTMCMC2.BS.D[1,] 
mean(prefBLt.BS.D)
# [1] 0.2209357

prefBSt.BS.D <- popPrefTMCMC2.BS.D[2,]
mean(prefBSt.BS.D)
# [1] 0.2128365

prefPLt.BS.D <- popPrefTMCMC2.BS.D[3,]
mean(prefPLt.BS.D)
# [1] 0.2706729

prefPSt.BS.D <- popPrefTMCMC2.BS.D[4,]
mean(prefPSt.BS.D)
# [1] 0.2955549


## pairwise probabilities
## in this case, you interpret the result such that
## you have 100% confidence (or almost) that preference for large (purple) is greater than preference for small (green),
## and that you have 0% confidence that preference for small is greater than preference for large.
pairwiseProb(DiffTestBS[[1]],burn=10000)

#[,1]      [,2]      [,3]       [,4]
#[1,] 0.0000000 0.5622000 0.1723333 0.08276667
#[2,] 0.4378000 0.0000000 0.1321667 0.05770000
#[3,] 0.8276667 0.8678333 0.0000000 0.33463333
#[4,] 0.9172333 0.9423000 0.6653667 0.00000000


#### COMPARISONS BETWEEN THE TWO PHASES FOR EACH TARGET FLOWER TYPE - BS ####

pop1Full.BS.D <- DiffPrefBS[[1]]$PopPref[1,] # Pull out BL for the Naive bees
pop1Chopped.BS.D <- pop1Full.BS.D[10001:40000] # saves the steps after the burn in
#then for the other population
pop2Full.BS.D <- DiffTestBS[[1]]$PopPref[1,] # Pull out BL for the Experienced bees
pop2Chopped.BS.D <- pop2Full.BS.D[10001:40000] #Post burn in
dif <- pop1Chopped.BS.D - pop2Chopped.BS.D #How many times in the posterior 1 is bigger than the other
BLprob.BS.d <- length(dif[dif>0]) / 30000 # should be the amount post burn in throw out
BLprob.BS.d
# [1] 0.4898667

pop1Full.BS.D <- DiffPrefBS[[1]]$PopPref[2,] # Pull out BS for the Naive bees
pop1Chopped.BS.D <- pop1Full.BS.D[10001:40000] # saves the steps after the burn in
#then for the other population
pop2Full.BS.D <- DiffTestBS[[1]]$PopPref[2,] # Pull out BS for the Experienced bees
pop2Chopped.BS.D <- pop2Full.BS.D[10001:40000] #Post burn in
dif <- pop1Chopped.BS.D - pop2Chopped.BS.D #How many times in the posterior 1 is bigger than the other
BSprob.BS.d <- length(dif[dif>0]) / 30000 # should be the amount post burn in throw out
BSprob.BS.d
# [1] 0.9124

pop1Full.BS.D <- DiffPrefBS[[1]]$PopPref[3,] # Pull out PL for the Naive bees
pop1Chopped.BS.D <- pop1Full.BS.D[10001:40000] # saves the steps after the burn in
#then for the other population
pop2Full.BS.D <- DiffTestBS[[1]]$PopPref[3,] # Pull out PL for the Experienced bees
pop2Chopped.BS.D <- pop2Full.BS.D[10001:40000] #Post burn in
dif <- pop1Chopped.BS.D - pop2Chopped.BS.D #How many times in the posterior 1 is bigger than the other
PLprob.BS.d <- length(dif[dif>0]) / 30000 # should be the amount post burn in throw out
PLprob.BS.d
# [1] 0.3891667

pop1Full.BS.D <- DiffPrefBS[[1]]$PopPref[4,] # Pull out PS for the Naive bees
pop1Chopped.BS.D <- pop1Full.BS.D[10001:40000] # saves the steps after the burn in
#then for the other population
pop2Full.BS.D <- DiffTestBS[[1]]$PopPref[4,] # Pull out PS for the Experienced bees
pop2Chopped.BS.D <- pop2Full.BS.D[10001:40000] #Post burn in
dif <- pop1Chopped.BS.D - pop2Chopped.BS.D #How many times in the posterior 1 is bigger than the other
PSprob.BS.d <- length(dif[dif>0]) / 30000 # should be the amount post burn in throw out
PSprob.BS.d
# [1] 0.1773667


#### *RE-RUNNING TO TEST FOR DIFFERENCES BY TREATMENT RATHER THAN SUB-TREATMENT* ####

#### FOR SIMPLE TRAINED BEES - INITIAL ####
FILENAME11 <- "STATS_Counts_Simp_InitialPhaseAnalysis-BS_12-19-23_MC.csv" 
simppref <- read.csv("~/LL stats/STATS_Counts_Simp_InitialPhaseAnalysis_2-14-24_MC.csv", sep=",", header=TRUE,check.names=FALSE)
head(simppref)

## all bees in one group ('pop')
data1 <- subset(simppref,Pop==1,select=-c(BeeID))
data1 <- droplevels(data1)
data1 <- as.matrix(data1)
head(data1)

# run the model
SimpPref <- bayesPref(pData=data1,mcmcL=40000,pops=TRUE,dicburn=10000,dirvar=9)
#dic =  115.6964 

# examine mixing, which is not great at 1000 reps
plot(SimpPref[[1]]$PopPref[1,],xlab="MCMC step",ylab="PopPref",cex=0.5)

## simple plot
prefPlot(SimpPref[[1]],ymax=35)

## means and credible intervals
## this next command will get you means and credible intervals returned in the $PopPref element
## the means are in the middle column, and the credible intervals are on either side
credibleIntervals(prefres = SimpPref[[1]],burn=10000,interval=0.95)  

#$IndPref
#, , 1
#
#             [,1]      [,2]      [,3]      [,4]
# [1,] 0.14985509 0.1181600 0.1964529 0.1885343
# [2,] 0.15839878 0.1163742 0.1290528 0.2065309
# [3,] 0.19495555 0.1150566 0.1704660 0.1493036
# [4,] 0.12409537 0.1435298 0.1683408 0.1720249
# [5,] 0.15785824 0.1752308 0.1243269 0.1890745
# [6,] 0.15063353 0.1378344 0.1568328 0.2103717
# [7,] 0.14635655 0.1625568 0.1933764 0.1448571
# [8,] 0.13858530 0.1234037 0.2058939 0.1779569
# [9,] 0.15630983 0.1540094 0.1543086 0.1582725
#[10,] 0.13772534 0.1777846 0.1665019 0.1479428
#[11,] 0.16053231 0.1445883 0.1218382 0.1931802
#[12,] 0.18339371 0.1792777 0.1701229 0.1248854
#[13,] 0.14652355 0.1546725 0.1655236 0.1684028
#[14,] 0.18238910 0.1503139 0.1365625 0.1285928
#[15,] 0.12676866 0.1330861 0.2031176 0.1863140
#[16,] 0.18919309 0.1746704 0.1344322 0.1465370
#[17,] 0.09816671 0.1131660 0.2166801 0.2185391
#
#, , 2
#
#           [,1]      [,2]      [,3]      [,4]
# [1,] 0.2336824 0.1941659 0.2862871 0.2785813
# [2,] 0.2539046 0.2034362 0.2203720 0.3117825
# [3,] 0.2929669 0.1959435 0.2624062 0.2392044
# [4,] 0.2131671 0.2372948 0.2671005 0.2722985
# [5,] 0.2441222 0.2630851 0.2038330 0.2799445
# [6,] 0.2325056 0.2172619 0.2410042 0.3012425
# [7,] 0.2294111 0.2488685 0.2844259 0.2290737
# [8,] 0.2203518 0.2035198 0.2996561 0.2677139
# [9,] 0.2497138 0.2447484 0.2457330 0.2495549
#[10,] 0.2246732 0.2712885 0.2583601 0.2365519
#[11,] 0.2538003 0.2348250 0.2070710 0.2942437
#[12,] 0.2693432 0.2656237 0.2542071 0.2031450
#[13,] 0.2344393 0.2424356 0.2552461 0.2596247
#[14,] 0.2853530 0.2481515 0.2330881 0.2227806
#[15,] 0.2052872 0.2133025 0.2964607 0.2767631
#[16,] 0.2806065 0.2635783 0.2159737 0.2315274
#[17,] 0.1725859 0.1914849 0.3120772 0.3155715
#
#, , 3
#
#           [,1]      [,2]      [,3]      [,4]
# [1,] 0.3325737 0.2872057 0.3907415 0.3802091
# [2,] 0.3680339 0.3132250 0.3326471 0.4332606
# [3,] 0.4052558 0.2979590 0.3723074 0.3476461
# [4,] 0.3251565 0.3522023 0.3852719 0.3890033
# [5,] 0.3471193 0.3668822 0.3030035 0.3854681
# [6,] 0.3318182 0.3131639 0.3395325 0.4057595
# [7,] 0.3293231 0.3496756 0.3881770 0.3296348
# [8,] 0.3214574 0.3011445 0.4072730 0.3718679
# [9,] 0.3617664 0.3545996 0.3574221 0.3613281
#[10,] 0.3287084 0.3787635 0.3658643 0.3438808
#[11,] 0.3649409 0.3440440 0.3135013 0.4103602
#[12,] 0.3697022 0.3634435 0.3532930 0.2963405
#[13,] 0.3374399 0.3470724 0.3619089 0.3661496
#[14,] 0.4064671 0.3669564 0.3499990 0.3373867
#[15,] 0.3051493 0.3108232 0.4014113 0.3810613
#[16,] 0.3866146 0.3665384 0.3152025 0.3328992
#[17,] 0.2676433 0.2873967 0.4201220 0.4249866
#
#
#$PopPref
#          [,1]      [,2]      [,3]
#[1,] 0.2001453 0.2429185 0.2897198
#[2,] 0.1909666 0.2337381 0.2784032
#[3,] 0.2144639 0.2574510 0.3048541
#[4,] 0.2174216 0.2647003 0.3107174
#
#$PopVar
#    2.5%      50%    97.5% 
#31.92268 45.41760 49.81595 


## It might also be handy to extract the pop level preference results, so they can be graphed
## or whatever.  Here I extract those two vectors for the two choices and find means manually
## which match the means found above:
## first extract the population preference element from the main result list
popPrefMCMC.S <- SimpPref[[1]]$PopPref
dim(popPrefMCMC.S)
# [1]     4 40000


## extract preferences for the three choices and find mean
prefCPIL.S <- popPrefMCMC.S[1,] 
mean(prefCPIL.S)
# [1] 0.2432266

prefCPCL.S <- popPrefMCMC.S[2,]
mean(prefCPCL.S)
# [1] 0.2347931

prefIPIL.S <- popPrefMCMC.S[3,]
mean(prefIPIL.S)
# [1] 0.2575888

prefIPCL.S <- popPrefMCMC.S[4,]
mean(prefIPCL.S)
# [1] 0.2643915

## pairwise probabilities
## in this case, you interpret the result such that
## you have 100% confidence (or almost) that preference for large (purple) is greater than preference for small (green),
## and that you have 0% confidence that preference for small is greater than preference for large.
pairwiseProb(SimpPref[[1]],burn=10000)

#          [,+-]      [,++]      [,--]      [,-+]
#[+-,] 0.0000000 0.5997000 0.3453667 0.2869333
#[++,] 0.4003000 0.0000000 0.2642000 0.2078333
#[--,] 0.6546333 0.7358000 0.0000000 0.4319667
#[-+,] 0.7130667 0.7921667 0.5680333 0.0000000


#### FOR SIMPLE TRAINED BEES - TEST ####
FILENAME12
simptest <- read.csv("~/LL stats/STATS_Counts_Simp_TestPhaseAnalysis_2-14-24_MC.csv", sep=",", header=TRUE,check.names=FALSE)
head(simptest)

## all bees in one group ('pop')
data2 <- subset(simptest,Pop==2,select=-c(BeeID))
data2 <- droplevels(data2)
## change pop label to 1, which is what bayesPref will expect for single analysis
data2$Pop <- rep(1,dim(data2)[1]) #called pop 1, but it's actually pop2. Just a quirk
data2 <- as.matrix(data2)
head(data2)

# run the model
SimpTest <-bayesPref(pData=data2,mcmcL=40000,pops=TRUE,dicburn=10000,dirvar=9)
#dic =  91.696 

# examine mixing, which is not great at 1000 reps
plot(SimpTest[[1]]$PopPref[1,],xlab="MCMC step",ylab="PopPref",cex=0.5) #some weird mixing at the burn-in part

## simple plot
prefPlot(SimpTest[[1]],ymax=35)

## means and credible intervals
## this next command will get you means and credible intervals returned in the $PopPref element
## the means are in the middle column, and the credible intervals are on either side
credibleIntervals(prefres = SimpTest[[1]],burn=10000,interval=0.95)  

#$IndPref
#, , 1
#
#          [,1]      [,2]       [,3]       [,4]
# [1,] 0.1840234 0.2038141 0.08572506 0.10124972
# [2,] 0.1460452 0.2103596 0.07802151 0.08067105
# [3,] 0.1328258 0.2286882 0.08821098 0.08807725
# [4,] 0.1821833 0.1861221 0.07702444 0.07872146
# [5,] 0.2010840 0.2041058 0.11297448 0.11415348
# [6,] 0.1551345 0.1584913 0.08372842 0.10306475
# [7,] 0.1491348 0.1737441 0.12917412 0.18325981
# [8,] 0.1933690 0.2274383 0.08958404 0.06201714
# [9,] 0.1266509 0.2231488 0.14327695 0.07578315
#[10,] 0.1918106 0.1964955 0.07512756 0.10501624
#[11,] 0.3078117 0.2024502 0.06772134 0.07786494
#[12,] 0.1918396 0.2120801 0.08927714 0.07598630
#[13,] 0.1562407 0.2424668 0.08074261 0.08088821
#[14,] 0.1483499 0.2310199 0.07861618 0.06268460
#[15,] 0.1979636 0.2388540 0.11277709 0.06926571
#[16,] 0.1623418 0.1859448 0.12022659 0.16460791
#[17,] 0.2493144 0.2165826 0.06950839 0.10924979
#
#, , 2
#
#           [,1]      [,2]      [,3]      [,4]
# [1,] 0.2990809 0.3220651 0.1733693 0.1927294
# [2,] 0.2746822 0.3496485 0.1787653 0.1812421
# [3,] 0.2513049 0.3621551 0.1847205 0.1861607
# [4,] 0.3128919 0.3194331 0.1748858 0.1766413
# [5,] 0.2995520 0.3038616 0.1931164 0.1941957
# [6,] 0.2869746 0.2949714 0.1875967 0.2132019
# [7,] 0.2378646 0.2674799 0.2112231 0.2743620
# [8,] 0.3097143 0.3529638 0.1795479 0.1448988
# [9,] 0.2335847 0.3450317 0.2474626 0.1598862
#[10,] 0.3095835 0.3163279 0.1613270 0.1996616
#[11,] 0.4149360 0.2969748 0.1328000 0.1460063
#[12,] 0.3099266 0.3340217 0.1792104 0.1638324
#[13,] 0.2697233 0.3727949 0.1710626 0.1725215
#[14,] 0.2742312 0.3725344 0.1793976 0.1573588
#[15,] 0.3012251 0.3485344 0.1979377 0.1425291
#[16,] 0.2533503 0.2836385 0.2008792 0.2529894
#[17,] 0.3505629 0.3169537 0.1362900 0.1872660
#
#, , 3
#
#           [,1]      [,2]      [,3]      [,4]
# [1,] 0.4325526 0.4562103 0.2930452 0.3152003
# [2,] 0.4266533 0.5119739 0.3200063 0.3210694
# [3,] 0.3941080 0.5147379 0.3190220 0.3218169
# [4,] 0.4662018 0.4737756 0.3138583 0.3144737
# [5,] 0.4122066 0.4170778 0.2960857 0.2944670
# [6,] 0.4447074 0.4549335 0.3347180 0.3617797
# [7,] 0.3424556 0.3744141 0.3118151 0.3842474
# [8,] 0.4462139 0.4932593 0.3020878 0.2592477
# [9,] 0.3636981 0.4838474 0.3813942 0.2761536
#[10,] 0.4471979 0.4539122 0.2810731 0.3236728
#[11,] 0.5298054 0.4066566 0.2216322 0.2367323
#[12,] 0.4497170 0.4722759 0.3024504 0.2817731
#[13,] 0.4084096 0.5201085 0.2957962 0.2977967
#[14,] 0.4260863 0.5349684 0.3189267 0.2938816
#[15,] 0.4192524 0.4683706 0.3074748 0.2416269
#[16,] 0.3613285 0.3937240 0.3016034 0.3595650
#[17,] 0.4647122 0.4272688 0.2262761 0.2845307
#
#
#$PopPref
#          [,1]      [,2]      [,3]
#[1,] 0.2397897 0.2950963 0.3564539
#[2,] 0.2662432 0.3285477 0.3909986
#[3,] 0.1403019 0.1864491 0.2414191
#[4,] 0.1452357 0.1876605 0.2418578
#
#$PopVar
#    2.5%      50%    97.5% 
#23.45906 40.03097 49.54358


## It might also be handy to extract the pop level preference results, so they can be graphed
## or whatever.  Here I extract those two vectors for the two choices and find means manually
## which match the means found above:
## first extract the population preference element from the main result list
popPrefTMCMC2.S <- SimpTest[[1]]$PopPref
dim(popPrefTMCMC2.S)
# [1]     4 40000

##FOR Experienced bees
prefCPILt.S <- popPrefTMCMC2.S[1,] 
mean(prefCPILt.S)
# [1] 0.3059817

prefCPCLt.S <- popPrefTMCMC2.S[2,]
mean(prefCPCLt.S)
# [1] [1] 0.3234709

prefIPILt.S <- popPrefTMCMC2.S[3,]
mean(prefIPILt.S)
# [1] [1] 0.186523

prefIPCLt.S <- popPrefTMCMC2.S[4,]
mean(prefIPCLt.S)
# [1] 0.1840244

## pairwise probabilities
## in this case, you interpret the result such that
## you have 100% confidence (or almost) that preference for large (purple) is greater than preference for small (green),
## and that you have 0% confidence that preference for small is greater than preference for large.
pairwiseProb(SimpTest[[1]],burn=10000)

#          [,+-]      [,++]      [,--]      [,-+]
#[+-,] 0.000000000 0.262466667 0.9930667 0.9870667
#[++,] 0.737533333 0.000000000 0.9911667 0.9969667
#[--,] 0.006933333 0.008833333 0.0000000 0.4702333
#[-+,] 0.012933333 0.003033333 0.5297667 0.0000000

#### COMPARISONS BETWEEN THE TWO PHASES FOR EACH TARGET FLOWER TYPE - Simple ####

pop1Full.S <- SimpPref[[1]]$PopPref[1,] # Pull out +- (correct color, incorrect leaf) for the Naive bees
pop1Chopped.S <- pop1Full.S[10001:40000] # saves the steps after the burn in
#then for the other population
pop2Full.S <- SimpTest[[1]]$PopPref[1,] # Pull out +- (correct color, incorrect leaf) for the Experienced bees
pop2Chopped.S <- pop2Full.S[10001:40000] #Post burn in
dif <- pop1Chopped.S - pop2Chopped.S #How many times in the posterior 1 is bigger than the other
CPILprob.s <- length(dif[dif>0]) / 30000 # should be the amount post burn in throw out
CPILprob.s
# [1] 0.08373333

pop1Full.S <- SimpPref[[1]]$PopPref[2,] # Pull out ++ (correct combo) for the Naive bees
pop1Chopped.S <- pop1Full.S[10001:40000] # saves the steps after the burn in
#then for the other population
pop2Full.S <- SimpTest[[1]]$PopPref[2,] # Pull out ++ (correct combo) for the Experienced bees
pop2Chopped.S <- pop2Full.S[10001:40000] #Post burn in
dif <- pop1Chopped.S - pop2Chopped.S #How many times in the posterior 1 is bigger than the other
CPCLprob.s <- length(dif[dif>0]) / 30000 # should be the amount post burn in throw out
CPCLprob.s
# [1] 0.0074

pop1Full.S <- SimpPref[[1]]$PopPref[3,] # Pull out -- (incorrect color, incorrect leaf) for the Naive bees
pop1Chopped.S <- pop1Full.S[10001:40000] # saves the steps after the burn in
#then for the other population
pop2Full.S <- SimpTest[[1]]$PopPref[3,] # Pull out -- (incorrect color, incorrect leaf) for the Experienced bees
pop2Chopped.S <- pop2Full.S[10001:40000] #Post burn in
dif <- pop1Chopped.S - pop2Chopped.S #How many times in the posterior 1 is bigger than the other
IPILprob.s <- length(dif[dif>0]) / 30000 # should be the amount post burn in throw out
IPILprob.s
# [1] 0.9776667

pop1Full.S <- SimpPref[[1]]$PopPref[4,] # Pull out -+ (incorrect color, correct leaf) for the Naive bees
pop1Chopped.S <- pop1Full.S[10001:40000] # saves the steps after the burn in
#then for the other population
pop2Full.S <- SimpTest[[1]]$PopPref[4,] # Pull out -+ (incorrect color, correct leaf) for the Experienced bees
pop2Chopped.S <- pop2Full.S[10001:40000] #Post burn in
dif <- pop1Chopped.S - pop2Chopped.S #How many times in the posterior 1 is bigger than the other
IPCLprob.s <- length(dif[dif>0]) / 30000 # should be the amount post burn in throw out
IPCLprob.s
# [1] 0.9796333


#### FOR DIFFICULT TRAINED BEES - INITIAL ####
FILENAME9 
diffpref <- read.csv("~/LL stats/STATS_Counts_Diff_InitialPhaseAnalysis_2-14-24_MC.csv", sep=",", header=TRUE,check.names=FALSE) 
head(diffpref)

## all bees in one group ('pop')
data3 <- subset(simppref,Pop==1,select=-c(BeeID))
data3 <- droplevels(data3)
data3 <- as.matrix(data3)
head(data3)

# run the model
DiffPref <- bayesPref(pData=data3,mcmcL=40000,pops=TRUE,dicburn=10000,dirvar=9)
#dic = 116.3186 

# examine mixing, which is not great at 1000 reps
plot(DiffPref[[1]]$PopPref[1,],xlab="MCMC step",ylab="PopPref",cex=0.5) #looks good

## simple plot
prefPlot(DiffPref[[1]],ymax=35)

## means and credible intervals
## this next command will get you means and credible intervals returned in the $PopPref element
## the means are in the middle column, and the credible intervals are on either side
credibleIntervals(prefres = DiffPref[[1]],burn=10000,interval=0.95)  

#$IndPref
#, , 1
#
#          [,1]      [,2]      [,3]      [,4]
# [1,] 0.1496171 0.1178737 0.1959039 0.1885875
# [2,] 0.1563806 0.1155076 0.1269299 0.2059881
# [3,] 0.1951137 0.1144186 0.1689651 0.1505138
# [4,] 0.1228339 0.1430240 0.1675794 0.1712116
# [5,] 0.1564025 0.1754402 0.1231383 0.1893114
# [6,] 0.1489897 0.1362879 0.1551556 0.2087163
# [7,] 0.1450999 0.1632273 0.1923741 0.1453623
# [8,] 0.1373358 0.1239761 0.2067177 0.1774477
# [9,] 0.1566147 0.1534736 0.1518883 0.1567727
#[10,] 0.1374591 0.1783593 0.1640393 0.1471667
#[11,] 0.1590343 0.1447488 0.1199097 0.1952777
#[12,] 0.1822518 0.1801398 0.1687095 0.1256966
#[13,] 0.1460642 0.1551963 0.1635697 0.1665256
#[14,] 0.1808060 0.1509956 0.1371214 0.1262274
#[15,] 0.1248007 0.1333206 0.2032427 0.1860385
#[16,] 0.1879169 0.1753801 0.1330488 0.1458585
#[17,] 0.0955864 0.1142274 0.2168690 0.2200325
#
#, , 2
#
#           [,1]      [,2]      [,3]      [,4]
# [1,] 0.2329518 0.1953437 0.2856224 0.2781628
# [2,] 0.2538642 0.2051533 0.2186233 0.3127611
# [3,] 0.2929925 0.1971807 0.2614934 0.2394615
# [4,] 0.2128439 0.2378310 0.2669732 0.2721564
# [5,] 0.2442699 0.2653625 0.2026459 0.2791010
# [6,] 0.2333528 0.2181685 0.2395580 0.3013394
# [7,] 0.2296066 0.2500255 0.2846223 0.2275660
# [8,] 0.2201696 0.2043799 0.2995779 0.2671213
# [9,] 0.2502383 0.2464742 0.2450897 0.2494428
#[10,] 0.2242861 0.2726613 0.2581103 0.2361162
#[11,] 0.2540525 0.2354909 0.2054868 0.2951700
#[12,] 0.2701283 0.2672503 0.2538954 0.2015901
#[13,] 0.2335498 0.2433537 0.2552285 0.2591589
#[14,] 0.2854249 0.2499762 0.2324090 0.2211671
#[15,] 0.2054389 0.2139331 0.2958525 0.2761909
#[16,] 0.2808694 0.2649705 0.2150789 0.2307024
#[17,] 0.1708294 0.1922504 0.3124148 0.3157870
#
#, , 3
#
#           [,1]      [,2]      [,3]      [,4]
# [1,] 0.3318141 0.2892443 0.3899432 0.3795675
# [2,] 0.3708731 0.3156273 0.3301163 0.4337903
# [3,] 0.4053256 0.3000857 0.3696544 0.3464469
# [4,] 0.3255355 0.3536816 0.3848871 0.3917708
# [5,] 0.3468215 0.3706421 0.3030818 0.3840984
# [6,] 0.3330094 0.3145054 0.3391668 0.4065384
# [7,] 0.3295338 0.3531647 0.3902800 0.3293265
# [8,] 0.3208877 0.3026680 0.4067669 0.3707691
# [9,] 0.3634018 0.3589225 0.3547540 0.3589344
#[10,] 0.3304632 0.3815121 0.3666652 0.3431792
#[11,] 0.3673449 0.3447670 0.3139368 0.4110365
#[12,] 0.3696034 0.3675276 0.3535612 0.2946936
#[13,] 0.3404354 0.3490030 0.3631342 0.3683666
#[14,] 0.4121998 0.3689934 0.3501590 0.3372153
#[15,] 0.3036680 0.3129046 0.4031462 0.3814912
#[16,] 0.3874604 0.3697463 0.3154319 0.3330410
#[17,] 0.2649016 0.2883957 0.4211321 0.4244776
#
#
#$PopPref
#          [,1]      [,2]      [,3]
#[1,] 0.1978347 0.2427299 0.2905741
#[2,] 0.1928286 0.2354465 0.2834153
#[3,] 0.2115875 0.2557924 0.3029814
#[4,] 0.2204808 0.2638021 0.3132048
#
#$PopVar
#    2.5%      50%    97.5% 
#28.81294 44.77552 49.80584 


## It might also be handy to extract the pop level preference results, so they can be graphed
## or whatever.  Here I extract those two vectors for the two choices and find means manually
## which match the means found above:
## first extract the population preference element from the main result list
popPrefMCMC.D <- DiffPref[[1]]$PopPref
dim(popPrefMCMC.D)
# [1]     4 40000


## extract preferences for the three choices and find mean
prefCPIL.D <- popPrefMCMC.D[1,] 
mean(prefCPIL.D)
# [1] 0.2437058

prefCPCL.D <- popPrefMCMC.D[2,]
mean(prefCPCL.D)
# [1] 0.2344112

prefIPIL.D <- popPrefMCMC.D[3,]
mean(prefIPIL.D)
# [1] 0.2571238 

prefIPCL.D <- popPrefMCMC.D[4,]
mean(prefIPCL.D)
# [1] 0.2647592

## pairwise probabilities
## in this case, you interpret the result such that
## you have 100% confidence (or almost) that preference for large (purple) is greater than preference for small (green),
## and that you have 0% confidence that preference for small is greater than preference for large.
pairwiseProb(DiffPref[[1]],burn=10000)

#       [,+-]      [,++]      [,--]      [,-+]
#[+-,] 0.0000000 0.5756667 0.3528333 0.2920667
#[++,] 0.4243333 0.0000000 0.2986667 0.2323667
#[--,] 0.6471667 0.7013333 0.0000000 0.4224667
#[-+,] 0.7079333 0.7676333 0.5775333 0.0000000


#### FOR DIFFICULT TRAINED BEES - TEST ####
FILENAME12
difftest <- read.csv("~/LL stats/STATS_Counts_Diff_TestPhaseAnalysis_2-14-24_MC.csv", sep=",", header=TRUE,check.names=FALSE)
head(difftest)

## all bees in one group ('pop')
data4 <- subset(difftest,Pop==2,select=-c(BeeID))
data4 <- droplevels(data4)
## change pop label to 1, which is what bayesPref will expect for single analysis
data4$Pop <- rep(1,dim(data4)[1]) #called pop 1, but it's actually pop2. Just a quirk
data4 <- as.matrix(data4)
head(data4)

# run the model
DiffTest <-bayesPref(pData=data4,mcmcL=40000,pops=TRUE,dicburn=10000,dirvar=9)
#dic =  105.0792 

# examine mixing, which is not great at 1000 reps
plot(DiffTest[[1]]$PopPref[1,],xlab="MCMC step",ylab="PopPref",cex=0.5) #mixing good

## simple plot
prefPlot(DiffTest[[1]],ymax=35)

## means and credible intervals
## this next command will get you means and credible intervals returned in the $PopPref element
## the means are in the middle column, and the credible intervals are on either side
credibleIntervals(prefres = DiffTest[[1]],burn=10000,interval=0.95)  

#$IndPref
#, , 1
#
#           [,1]      [,2]       [,3]      [,4]
# [1,] 0.1570443 0.1638046 0.10927373 0.1392736
# [2,] 0.1963140 0.1988998 0.11385192 0.1432117
# [3,] 0.1711571 0.1857828 0.13797852 0.1488117
# [4,] 0.1771377 0.1930935 0.15272754 0.1079262
# [5,] 0.1747626 0.1680117 0.10701680 0.1820998
# [6,] 0.2015020 0.1274823 0.14592770 0.1211150
# [7,] 0.1933389 0.1979047 0.13239432 0.1317145
# [8,] 0.1632776 0.1669610 0.11648855 0.1305103
# [9,] 0.2162368 0.1164571 0.13044103 0.1820127
#[10,] 0.1625528 0.1883288 0.11600149 0.1690599
#[11,] 0.1737709 0.1651136 0.10531755 0.1557286
#[12,] 0.1919887 0.1272503 0.09798875 0.1163784
#[13,] 0.1823164 0.1478368 0.15143161 0.1703199
#[14,] 0.1438499 0.1818018 0.09736971 0.1413783
#[15,] 0.1484390 0.1232249 0.12902757 0.1601486
#[16,] 0.1779891 0.1441648 0.13583064 0.1483043
#[17,] 0.1471097 0.1685265 0.20179263 0.1445907
#[18,] 0.1744788 0.1679874 0.13922612 0.1604084
#[19,] 0.1478945 0.1531646 0.11273364 0.1439346
#[20,] 0.1704804 0.1559894 0.14932496 0.1701199
#
#, , 2
#
#           [,1]      [,2]      [,3]      [,4]
# [1,] 0.2653840 0.2730427 0.2058984 0.2428765
# [2,] 0.2866955 0.2910159 0.1899253 0.2243089
# [3,] 0.2612942 0.2783110 0.2204154 0.2328000
# [4,] 0.2715337 0.2896069 0.2405789 0.1889840
# [5,] 0.2677586 0.2610070 0.1860526 0.2768650
# [6,] 0.3113083 0.2225705 0.2424007 0.2121263
# [7,] 0.2824442 0.2877228 0.2108075 0.2110190
# [8,] 0.2703570 0.2758010 0.2120701 0.2304207
# [9,] 0.3125381 0.1953161 0.2111532 0.2725983
#[10,] 0.2518850 0.2825627 0.1964931 0.2605078
#[11,] 0.2747711 0.2664255 0.1928347 0.2550749
#[12,] 0.3170169 0.2397679 0.2020869 0.2247265
#[13,] 0.2708772 0.2308935 0.2329703 0.2568837
#[14,] 0.2526463 0.2968837 0.1907627 0.2471036
#[15,] 0.2573053 0.2267011 0.2317746 0.2704519
#[16,] 0.2782166 0.2393092 0.2286627 0.2435527
#[17,] 0.2270147 0.2526229 0.2889175 0.2238538
#[18,] 0.2640576 0.2561963 0.2234102 0.2482088
#[19,] 0.2572016 0.2641465 0.2130373 0.2515192
#[20,] 0.2605427 0.2407731 0.2328123 0.2572231
#
#, , 3
#
#           [,1]      [,2]      [,3]      [,4]
# [1,] 0.3961993 0.4042494 0.3299855 0.3727460
# [2,] 0.3907606 0.3943722 0.2831573 0.3233402
# [3,] 0.3645417 0.3839791 0.3177964 0.3336581
# [4,] 0.3827986 0.4027691 0.3481935 0.2893293
# [5,] 0.3777335 0.3692574 0.2840537 0.3856826
# [6,] 0.4366030 0.3405800 0.3617324 0.3275424
# [7,] 0.3852539 0.3917227 0.3067749 0.3080953
# [8,] 0.3974656 0.4034091 0.3320684 0.3540899
# [9,] 0.4206098 0.2929705 0.3102545 0.3784684
#[10,] 0.3588992 0.3929577 0.2973522 0.3677245
#[11,] 0.3965509 0.3862520 0.3034848 0.3751860
#[12,] 0.4677051 0.3812189 0.3402693 0.3663780
#[13,] 0.3733065 0.3292377 0.3312041 0.3583082
#[14,] 0.3824537 0.4341851 0.3136230 0.3779733
#[15,] 0.3920555 0.3552570 0.3621920 0.4045242
#[16,] 0.3954207 0.3524077 0.3407577 0.3589150
#[17,] 0.3220501 0.3512419 0.3890687 0.3192565
#[18,] 0.3686577 0.3594171 0.3240202 0.3516917
#[19,] 0.3915426 0.3987202 0.3421065 0.3833639
#[20,] 0.3652667 0.3442317 0.3340797 0.3607247
#
#
#$PopPref
#          [,1]      [,2]      [,3]
#[1,] 0.2271570 0.2738874 0.3229596
#[2,] 0.2137936 0.2601857 0.3042664
#[3,] 0.1769893 0.2207240 0.2639553
#[4,] 0.2014344 0.2446150 0.2904214
#
#$PopVar
#    2.5%      50%    97.5% 
#33.22634 45.75335 49.82945 


## It might also be handy to extract the pop level preference results, so they can be graphed
## or whatever.  Here I extract those two vectors for the two choices and find means manually
## which match the means found above:
## first extract the population preference element from the main result list
popPrefTMCMC2.D <- DiffTest[[1]]$PopPref
dim(popPrefTMCMC2.D)
# [1]     4 40000

##FOR Experienced bees
prefCPILt.D <- popPrefTMCMC2.D[1,] 
mean(prefCPILt.D)
# [1] 0.2740753

prefCPCLt.D <- popPrefTMCMC2.D[2,]
mean(prefCPCLt.D)
# [1] 0.2602124 

prefIPILt.D <- popPrefTMCMC2.D[3,]
mean(prefIPILt.D)
# [1] 0.2204296

prefIPCLt.D <- popPrefTMCMC2.D[4,]
mean(prefIPCLt.D)
# [1] 0.2452827

## pairwise probabilities
## in this case, you interpret the result such that
## you have 100% confidence (or almost) that preference for large (purple) is greater than preference for small (green),
## and that you have 0% confidence that preference for small is greater than preference for large.
pairwiseProb(DiffTest[[1]],burn=10000)

#      [,+-]      [,++]      [,--]      [,-+]
#[+-,] 0.00000000 0.6365000 0.9212333 0.7858667
#[++,] 0.36350000 0.0000000 0.8502667 0.6621000
#[--,] 0.07876667 0.1497333 0.0000000 0.2568333
#[-+,] 0.21413333 0.3379000 0.7431667 0.0000000


#### COMPARISONS BETWEEN THE TWO PHASES FOR EACH TARGET FLOWER TYPE - Difficult ####

pop1Full.D <- DiffPref[[1]]$PopPref[1,] # Pull out +- (correct color, incorrect leaf) for the Naive bees
pop1Chopped.D <- pop1Full.D[10001:40000] # saves the steps after the burn in
#then for the other population
pop2Full.D <- DiffTest[[1]]$PopPref[1,] # Pull out +- (correct color, incorrect leaf) for the Experienced bees
pop2Chopped.D <- pop2Full.D[10001:40000] #Post burn in
dif <- pop1Chopped.D - pop2Chopped.D #How many times in the posterior 1 is bigger than the other
CPILprob.d <- length(dif[dif>0]) / 30000 # should be the amount post burn in throw out
CPILprob.d
# [1] 0.1807333

pop1Full.D <- DiffPref[[1]]$PopPref[2,] # Pull out ++ (correct combo) for the Naive bees
pop1Chopped.D <- pop1Full.D[10001:40000] # saves the steps after the burn in
#then for the other population
pop2Full.D <- DiffTest[[1]]$PopPref[2,] # Pull out ++ (correct combo) for the Experienced bees
pop2Chopped.D <- pop2Full.D[10001:40000] #Post burn in
dif <- pop1Chopped.D - pop2Chopped.D #How many times in the posterior 1 is bigger than the other
CPCLprob.d <- length(dif[dif>0]) / 30000 # should be the amount post burn in throw out
CPCLprob.d
# [1] 0.2268

pop1Full.D <- DiffPref[[1]]$PopPref[3,] # Pull out -- (incorrect color, incorrect leaf) for the Naive bees
pop1Chopped.D <- pop1Full.D[10001:40000] # saves the steps after the burn in
#then for the other population
pop2Full.D <- DiffTest[[1]]$PopPref[3,] # Pull out -- (incorrect color, incorrect leaf) for the Experienced bees
pop2Chopped.D <- pop2Full.D[10001:40000] #Post burn in
dif <- pop1Chopped.D - pop2Chopped.D #How many times in the posterior 1 is bigger than the other
IPILprob.d <- length(dif[dif>0]) / 30000 # should be the amount post burn in throw out
IPILprob.d
# [1] 0.8682667

pop1Full.D <- DiffPref[[1]]$PopPref[4,] # Pull out -+ (incorrect color, correct leaf) for the Naive bees
pop1Chopped.D <- pop1Full.D[10001:40000] # saves the steps after the burn in
#then for the other population
pop2Full.D <- DiffTest[[1]]$PopPref[4,] # Pull out -+ (incorrect color, correct leaf) for the Experienced bees
pop2Chopped.D <- pop2Full.D[10001:40000] #Post burn in
dif <- pop1Chopped.D - pop2Chopped.D #How many times in the posterior 1 is bigger than the other
IPCLprob.d <- length(dif[dif>0]) / 30000 # should be the amount post burn in throw out
IPCLprob.d
# [1] 0.7234 


```
````
